# Covalent targeting leads to the development of LIMK1 isoform-selective inhibitors

**DOI:** 10.1101/2025.04.17.649341

**Authors:** Sebastian Mandel, Thomas Hanke, Niall Prendiville, María Baena-Nuevo, Lena Berger, Frederic Farges, Martin-Peter Schwalm, Benedict-Tilmann Berger, Andreas Kraemer, Lewis Elson, Hayuningbudi Saraswati, Kamal Rayees Abdul Azeez, Verena Dederer, Sebastian Mathea, Ana Corrionero, Patricia Alfonso, Sabrina Keller, Matthias Gstaiger, Daniela S. Krause, Susanne Müller, Sandra Röhm, Stefan Knapp

## Abstract

Selectivity for closely related isoforms of protein kinases is a major challenge in the design of drugs and chemical probes. Covalent targeting of unique cysteines is a potential strategy to achieve selectivity for highly conserved binding sites. Here, we used a pan-LIMK inhibitor to selectively probe LIMK1 over LIMK2 by targeting the LIMK1-specific cysteine C349 located in the glycine-rich loop region. Binding kinetics of both non-covalent and covalent LIMK inhibitors were investigated, and the fast on-rate and small size of type-I inhibitors were used in the design of a covalent LIMK1 inhibitor. The developed cell-active, isoform-selective LIMK1 inhibitor showed excellent proteome-wide selectivity in pull-down assays, enabling studies of LIMK1 isoform-selective functions in cellular model systems and providing a versatile chemical tool for studies of the LIMK signalling pathway.

## Introduction

The LIM kinase (LIMK) family comprises two isoforms in humans, LIMK1 and LIMK2, which exhibit dual serine/threonine and tyrosine kinase activity.^1,2^ Both enzymes share a high sequence identity of approximately 70% within their kinase domains.^3^ LIMKs play a crucial role in regulating actin cytoskeletal dynamics, including G-actin and stress fiber formation.^4^ The LIMK signaling pathway is primarily activated by small GTPases of the Rho family, such as RhoA, Rac, and CDC42, which stimulate the downstream kinases PAKs, MRCKa, or ROCK1/2, to activate LIMK1 and LIMK2. Activated LIMKs typically phosphorylate the actin-binding proteins cofilin1, cofilin2, and destrin, which lose their ability to cause actin depolymerization.^5–8^ As a result of cofilin inactivity, actin polymerizes into stable F-actin and forms stress fibers that affect various cellular functions such as motility, differentiation, spreading, and apoptosis.^9^ The invasive phenotype has been implicated in the oncogenesis of colon cancer, breast cancer,^10,11^ and glaucoma.^12^ In addition, LIM kinases affect synaptic plasticity, dendritic spine formation, and cellular mechanisms such as long-term potentiation (LTP), which is essential for learning and memory function in the brain.^13^ Following LTP induction, the LIMK1-cofilin pathway affects the trafficking and accumulation of AMPA receptors at the synapse.^14^ Consequently, alterations predominantly in LIMK1 signaling have been implicated in several neurological disorders, including Alzheimer’s disease, Parkinson’s disease, and Williams-Beuren syndrome.^5^ Interestingly, in Fragile X syndrome (FXS), increased synthesis of bone morphogenetic protein (BMP) type 2 receptor (BMPR2) results in LIMK1 activation, and pharmacological inhibition of the BMPR2-LIMK pathway ameliorates the synaptic abnormalities and locomotor phenotypes, suggesting LIMK1 selective inhibitors as a therapy for FXS.^15,16^

However, isoform-selective targeting within highly conserved kinase families is a formidable challenge. In some kinase families, cysteines are present in only one isoform, providing a strategy for isoform-selective targeting by covalent inhibitors.^17^ For instance, excellent isoform selectivity has been achieved by covalent inhibitors selectively targeting JAK3 within the JAK kinase family (JAK1-3 and TYK2)^18–20^ as well as targeting the Ephrin receptor isoform EphB3.^21^ Since the kinetics and target engagement of covalent inhibitors are different compared to their non-covalent counterparts, a deeper understanding of different mechanisms is inevitable. Several inhibitors have been developed for the LIM kinase family, including inhibitors with type-I, type-II, and type-III binding modes (Figure 1). However, many of these inhibitors have a fairly poor selectivity profile or contain functional groups in their chemical structure that should be considered critically, especially LX7101, Pyr1, and Damnacanthal which should not be used as chemical probes for LIMKs.^22^

**Figure 1.**
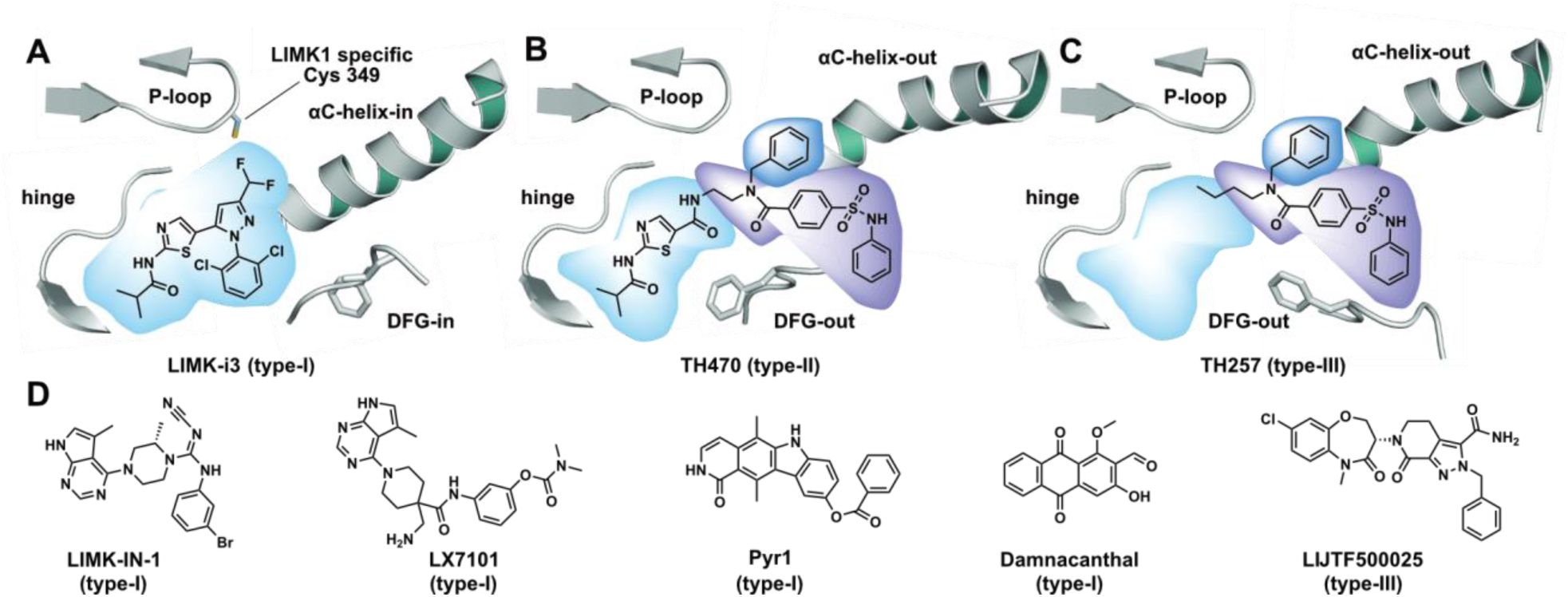
Binding mode of dual LIM kinase inhibitors. (A) Type-I binding mode: LIMK-i3 targeting the ATP pocket with active DFG-in conformation in LIMK1 (PDB: 8AAU). (B) Type-II binding mode: TH470 targeting the front and back-pocket area of LIMK2 in the inactive DFG-out conformation. The benzyl moiety engaged with the glycine-rich loop pocket created by the movement of the αC-helix toward an outward position (PDB: 7QHG). (C) Type-III binding mode: TH257 binding in the allosteric DFG-out and P-loop pocket of LIMK2 (PDB: 5NXD). (D) Examples of published LIMK1/2 inhibitors and their proposed binding modes.

LIMK-i3, developed by Bristol-Myers Squibb in 2006, was the first inhibitor reported to potent and selectively inhibit LIMK1/2-dependent phosphorylation of cofilin.^4,23^ Animal studies in mice conditioned with a contextual fear paradigm resulted in impaired memory reconsolidation after contextual re-exposure, highlighting the role of LIM kinases in neuronal function.^24^ LIMK-IN-I has been studied in the context of open-angle glaucoma, a progressive neurodegenerative disease of the inner retina that also damages the optic nerve.^4,25^ Very potent chemical probes were developed by the Structural Genomics Consortium alone, TH257, and in collaboration with Takeda, LIJTF500025.^22,26^ Both type-III inhibitors bind to LIMK1/2 in an allosteric pocket and showed very high selectivity against a large panel of kinases. The scaffolds of LIMK-i3 and TH257 were later used to design the type-II inhibitor TH470 by fusing both the ATP and DFG-out pocket moieties.^22^ Furthermore, TH257 served as a lead for the development of MDI-114215, which was used for studying cofilin phosphorylation in iPSC neurons derived from FXS patients.^27^ Due to the high sequence identity of both LIMK1/2 kinase domains, none of these inhibitors exhibit isoform selectivity for LIMK1 or LIMK2. Sequence analysis revealed that LIMK1 contains a reactive cysteine in the glycine-rich P-loop region. This cysteine was previously identified to be covalently modified by the promiscuous kinase inhibitor SM1-71 and may be a promising starting point for developing LIMK1 selective inhibitors.^28^ In this work, we report the first selective LIMK1 inhibitor based on the well-characterized non-covalent dual LIMK1/2 inhibitor LIMK-i3 by targeting an isoform-specific cysteine in the P-loop of the kinase domain. The presented inhibitor exhibits an excellent kinome-wide selectivity profile and submicromolar cellular on-target potency. The activity against the close isoform LIMK2 is >30-fold in cells, making SM311 (**10**) a valuable chemical probe that selectively targets LIMK1.

### Chemistry

All compounds were synthesized by a seven-step synthetic route in slight modification to the previous publication by Ross-McDonald et al. (Scheme 1).^23^ In the first step, 2,4-dimethoxybenzylamine (**1**) was reacted with thiocarbonyldiimidazole in a nucleophilic substitution. Subsequent aminolysis led to the thiourea derivative **2**. The compound was then reacted with dimethylformamide dimethylacetal to form the imine derivative **3**, which was then reacted with chloroacetate in a Hantzsch-like reaction to build the thiazole heterocycle. Afterward, **4** was treated with sodium ethoxide and diethyldifluoromalonate to obtain **5**. In the fifth step of the reaction, the pyrazoles, **6a-m,** were formed by a Knorr-pyrazole synthesis using the corresponding phenylhydrazine derivatives **7a-m**. After deprotection of compounds **6a-m** with TFA, the aminothiazole derivatives **8a-m** were alkylated with isobutyryl chloride to obtain the final compounds **9a-m**.

**Scheme 1.**
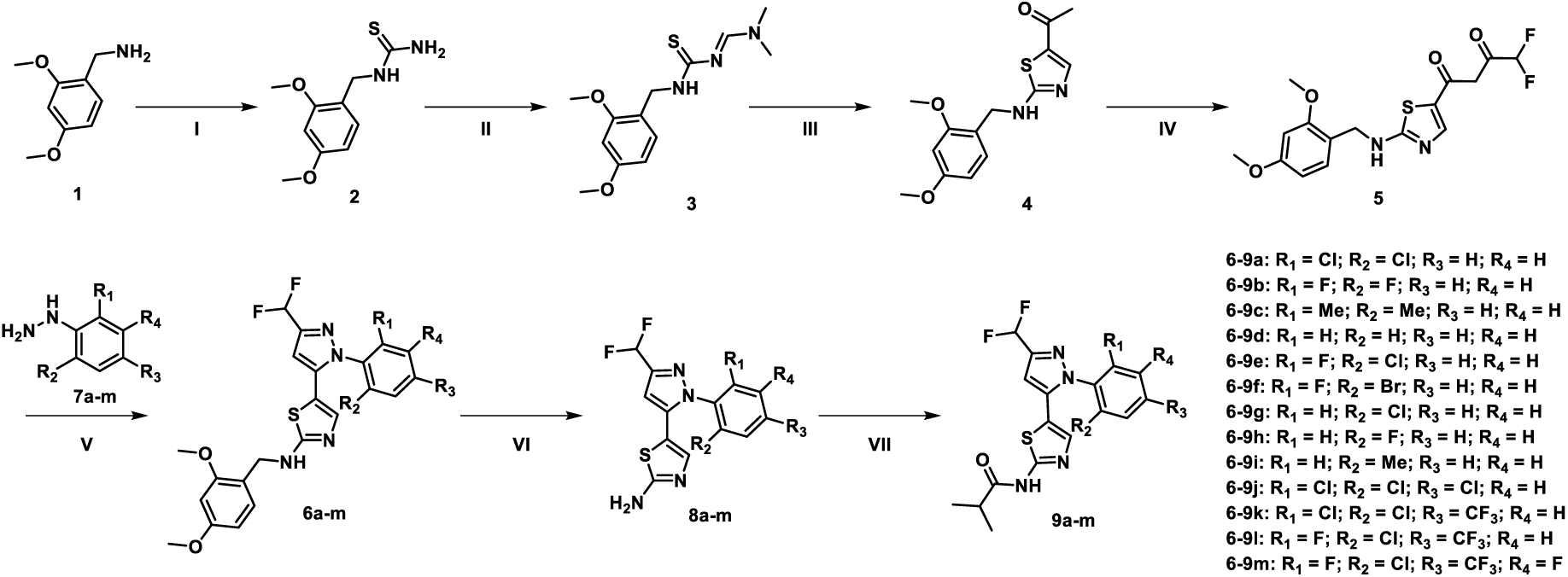
Synthetic route for the synthesis of LIMK1 type-I inhibitors **9a-m.**^a^ ^a^(Ia) TCDI, DCM, 3 h, RT; (Ib) NH_3_/MeOH (7M), 72 h, RT; (II) DMF-DMA, EtOH, 1 h, 80 °C; (III) chloroacetone, ACN, 3 h, 75°C; (IV) diethyl 2,2-difluoromalonate, NaOEt/EtOH, 72 h, 75 °C; (V) phenylhydrazine (**7a-m**), EtOH, 2 h, 75°C; (VI) TFA, H_2_O, 4 h, RT; (VII) isobutyryl chloride, DCM, pyridine, 20 h, RT. For final compounds, see Table 1.

For the synthesis of the covalent tool compound SM311 (**10**), the intermediate **5** was reacted with (4-nitrophenyl) hydrazine to build the pyrazole heterocycle **11** (Scheme 2). After deprotection of the amine with TFA, the aminothiazole **12** was treated with isobutyryl chloride to obtain **13**. The nitro group was reduced with iron to form compound **14**. Finally, the acrylamide warhead was installed by the reaction with acryloyl chloride to yield the title compound **10**.

**Scheme 2.**
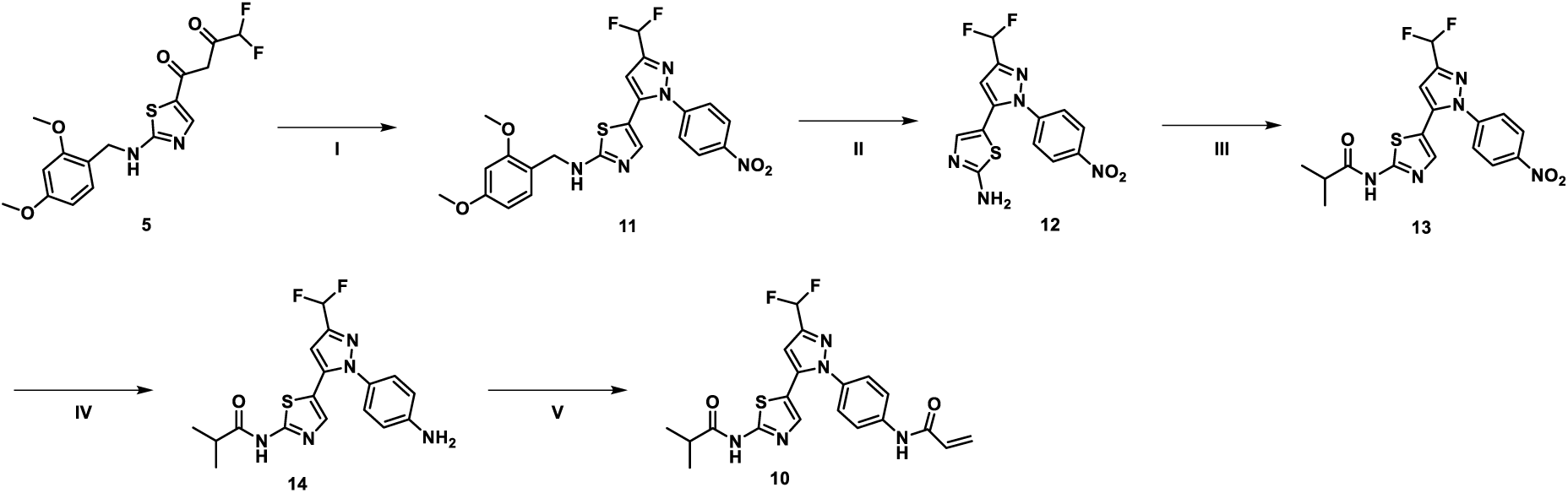
Synthetic route for synthesizing the covalent compound SM311 (**10**).^b^ ^b^(I) (4-Nitrophenyl) hydrazine, EtOH, 2 h, 75°C; (II) TFA/H_2_O, 4 h, RT; (III) isobutyryl chloride, ACN, DIEA, 20 h, RT; (IV) Fe, NH_4_Cl, MeOH/H_2_O, 20 h, 80°C; (V) acryloyl chloride, ACN, DIEA, 20 h, RT.

The synthesis of the biotin derivative **15** and the negative control **16** followed a similar synthetic route.(Scheme 3, Scheme 4). For the biotin derivative an aliphatic linker was attached to the aminothiazole **13**. After deprotection with TFA, biotin was introduced via an NHS ester. Reduction of the nitro group led to compound **24**, that was further decorated with an acrylamide warhead.

**Scheme 3.**
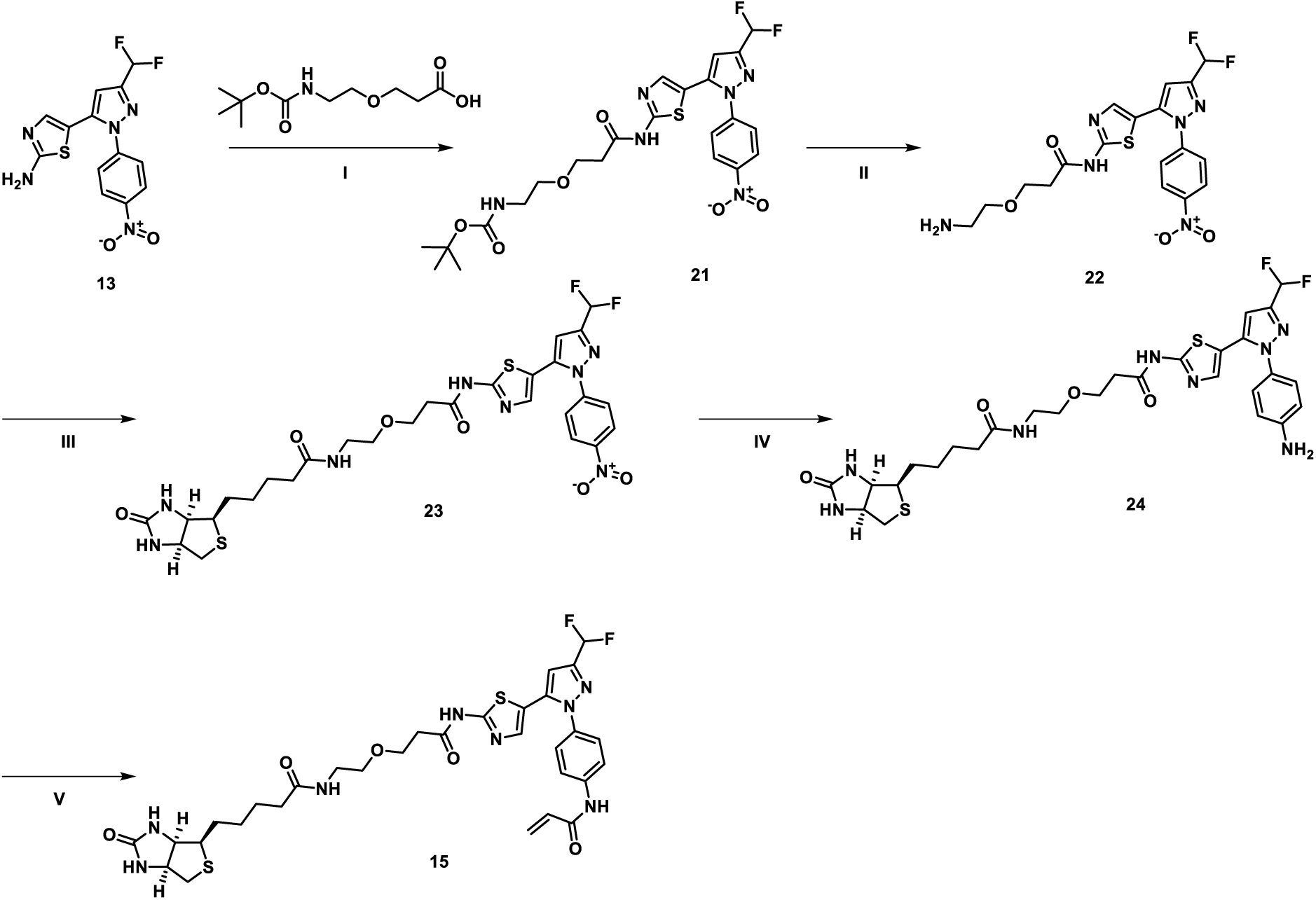
Synthetic route for synthesizing the biotin adduct 15^b^. ^b^(I) HATU, DIEA, DMF, 16 h, RT; (II) TFA, DCM, 1 h, RT; (III) Biotin-NHS, DIEA, 1h, RT; (IV) Zn, AcOH, MeOH, 3 h, RT; (V) acryloyl chloride, ACN, DIEA, 20 h, RT.

For the negative control, **16**, the aminothiazole derivative **12** was methylated. After deprotection, **18** was treated with isobutyryl chloride to obtain **19.** Reduction of the nitro group led to compound **20**, which was further decorated with an acrylamide warhead.

**Scheme 4.**
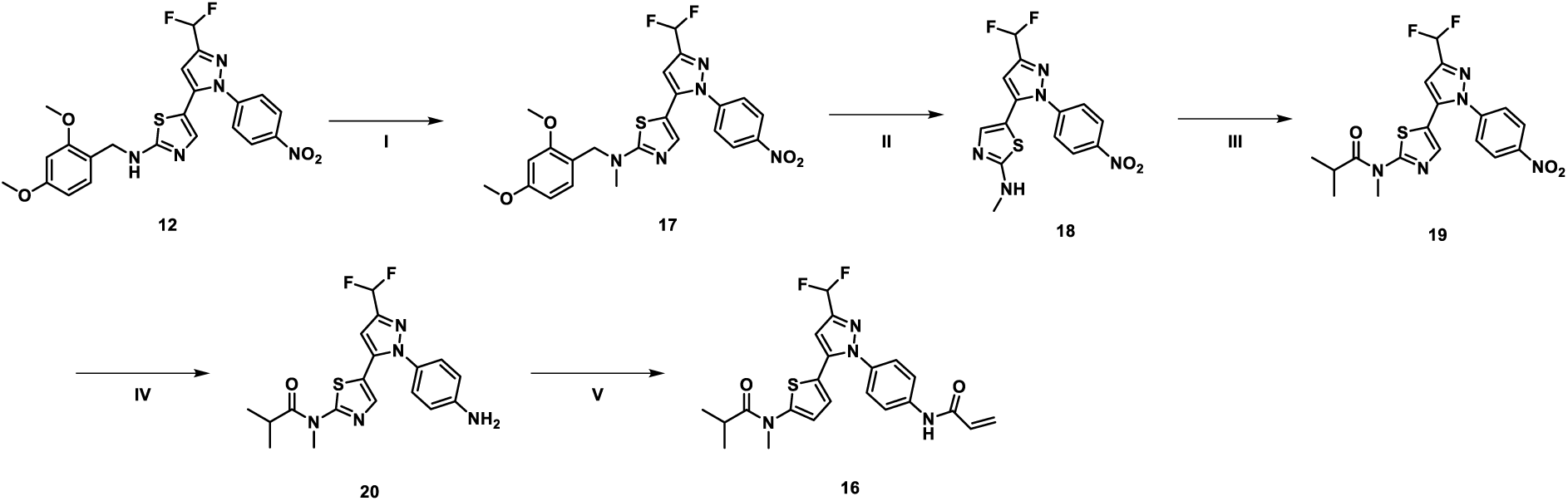
Synthetic route for synthesizing the LIMK1 negative control compound 16^a^. ^a^(Ia) NaH (60% in mineral oil), DMF, 10 min, 0 °C; (Ib) MeI (2M in THF), 16 h, RT; (II) TFA/H_2_O, 4 h, RT; (III) isobutyryl chloride, ACN, DIEA, 20 h, RT; (IV) Fe, NH_4_Cl, MeOH/H_2_O, 20 h, 80°C; (V) acryloyl chloride, ACN, DIEA, 20 h, RT.

## Results and Discussion

### Kinetic studies of LIMK inhibitors

Covalent inhibitors are characterized by long target residence times. Two types of covalent inhibitors can be distinguished: irreversible inhibitors, which permanently block the target binding pocket once the covalent bond is formed, and reversible-covalent inhibitors, such as cyanoacrylamides, which bind to the target for a prolonged period of time depending on the stability of the covalent bond formed. Reversible inhibitors, on the other hand, may have slow on- and off-rates due to conformational changes required for binding or special interactions, including halogen bonds that may be formed with aromatic amino acid side chains.^29^ Binding kinetics may differ between closely related isoforms or targets, resulting in kinetic selectivity.^30^ In addition, long residence times may facilitate covalent bond formation, particularly when the targeted cysteine is located in a dynamic structural element such as the glycine-rich loop in LIMK1. We were interested in whether this strategy would be applicable to the design of LIMK1-selective tool compounds. To develop a covalent inhibitor, we therefore synthesized a series of LIMK-i3-based type-I inhibitors, differing in their binding kinetics, before endeavoring to address covalent targeting strategies. LIMK-i3 contains two chlorine atoms *ortho* to the pendant pyrazole heterocycle, suggesting that residence times may be modulated by halogen bond-mediated interactions. In the crystal structure, the two ring systems are oriented orthogonally to each other and appear to lock the binding pocket. As an initial assessment, we analyzed the activity of all synthesized compounds by using the KINETICfinder® platform in TR-FRET binding assays. We further included literature compounds of either type-I, type-II, and type-III binding modes to our kinetic study to select the most favorable lead for designing a covalent LIMK1 inhibitor. In general, the active site-directed type-I inhibitors showed the fastest on-rates but also the shortest target residence times in our screen (Table 1). The potencies of the synthesized type-I inhibitors were between K_d_ = 6.9 nM and K_d_ = 248 nM. In comparison, the type-II inhibitor TH470 had a very long target residence time of 335 min, probably due to the larger structural rearrangements required for the kinase to adopt an inactive DFG-out/αC-out conformation. The on-rate of our type-II inhibitor was significantly slower than the type-I inhibitors by a factor of 10^−3^. For the smaller type-III inhibitors, which bind to the allosteric pocket of LIMK1, the target residence time was between the residence times observed for type-I and type-II inhibitors.

**Table 1.** Kinetic profiling of LIMK1 inhibitors (TR-FRET binding kinetic assay, KINETICfinder®).

| Compd. | Binding mode | R<sub>1</sub>, R<sub>2</sub>, R<sub>3</sub>, R<sub>4</sub> | $k_{on}$ (M<sup>-1</sup>s<sup>-1</sup>) | $k_{off}$ (s<sup>-1</sup>) | $\tau$ (min) | $K_d$ (μM) |
| --- | --- | --- | --- | --- | --- | --- |
| LIMK-i3 (9a) | type-I | Cl, Cl, H, H | $1.12 \times 10^6$ | $4.21 \times 10^{-2}$ | 0.40 | 0.038 |
| 9b | type-I | F, F, H, H | $3.47 \times 10^6$ | $2.43 \times 10^{-2}$ | 0.69 | 0.007 |
| 9c | type-I | Me, Me, H, H | $4.86 \times 10^5$ | $2.92 \times 10^{-2}$ | 0.57 | 0.060 |
| 9d | type-I | H, H, H, H | $2.05 \times 10^5$ | $3.01 \times 10^{-2}$ | 0.55 | 0.147 |
| 9e | type-I | F, Cl, H, H | $1.91 \times 10^6$ | $2.33 \times 10^{-2}$ | 0.72 | 0.012 |
| 9f | type-I | F, Br, H, H | $1.09 \times 10^6$ | $1.82 \times 10^{-2}$ | 0.92 | 0.017 |
| 9g | type-I | H, Cl, H, H | $2.16 \times 10^6$ | $4.85 \times 10^{-2}$ | 0.34 | 0.022 |
| 9h | type-I | H, F, H, H | $1.37 \times 10^6$ | $3.15 \times 10^{-2}$ | 0.53 | 0.023 |
| 9i | type-I | H, Me, H, H | $1.27 \times 10^6$ | $2.10 \times 10^{-2}$ | 0.80 | 0.017 |
| **9j** | type-I | Cl, Cl, Cl,<br>H | $3.07 \times 10^5$ | $3.71 \times 10^{-2}$ | 0.45 | 0.121 |
| **9k** | type-I | Cl, Cl, CF<sub>3</sub>,<br>H | $1.15 \times 10^5$ | $2.58 \times 10^{-2}$ | 0.65 | 0.225 |
| **9l** | type-I | F, Cl, CF<sub>3</sub>,<br>H | $2.50 \times 10^5$ | $2.09 \times 10^{-2}$ | 0.80 | 0.083 |
| **9m** | type-I | F, Cl, CF<sub>3</sub>,<br>F | $1.00 \times 10^5$ | $2.48 \times 10^{-2}$ | 0.67 | 0.248 |
| **TH470** | type-II | n.a. | $2.23 \times 10^4$ | $4.97 \times 10^{-5}$ | 335 | 0.002 |
| **LIJTF-**<br>**500025** | type-III | n.a. | $1.56 \times 10^4$ | $7.48 \times 10^{-4}$ | 22.3 | 0.048 |
| **TH257** | type-III | n.a. | $3.06 \times 10^3$ | $1.75 \times 10^{-3}$ | 9.5 | 0.572 |
| **Stauro-**<br>**sporine** | positive<br>control | n.a. | $1.44 \times 10^7$ | $3.15 \times 10^{-3}$ | 5.29 | 0.0002 |

Although the mono and di-halogenated type-I inhibitors were equally potent, we observed that the binding kinetics were slightly different due to the substitution patterns. The most potent compound in our screen was **9b**, containing two small fluorine substituents in the ortho position. **9b** showed the fastest on-rate with *k_on_* = 3.47×10^6^ with a K_d_ of 6.9 nM. The lead structure LIMK-i3 showed a type-I typical, fast-on rate and a K_d_ of 38 nM. Except for **9l**, the 3-times and 4-times phenyl-decorated compounds **9j**, **9k**, and **9m** showed only weak activity due to the slow on-rate in the range of *k_on_* = 1.0×10^5^ and 3.1×10^5^. A similar behavior was found for the non-decorated LIMK-i3 analogue **9d**.

We next investigated whether the activity and kinetics could be transferred to a cellular system. All compounds were tested in NanoBRET dose-response assays to determine the EC_50_ for LIMK1 and LIMK2 in HEK293T cells (Figure 2A). Gratifyingly, the in-vitro potencies correlated well with the cellular values. Here, the most potent type-I inhibitor was **9e** with an EC_50_ of 0.037 µM. The kinetics of selected type-I inhibitors were analyzed using NanoBRET time-dependent wash-out assays on LIMK1 (Figure 2B).

**Figure 2.**
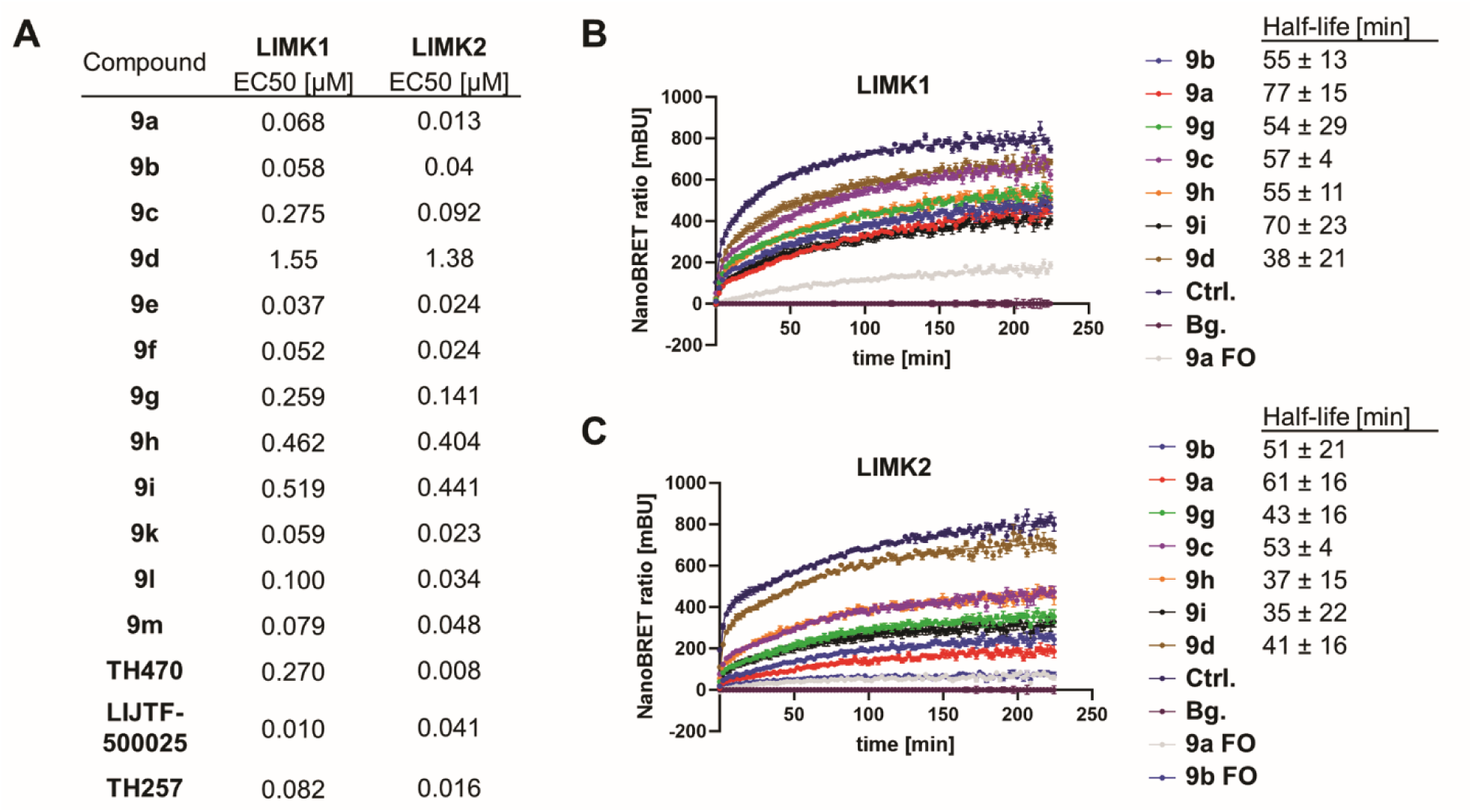
(A) NanoBRET assays in HEK293T cells. (B), (C) Cellular binding kinetics, NanoBRET wash-out assays, of type-I inhibitors towards LIMK1 and LIMK2. FO = full occupancy, no washout was performed for these compounds to simulate the longest possible kinetics. Half-lifes of protein-ligand interactions are depicted (n=3).

Compared to the in vitro experiment, the observed off-rates were significantly longer in the cellular system, possibly due to differences in kinase activation states and the cellular microenvironment that includes protein interaction and competition with cellular cofactors and metabolites. While most compounds showed a similar half-life, the residence time half-life of protein-ligand interactions was with 77 and 70 minutes, respectively, longer for LIMK-i3 (**9a**) and **9i**. Since **9i** showed only weak activity in NanoBRET assays, we believe that the prolonged half-life was due to dissociation via a more conformationally restricted exit from the binding pocket. To investigate if the activity and kinetics differed between isoforms, the experiments were also performed on LIMK2 (Figure 2C). In these experiments, a comparable behavior with slightly weaker EC_50_s and a minimally shorter half-life was detected.

### Development of a LIMK1 selective chemical probe

Our binding analysis, which also included binding kinetics, demonstrated that current reversible inhibitors were not able to discriminate between the two kinases and act as dual LIMK1/LIMK2 inhibitors. Inspired by the publication of Gray et al.,^28^ who explored the kinase cysteinome by using promiscuous covalent inhibitors, demonstrating that cysteines in flexible structural elements can be targeted, we next focused on the design of a LIMK1 selective tool compound. We superimposed the crystal structures of the pan-kinase inhibitor SM1-71 in complex with SRC with the selective inhibitor LIMK-i3 in LIMK1 (Figure 3). Both compounds showed structural overlap in the hinge region and in the glycine-rich loop, which in the case of SM1-71 was folded due to the covalent interaction with Cys280. In the superimposition, the phenyl residue of LIMK1 was oriented with its side chain perpendicular to the electrophilic warhead of SM1-71, which pointed in the direction of the glycine-rich loop. We hypothesized that the phenyl ring system could therefore serve as an optimal attachment point for an electrophilic warhead.

**Figure 3.**
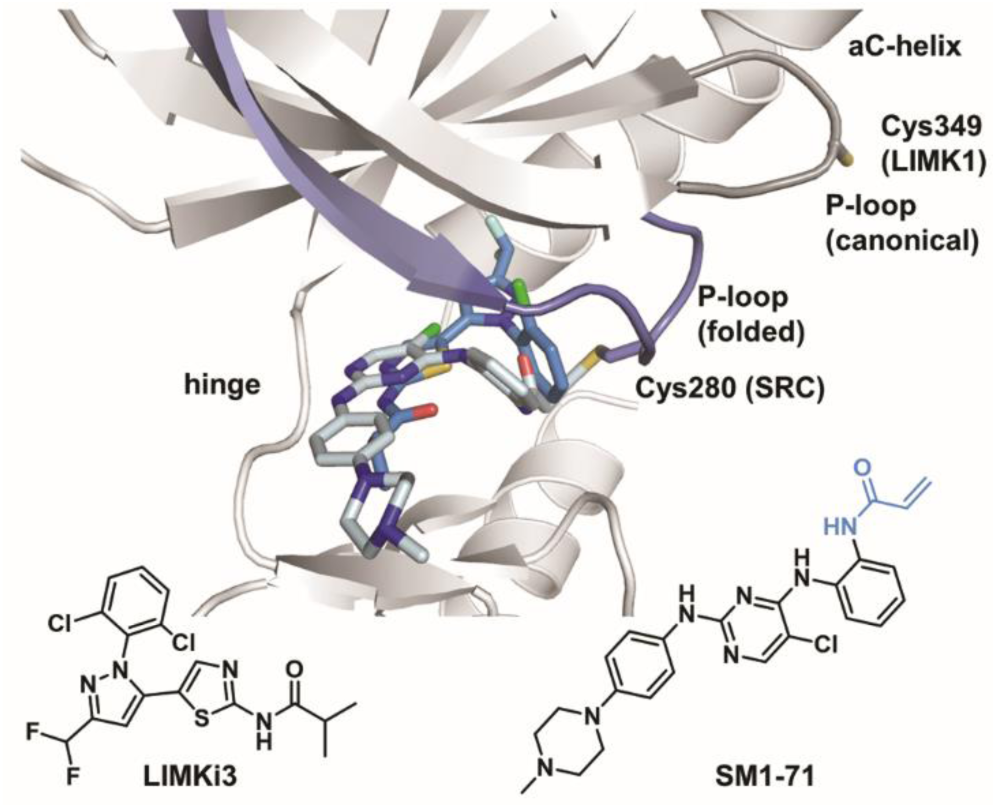
Crystal structure overlay of the promiscuous kinase inhibitor SM1-71 (light blue) in complex with SRC (purple, PDB: 6ATE) with LIMK-i3 (blue) in complex with LIMK1 (gray, PDB: 8AAU). Orientation of the P-loop and reactive cysteine is highlighted in each kinase structure.

For the design of our covalent LIMK1 inhibitor series, we took advantage of the outstanding selectivity profile of the LIMK-i3 parent compound towards LIM kinases. However, the potent activity of LIMK-i3 suggested that introducing an electrophile for selective binding to the glycine-rich loop cysteine in LIMK1 would not result in significant selectivity, as potency for both isoforms was driven by non-discriminating non-covalent interactions. We therefore aimed to select a dual type-I LIMK inhibitor with moderate potency in order to create the desired selectivity gap, while taking advantage of typical type-I inhibitor benefits like fast on-rates and low molecular weight. The in vitro kinetics of the studied pan-LIMK type II and type III inhibitors were already characterized as inhibitors with slow binding kinetics (slow on- and off-rates), offering limited advantages for covalent inhibitor design and associated long target residence times caused by the irreversible binding mode. In contrast, the kinetics of a covalently modified type-I inhibitor could be dramatically improved compared to its non-covalent counterpart, by increasing its fairly fast off-rates and short residence time. We therefore selected **9d** as a starting point for covalent inhibitor development, based on its moderate affinity (NanoBRET assay, EC_50_(LIMK1) = 1.55 µM, EC_50_(LIMK2) = 1.38 µM). The synthetic route of **9d** was modified to allow the introduction of an acrylamide warhead attached to the para-position of the phenyl moiety. We then evaluated the affinity of the resulting inhibitors **10** by measuring the thermal stabilization of LIMK1 and LIMK2 using DSF assays. The covalent compound showed a higher stabilization of LIMK1 with Δ*T_m_* = 8.1 K compared to LIMK2 (Δ*T_m_* = 1.0 K). For the lead structure, **9d**, these values were significantly lower (Δ*T_m_* (LIMK1/2) = 4.0/1.6 K) and equally balanced between both kinases.

To confirm covalent bond formation with LIMK1, mass spectrometry experiments were conducted. The catalytic domain of LIMK1 was incubated with **10** and subsequently analyzed. As predicted, an increase of the native protein mass by 432 Da, corresponding to the molecular weight of the compound, was observed (Figure 4A). To prove that indeed Cys349 in the P-loop of LIMK1 was covalently targeted by our inhibitor, we repeated the MS experiment with the cysteine-deficient LIMK1 variant C349A (Figure 4B). For covalent inhibitors, the kinetics of pre-binding is crucial for specificity. The rate of the chemical reaction (*k_inact_*) with the target can determine the difference between a highly effective inhibitor and a non-specific, toxic molecule. We therefore determined the activity and inactivation kinetics of **10** using the COVALfinder® platform in TR-FRET binding assays. To obtain high-quality data on the binding kinetics of LIMK1 (wt), we monitored the time- and dose-dependent binding to LIMK1 protein over 248 minutes at 20 different concentrations (Figure 4C-F).

**Figure 4.**
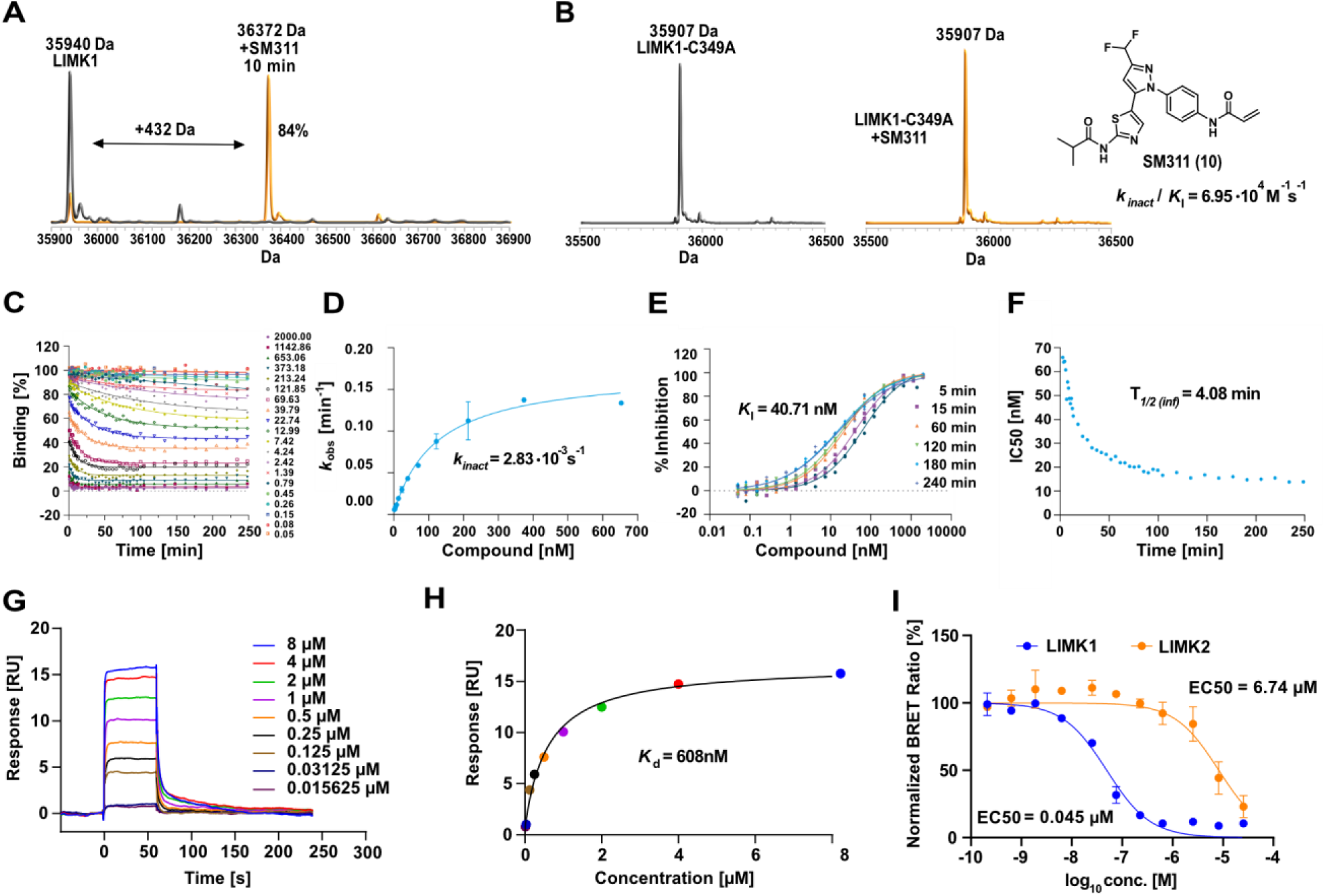
Biochemical evaluation of 10 (SM311). (A) ESI-TOF (Electrospray ionization time-of-flight) mass spectra of LIMK1 before (left panel) and after a 10-minute incubation with **10,** demonstrating rapid covalent bond formation. (B) ESI-TOF mass spectra of the LIMK1/C349A variant in the presence and absence of **10,** showing no bond formation to the mutant protein. (C) Progress curve of LIMK1 generated with COVALfinder TR-FRET kinetic assay after incubation with increasing concentrations of compound **10**. (D) Dependence of *k_obs_* on compound **10** concentration. (E) Dose-response curves over time. (F) IC_50_ values over time. (G, H) Surface plasmon resonance experiments using LIMK1/C349A variant. Calculated K_d_ for the non-covalent interaction with LIMK1/C349A. (I) In-cellular target engagement assay for LIMK1 and LIMK2. NanoBRET was measured in HEK293T cells after 2 hours of incubation.

Compound **10** exhibited a two-step irreversible inactivation mode with *k_inact_* = 2.83×10^−3^ s^−1^ and *k_inact_/K_I_* = 6.95×10^4^ M^−1^ s^−1^. The half-life for inactivation (T_1/2_) of LIMK1 at an infinite concentration of **10** was 4.08 minutes, and a K_I_ value of 40.71 nM was determined. The experiment was followed up in an orthogonal, surface plasmon resonance (SPR) assay with the LIMK1/C349A variant (Figure 4G, 4H). For **10,** a K_d_ = 608 nM was calculated, demonstrating the contribution of the covalent warhead towards binding. To assess the non-specific reactivity towards cysteine residues, we further tested the reactivity towards glutathione. In the GSH assay, **10** showed moderate reactivity towards this abundant cellular thiol, with a half-life of 0.96 hours, which was in the range of the FDA-approved covalent kinase inhibitor Afatinib.

The target engagement of **10** was studied in the cellular environment by NanoBRET assays using both LIMK1/2 isoforms (Figure 4I). The developed covalent compound showed potent binding to LIMK1 with an EC_50_ = 0.045 µM and a 150-fold weaker potency for the closely related LIMK2 isoform (EC_50_ = 6.74 µM).

The selectivity of **10** was tested in a DSF assay using a selection of 91 kinases with staurosporine serving as a positive control (Figure 5A). Within this panel, the covalently modified lead structure showed a clean selectivity profile. Only three kinases were significantly stabilized. The c-Jun N-terminal kinases showed significant Tm shifts (JNK1 (Δ*T_m_* = 7.4 K), and JNK2 (Δ*T_m_* = 8.1 K). JNK1-3 contain a reactive cysteine in the front-pocket region (F3, αD + 2 position) and covalent as well as reversible-covalent inhibitors have been developed previously.^31,32^ As reorientation of **10** could likely lead to covalent binding of these kinases, we analyzed the selectivity and off-target activities using NanoBRET assays, against a panel of 192 kinases at 1 µM compound concentration (Figure 5B). In this assay, the highest tracer displacement, and therefore the most significant target engagement, was found to be LIMK1 (91%), followed by CDKL2 (85%), NLK (73%), TXK (73%), AAK1 (66%), and JNK2 (65%). Interestingly, CDKL2, NLK, and AAK1 also contain a cysteine in the DFG region (D1 cysteine, - position preceding the DFG motif), but no covalent inhibitors have been reported for these kinases so far. We followed up on the cellular activity of **10** in dose-response assays (Figure 5C). Gratifyingly, all targets were only weakly inhibited, with the most potently inhibited off-target (CDKL2) showing only an EC_50_ of 1.51 μM.

**Figure 5.**
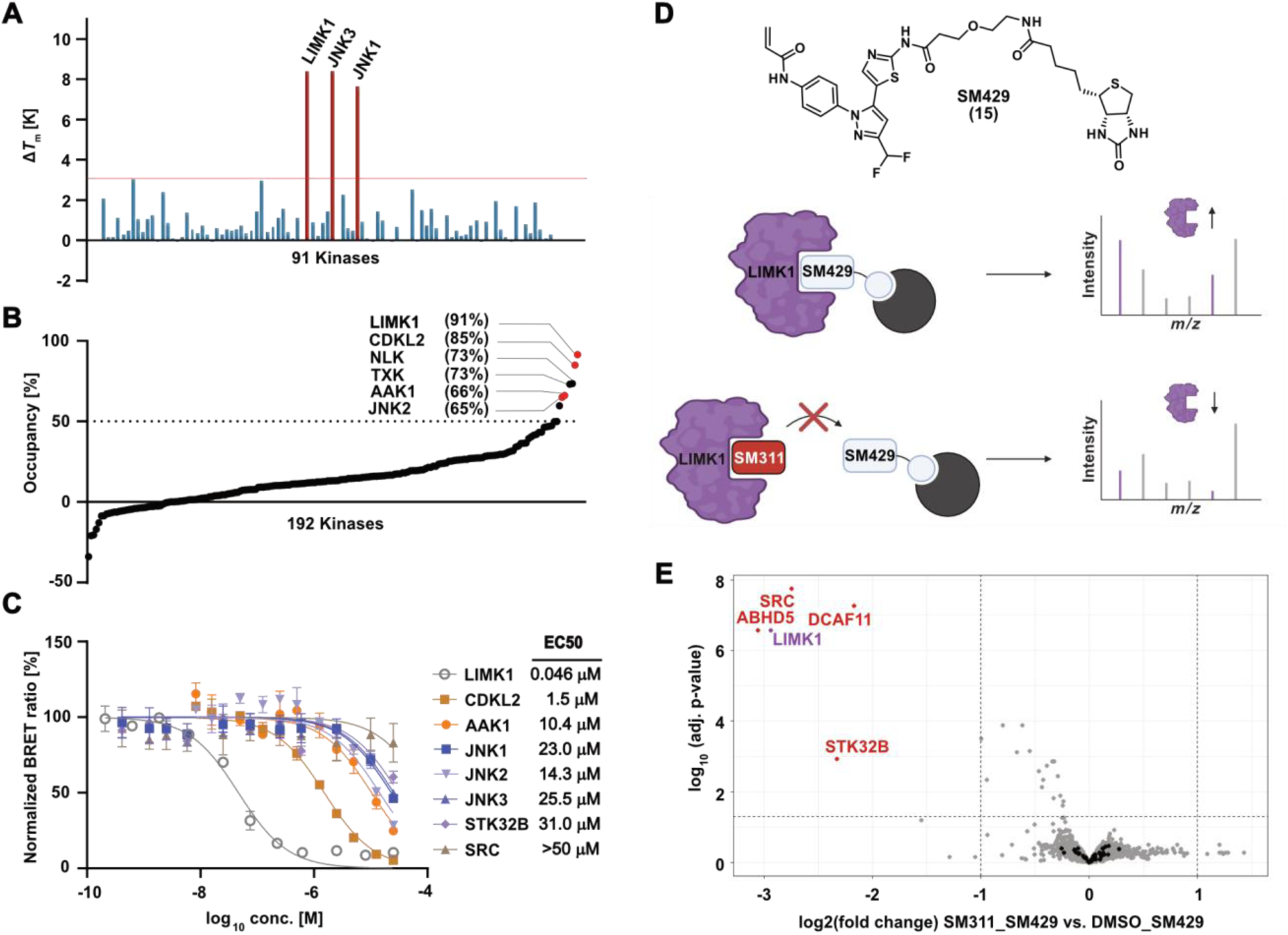
Kinome-wide selectivity and chemo-proteomics profiling of 10 (SM311). (A) DSF screening data using a panel with 91 kinases. Targets with Δ*T_m_* >3 K are labeled. Detailed screening data are available in the supplemental data section (Table S1). (B) Cellular selectivity assessed by a panel of 192 kinases with NanoBRET assays in HEK293T cells. Targets showing tracer displacement above 65% are highlighted. All screening data are available as supplemental data (Table S2). (C) Cellular BRET assays on identified off-targets. LIMK1 showed excellent selectivity against all detected off-targets in our three selectivity screens. (D) Structure of the biotin-modified derivative (**15**) and schematic mechanism of the chemoproteomic competition experiment. (E) Chemo-proteomics competition experiment performed with HEK293 cell lysate for proteome-wide selectivity profiling.

The developed covalent LIMK1 inhibitor might also interact with other reactive cysteines of non-kinase targets. To address this, we performed chemo-proteomics experiments with HEK293 cell lysates. We therefore synthesized a derivative of **10**, that retained the electrophilic warhead and the inhibitor core structure but installed a linker and biotin moiety at the front (solvent)-pocket interacting region of the compound **15**. In the following pulldown experiment, **15** was immobilized on Streptactin beads and incubated with HEK293 lysate in the absence or presence of a 10-fold excess of **10**. While in the absence of **10,** covalent and non-covalent targets and off-targets are expected to be enriched on the beads, the presence of an excess of **10** is expected to saturate specific interactors, leading to a depletion compared to the condition without **10**. Interactors were identified using LC-MS/MS, and the differential enrichment was analyzed in a volcano plot (Figure 5E). As expected, LIMK1 was found to be depleted in the presence of an excess of **10**. Moreover, no peptides for LIMK2, which is also expressed in HEK293 cells, were identified in both conditions.^33^ Additionally, four other proteins were identified as being depleted in the screen (Figure 5E), including the kinases SRC and STK32C, as well as the E3 ligase DCAF11 (DDB1 and CUL4 Associated Factor 11) and Abhydrolase Domain Containing 5 (ABHD5). While no covalent ligands for ABHD5 have been described so far, DCAF11 was among the first E3 ligases to be successfully targeted with covalent inhibitors.^34,35^

In contrast, the detection of SRC as a potential off-target was concerning, as SRC also bears a cysteine residue in its glycine-rich loop.^36^ We therefore tested **10** against SRC using NanoBRET assays, showing no target engagement (EC_50_ >50 µM, Figure 5D), consequently confirming selectivity against this kinase despite the shared cysteine residues at similar positions in the glycine-rich loop. STK32C also showed affinity-driven enrichment and significant depletion, but no tracer molecule has been developed against this poorly studied kinase, preventing us from assessing the selectivity against this kinase. We therefore only annotate the potential interaction of **10** with this kinase based on our chemo-proteomic experiment. Taken together, the results demonstrate a high selectivity of **10** for LIMK1 using three orthogonal screening panels, including selectivity data against the proteome. In addition, all three selectivity assays confirmed isoform selectivity against the closely related LIMK2.

### Cellular potency on endogenous LIMK1 signaling

LIM kinases are key regulators of cofilin phosphorylation, a critical downstream target of the LIM signaling pathway modulating actin function and cell migration. To assess the impact of the covalent inhibitor on cofilin phosphorylation, western blot assays were performed using LN229 glioblastoma cells (Figure 6). Cells were treated with varying concentrations of **10** to evaluate dose-dependent effects on cofilin phosphorylation. The reversible dual LIMK1/2 inhibitor LIMK-i3 was used for comparison. The analysis revealed suppression of cofilin phosphorylation already at 100 nM compound concentration, with increased inhibition observed with up to 1 µM. Notably, LIMK-i3 (**9a**) exhibited the strongest inhibition of cofilin phosphorylation at 1 µM, consistent with its dual activity on both LIMK1 and LIMK2. Since both kinases contribute to cofilin phosphorylation, the more pronounced effect of the dual inhibitor aligned with our expectations of **10** as an isoform-selective LIMK inhibitor.

**Figure 6.**
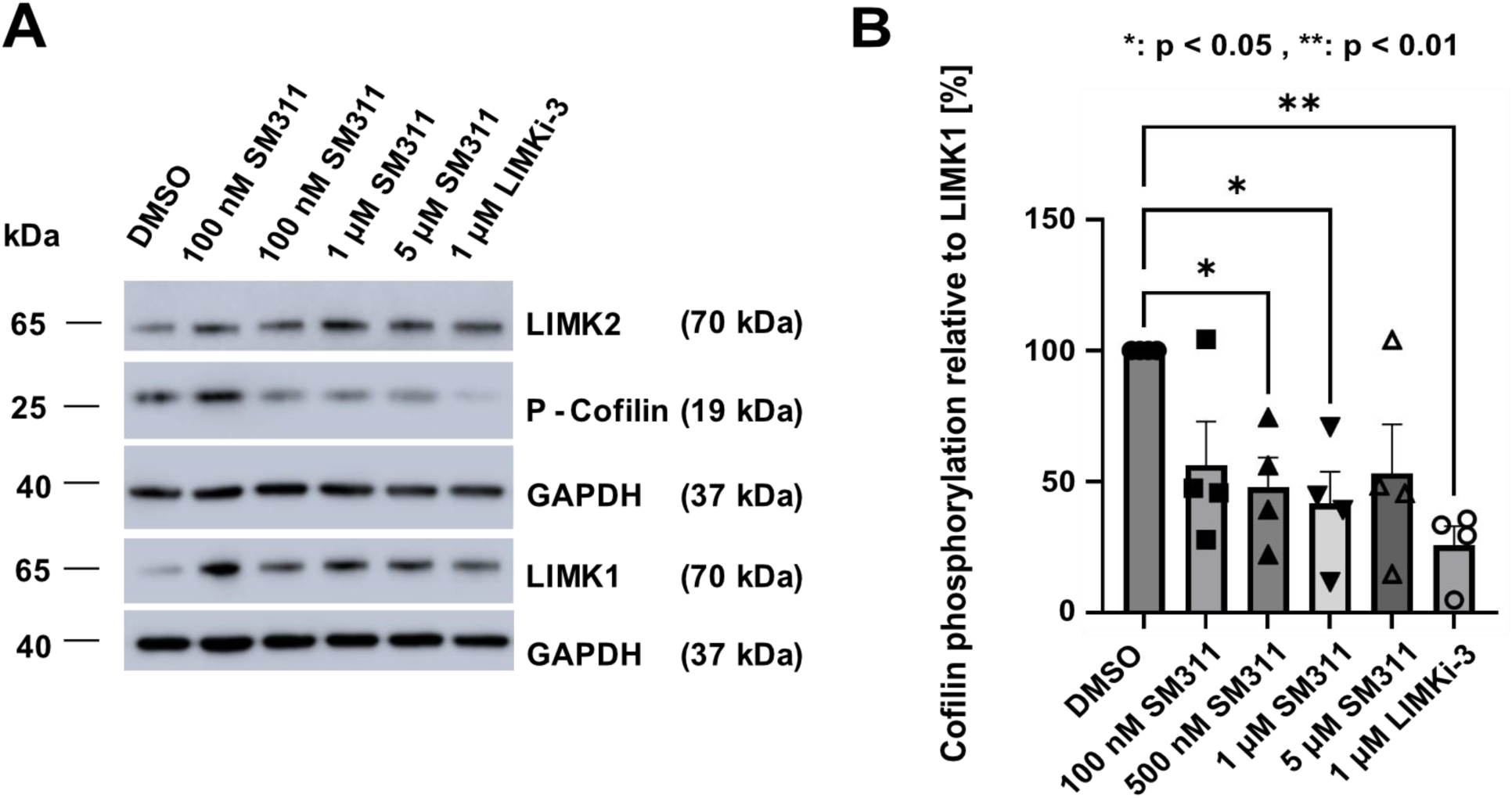
Effect on LIMK1 phosphorylation of the downstream target cofilin. (A) Representative set of Western blot data monitoring cofilin phosphorylation in LN229 glioblastoma cells. LIMK1 selective Inhibitor SM311 (**10**) was investigated at 4 concentrations and compared to dual LIMK1/2 inhibitor LIMK-i3 (**9a**). (B) Quantification of cofilin phosphorylation levels, taking data from 4 biological replicates. Shown are phosphorylation levels compared to the DMSO control (100%) as well as the standard error (SEM) of the data, one star: p<0.05 and two stars: p<0.01.

As the LIMK1 inhibitor and its covalent warhead could affect cellular health, we performed cell viability and cell health assays using a multiplex assay format. **10** was tested in HEK293, U2OS, and MRC-9 cells at two concentrations, and images were taken after 48 hours (Figure 7). At this time point, cells appeared with rounded morphology while nuclei remained intact. When dosed at 1 µM, no cytotoxicity was detected in all three cell lines. However, at 10 µM, the cell viability was significantly reduced after 24 hours of treatment in both HEK293 and USOS cells. An increase in mitochondrial mass and an influence on tubulin structure were observed at 10 µM. Since previous experiments showed a reduction in cofilin phosphorylation, the primary mechanism of action is most likely actin cytoskeleton-related. Therefore, the changes in tubulin dynamics and mitochondrial mass may be secondary effects caused by disruptions of cytoskeletal functions. These effects were less pronounced in the slower-proliferation MRC-9 cells. The expression of phosphatidylserine, as a marker of early apoptosis, was not significantly different from DMSO controls. Based on the excellent activity of **10** in cellular systems, the activity of **10** on off-targets, and the onset of toxicity at 10 µM in some cell lines, we recommend using this LIMK1 inhibitor at concentrations below 1 µM.

**Figure 7:**
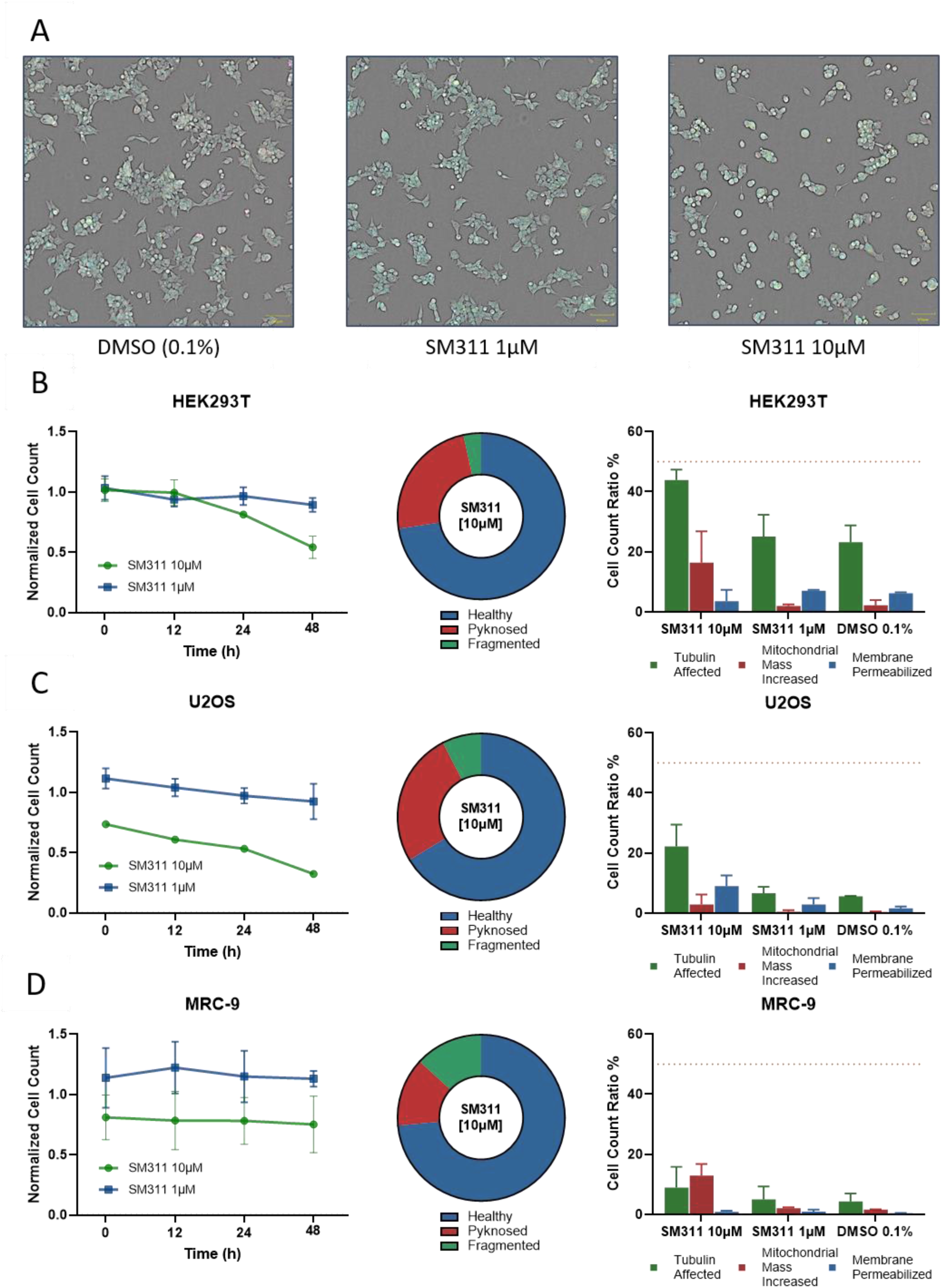
Live-cell viability assessment in HEK293T, U2OS, and MRC-9 cells. (A) Confocal brightfield image (10x) and highlighted fluorescently stained (blue: DNA/Nuclei, green: Microtubule, red: mitochondria, magenta: Annexin apoptosis marker) of representative HEK293T cells after 48 hours of 10 µM and 1 µM compound exposure (SM311, **10**) in comparison to 0.1% DMSO. (B), (C), (D) Normalized cell count over time (0 h, 12 h, 24 h, 48 h) of 10 µM and 1 µM compound exposure (SM311, **10**) against those exposed to 0.1% DMSO in HEK293T (B), U2OS (C), and MRC-9 (D) cells. Fraction of cells with healthy, pyknosed, and fragmented nuclei when exposed to 10 µM displayed as a pie diagram for each cell type. The healthy population was further evaluated for changes in microtubule structure, mitochondrial mass, or membrane permeabilization in contrast to 0.1% DMSO control. The threshold value of 50% is depicted by an orange line. Error bars show SEM of two biological replicates.

## Conclusions

In this work, we developed the first isoform-selective LIMK1 inhibitor by targeting a unique cysteine residue in the P-loop region. This targeting strategy supported our hypothesis that selectivity between the structural homologs LIMK1 and LIMK2 can be achieved by modifying a low potency and fast off-rate inhibitor with an electrophilic warhead targeting a cysteine that is present only in the LIMK1 isoform. Our study also provides evidence that cysteine located in the highly mobile P-loop can be selectively targeted by mild electrophiles. With our covalent targeting approach, we improved target engagement for LIMK1, while retaining key advantages of type-I inhibitors such as fast on-rates and the relatively low molecular weight. The inhibitor discovered, **10** (SM311), showed nanomolar potency in a cellular context and an exceptional selectivity profile not only across the kinome but also for the cellular proteome. The closest off-target within a >30-fold potency window in cellular on-target assays (BRET) was CDKL2. In addition, the developed covalent chemical probe showed low cytotoxicity and a significant effect on cofilin phosphorylation. In summary, the reported covalent LIMK1 inhibitor, **10**, is the ideal probe to study LIMK1-specific signaling that can be used together with previously reported dual LIMK1/2 inhibitors such as TH257, LIJTF500025. In addition, we have developed a structurally related negative control, **16**, which was inactive against both LIMK1 and LIMK2 isoforms, representing a useful control for biological studies in combination with **10** (SM311). We hope that the developed tool compounds will help to further elucidate the detailed roles of LIMK1-specific signaling and may serve as a starting point for the treatment of Fragile X syndrome, in which LIMK1 activity is selectively upregulated.

## Experimental Section

### General Procedures and Chemical Synthesis

Starting materials were purchased from commercial suppliers and used without further purification. NMR spectra were recorded on a Bruker Avance (500 MHz, 400 MHz, 300 MHz, or 250 MHz), and chemical shifts (δ) are reported in ppm, using residual protic solvent as reference. Ready-to-use ALUGRAM Xtra SIL G/UV254 polyester sheets from Machery-Nagel were used for thin-layer chromatography. Detection was achieved using UV light at the wavelengths λ = 254 nm and λ = 336 nm. The puriFlash^®^ X420Plus system with a UV-VIS multi-wave detector (200 – 400 nm) from Interchim and packed silica gel cartridges were used for purifications by column chromatography. For normal phase: PF-SIHP silica columns with a particle size of 15, 30, or 50 µm, and reversed phase: RP-C18 columns. The Agilent HPLC system 1260 Infinity II was used to evaluate the purity of the compounds. This system included a single quadrupole LC/MSD system, InfinityLab (G6125B, ESI pos. 100-1000), a diode array detector 1260 DAD HS (G7177C), a flexible pump (G7104C), a multi-column thermostat (G7166A), a multisampler (G7167A), and a column compartment (G7117A). A Poroshell 120 EC-C18 column from Agilent (3.0 x 150 mm, 2.7 µm) served as the stationary phase. A gradient of H_2_O (A) /ACN (B) with 0.1% formic acid was used as the mobile phase. UV detection was performed at 254 nm, 280 nm, and 310 nm. Two different gradient methods were used with a flow rate of 0.6 mL/min: method 1: 0 min, 5% B – 2 min, 80% B – 5 min, 95% B – 7 min, 95% B. Method 2: 0 min, 5% B – 0.4 min 5% B – 8 min, 100% B – 10 min, 100% B. The chemicals purchased commercially were used without further purification. All final compounds were obtained with a purity of >95% by HPLC analysis. Mass spectrometry (ESI+/ESI-) was measured on a VG Platform II spectrometer from Fisons.HRMS was measured on a ThermoScientific MALDI LTQ Orbitrap XL or a Bruker micOTOF.

### Synthesis of 1-(2,4-dimethoxybenzyl)thiourea (2)

1,1-Thiocarbonyldiimidazole (2.00 g, 11.22 mmol, 1.1 eq.) was suspended in DCM (dry, 50 mL). 2,4-Dimethoxybenzylamine (1.5 mL, 9.96 mmol, 1.0 eq) was dissolved in DCM (dry, 30 mL) and added dropwise over 3 h at RT. Subsequently, a solution of ammonia in MeOH (2M, 25 mL) was added, and the mixture was stirred for an additional 16 h at RT. The solvent was removed under reduced pressure, and the resultant yellow oil was dissolved in DCM (5 mL) and precipitated with *n*-hexane (30 mL). The suspension was filtered, washed with water, and dried under reduced pressure. The product was obtained as a colourless solid. Yield: 1.68 g (7.42 mmol, 75%). ^1^H NMR (400 MHz, DMSO-d_6_): δ = 7.65 (s, 3H), 7.13 (d, *J* = 7.8 Hz, 3H), 6.98 (s, 5H), 6.56 (d, *J* = 2.3 Hz, 3H), 6.49 (d, *J* = 8.2 Hz, 3H), 4.46 (s, 5H), 3.79 (s, 9H), 3.75 (s, 9H) ppm. LC-MS (ESI+): m/z [M+H]^+^ calculated: 227.0. Found: 227.0.

### Synthesis of (*E*)-*N*’-((2,4-dimethoxybenzyl)carbamothioyl)-*N*,*N*-dimethylformimidamide (3)

1-(2,4-Dimethoxybenzyl)thiourea (1.53 g, 6.76 mmol, 1 eq) was suspended in EtOH (abs., 7.5 mL). DMF-DMA (1.5 mL, 10.8 mmol, 1.6 eq) was added, and the mixture was stirred for 90 min at 80 °C under an Ar atmosphere. The resultant suspension was cooled to 4 °C, filtered, and washed with cold EtOH (50 mL). The product was obtained as a colorless solid. Yield: 1.60 g (5.69 mmol, 84%). ^1^H NMR (400 MHz, DMSO-d_6_): δ = 8.79 – 8.58 (m, 2H), 7.02 (t, *J* = 7.0 Hz, 1H), 6.56 – 6.39 (m, 2H), 4.63 (d, *J* = 5.9 Hz, 1H), 4.45 (d, *J* = 6.0 Hz, 1H), 3.78 (d, *J* = 5.8 Hz, 3H), 3.73 (s, 3H), 3.12 (s, 3H), 2.98 (d, *J* = 7.6 Hz, 3H) ppm. LC-MS (ESI+): m/z [M+H]^+^ calculated: 282.1. Found: 282.1.

### Synthesis of 1-(2-((2,4-dimethoxybenzyl)amino)thiazol-5-yl)ethan-1-one (4)

(*E*)-*N*’-((2,4-Dimethoxybenzyl)carbamothioyl)-*N*,*N*-dimethylformimidamide (3.03 g, 10,8 mmol, 1 eq) was suspended in ACN (dry, 50 mL), and chloroacetone (1.1 mL, 13.5 mmol, 1.25 eq) was added. The reaction mixture was stirred for 90 min at 75 °C. After cooling to RT, NaHCO_3_ (aq. sat., 30 mL) was added, and the mixture was cooled to 4 °C. The suspension was filtered, washed with water and *n*-hexane/diethyl ether mixture (4:1), and dried. The product was obtained as a colourless solid. Yield: 1.37 g (4.69 mmol, 88%). ^1^H NMR (400 MHz, DMSO-d_6_): δ = 8.82 (s, 1H), 7.96 (s, 1H), 7.15 (d, *J* = 8.3 Hz, 1H), 6.57 (d, *J* = 2.2 Hz, 1H), 6.48 (dd, *J* = 8.3, 2.3 Hz, 1H), 4.37 (s, 2H), 3.80 (s, 3H), 3.74 (s, 3H), 2.34 (s, 3H) ppm. LC-MS (ESI+): m/z [M+H]^+^ calculated: 293.1. Found: 293.1.

### Synthesis of 1-(2-((2,4-dimethoxybenzyl)amino)thiazol-5-yl)-4,4-difluorobutane-1,3-dione (5)

1-(2-((2,4-Dimethoxybenzyl)amino)thiazol-5-yl)ethan-1-one (2.30 g, 7,87 mmol, 1 eq) and diethyl 2,2-difluoromalonate (5.2 mL, 31.4 mmol, 4 eq) were dissolved in a NaOEt/EtOH solution (21%-mw, 9 mL) and stirred for 4 h at 75 °C. After cooling to RT, the pH was adjusted to pH = 5, by adding glacial acetic acid (q.s.). The yellow precipitate was filtered and washed with a mixture of MeOH/H_2_O (1:2) as well as a mixture of *n*-hexane/diethyl ether (4:1). The product was obtained as a yellow solid. Yield: 2.21 g (5.97 mmol, 76%). ^1^H NMR (400 MHz, CDCl_3_): δ = 7.81 (s, 1H), 7.54 (s, 1H), 7.22 (d, *J* = 8.0 Hz, 1H), 6.48 (d, *J* = 9.5 Hz, 2H), 6.11 (s, 1H), 6.05 (t, 1H), 4.36 (s, 2H), 3.81 (d, *J* = 2.8 Hz, 6H). LC-MS (ESI+): m/z [M+H]^+^ calculated: 371.1. Found: 371.1.

### Synthesis of 5-(3-(difluoromethyl)-1-(2,6-difluorophenyl)-1*H*-pyrazol-5-yl)-*N*-(2,4-dimethoxybenzyl)thiazol-2-amine (6b)

1-(2-{[(2,4-Dimethoxyphenyl)methyl]amino}-1,3-thiazol-5-yl)-4,4-difluorobutane-1,3-dione (200 mg, 0.54 mmol, 1 eq) and (2,6-difluorophenyl)hydrazine hydrochloride (117 mg, 0.65 mmol, 1.2 eq) were dissolved in EtOH (abs., 4 mL) and stirred for 16 h at 75 °C. The mixture was cooled to RT and quenched with H_2_O (50 mL) and a solution of NaHCO_3_ (aq., sat., 30 mL). The precipitate was filtered and washed with MeOH/H_2_O (1:2) and diethylether/*n*-hexane (1:4) to obtain the product as a colorless solid. Yield: 215 mg (0.45 mmol, 83%). R_f_-Value TLC: 0.57 (*n*-hexane/EtOAc 1:1). ^1^H NMR (500 MHz, CDCl_3_): δ = 7.67 (tt, *J* = 8.6, 6.2 Hz, 1H), 7.24 (t, *J* = 8.2 Hz, 2H), 7.10 (d, *J* = 8.3 Hz, 1H), 7.00 (s, 1H), 6.83 (s, 1H), 6.78 (t, *J* = 54.7 Hz, 1H), 6.51 (d, *J* = 2.3 Hz, 1H), 6.44 (dd, *J* = 8.3, 2.3 Hz, 1H), 4.30 (s, 2H), 3.80 (s, 3H), 3.77 (s, 3H) ppm. ^13^C NMR (126 MHz, CDCl_3_): δ = 172.4, 162.2, 161.4 (d, *J* = 2.8 Hz), 159.9, 159.43 (d, *J* = 2.9 Hz), 150.5 (t, *J* = 29.4 Hz), 141.2, 139.7, 134.2 (t, *J* = 10.0 Hz), 130.8, 118.8, 117.7 (t, *J* = 16.4 Hz), 113.7 (d, *J* = 3.8 Hz), 113.5 (d, *J* = 3.7 Hz), 112.2 (t, *J* = 233.7 Hz), 111.3, 105.2, 103.4, 99.3, 55.8, 55.7, 44.7 ppm. LC-MS (ESI+): m/z [M+H]^+^ calculated: 479.1. Found: 479.1.

### Synthesis of 5-(3-(difluoromethyl)-1-(2,6-dimethylphenyl)-1*H*-pyrazol-5-yl)-*N*-(2,4-dimethoxybenzyl)thiazol-2-amine (6c)

1-(2-((2,4-Dimethoxybenzyl)amino)thiazol-5-yl)-4,4-difluorobutane-1,3-dione (100 mg, 0.27 mmol, 1 eq) and (2,6-dimethylphenyl)hydrazine hydrochloride (56 mg, 0.32 mmol, 1.2 eq) were dissolved in EtOH (abs., 2 mL). The reaction mixture was heated for 16 h at 75 °C. The mixture was cooled to RT and quenched with H_2_O (50 mL) and a solution of NaHCO_3_ (aq., sat., 30 mL). The formed brown precipitate was filtered and washed with a mixture of MeOH/H_2_O (1:2) and *n-*hexane/diethyl ether (4:1) to obtain the product as a colorless solid. Yield: 99 mg (0.21 mmol, 79%). R_f_-Value TLC: 0.85 (*n*-hexane/EtOAc 1:3). ^1^H NMR (500 MHz, CDCl_3_): δ = 7.32 (t, *J* = 7.6 Hz, 1H), 7.16 (d, *J* = 7.6 Hz, 2H), 7.10 (d, *J* = 8.2 Hz, 1H), 6.79 (s, 1H), 6.72 (t, *J* = 55.0 Hz, 1H), 6.68 (s, 1H), 6.44 (d, *J* = 2.1 Hz, 1H), 6.41 (dd, *J* = 8.2, 2.3 Hz, 1H), 4.27 (d, *J* = 3.5 Hz, 2H), 3.80 (s, 6H), 1.95 (s, 6H), 1.78 (s, 1H) ppm. ^13^C NMR (126 MHz, CDCl_3_): δ = 170.2, 161.0, 158.7, 147.7 (t, *J* = 29.6 Hz), 138.0, 137.8, 137.4, 137.2, 130.6, 130.3, 128.7, 117.6, 112.7, 111.4 (t, *J* = 234.0 Hz), 103.9, 100.9, 98.8, 55.5, 55.5, 45.4, 17.4 ppm. LC-MS (ESI+): m/z [M+H]^+^ calculated: 471.5. Found: 471.1.

### Synthesis of 5-(3-(difluoromethyl)-1-phenyl-1*H*-pyrazol-5-yl)-*N*-(2,4-dimethoxybenzyl)thiazol-2-amine (6d)

1-(2-((2,4-Dimethoxybenzyl)amino)thiazol-5-yl)-4,4-difluorobutane-1,3-dione (300 mg, 0.81 mmol, 1 eq) and phenylhydrazine hydrochloride (108 mg, 1.0 mmol, 1.2 eq) were dissolved in EtOH (abs., 6 mL). The reaction mixture was heated for 22 h at 75 °C. The mixture was cooled to RT and quenched with H_2_O (50 mL) and a solution of NaHCO_3_ (aq., sat., 30 mL). The formed brown precipitate was filtered and washed with a mixture of MeOH/H_2_O (1:2) and *n-*hexane/diethyl ether (4:1) to obtain the product as a colorless solid. Yield: 305 mg (0.69 mmol, 85%). R_f_-Value TLC: 0.84 (*n*-hexane/EtOAc 1:3). ^1^H NMR (500 MHz, CDCl_3_): δ = 7.49–7.37 (m, 5H), 7.11 (d, *J* = 8.2 Hz, 1H), 6.90 (s, 1H), 6.72 (t, *J* = 54.9 Hz, 1H), 6.65 (s, 1H), 6.45 (d, *J* = 2.3 Hz, 1H), 6.41 (dd, *J* = 8.2, 2.3 Hz, 1H), 4.31 (s, 2H), 3.81 (s, 3H), 3.80 (s, 3H), 2.79 (s, 1H) ppm. ^13^C NMR (126 MHz, CDCl_3_): δ = 161.2, 158.7, 147.4 (t, *J* = 29.9 Hz), 139.0 137.5, 136.3, 130.6, 129.5, 129.3, 126.3, 117.0, 111.2 (t, *J* = 234.1 Hz), 104.3, 103.9, 98.8, 55.5, 55.5, 45.8 ppm. LC-MS (ESI+): m/z [M+H]^+^ calculated: 433.1. Found: 443.1.

### Synthesis of 5-(1-(2-chloro-6-fluorophenyl)-3-(difluoromethyl)-1*H*-pyrazol-5-yl)-*N*-(2,4-dimethoxybenzyl)thiazol-2-amine (6e)

1-(2-((2,4-Dimethoxybenzyl)amino)thiazol-5-yl)-4,4-difluorobutane-1,3-dione (200 mg, 0.54 mmol, 1 eq) and (2-chloro-6-fluorophenyl)hydrazine hydrochloride (128 mg, 0.65 mmol, 1.2 eq) were dissolved in EtOH (abs., 8 mL). The reaction mixture was heated for 18 h at 75 °C. The mixture was cooled to RT and quenched with H_2_O (50 mL) and a solution of NaHCO_3_ (aq., sat., 30 mL). The formed brown precipitate was filtered and washed with a mixture of MeOH/H_2_O (1:2) and *n-*hexane/diethyl ether (4:1). The product was purified by flash chromatography on silica (*n-*hexane/EtOAc + 2% TEA) to obtain the product as a colorless solid. Yield: 48 mg (0.097 mmol, 18%). R_f_-Value TLC: 0.41 (*n*-hexane/EtOAc 1:1). ^1^H NMR (500 MHz, CDCl_3_): δ = 7.45 (td, *J* = 5.6, 2.6 Hz, 1H), 7.34 (dt, *J* = 8.2, 1.0 Hz, 1H), 7.17 (td, *J* = 8.4 Hz, 1.1 Hz, 1H), 7.12 (d, *J* = 8.2 Hz, 1H), 6.72 (t, *J* = 5.4 Hz, 1H), 6.68 (s, 1H), 6.45 (d, *J* = 2.4 Hz, 1H), 6.41 (dd, *J* = 8.2, 2.5 Hz, 1H), 5.68 (bs, 1H), 4.30 (d, *J* = 5.6 Hz, 2H), 3.80 (s, 6H) ppm. ^13^C NMR (126 MHz, CDCl_3_): δ =170.6, 161.1, 160.7, 158.8, 149.0 (t, *J* = 29.9 Hz), 139.0, 135.5, 132.1, 130.6, 126.0, 117.6, 115.6, 113.0, 111.7, 111.1, 109.2, 104.0, 102.8, 98.9, 55.6, 55.5, 45.6 ppm. LC-MS (ESI+): m/z [M+H]^+^ calculated: 493.1. Found: 493.1.

### Synthesis of 5-(1-(2-bromo-6-fluorophenyl)-3-(difluoromethyl)-1*H*-pyrazol-5-yl)-*N*-(2,4-dimethoxybenzyl)thiazol-2-amine (6f)

1-(2-((2,4-Dimethoxybenzyl)amino)thiazol-5-yl)-4,4-difluorobutane-1,3-dione (200 mg, 0.54 mmol, 1 eq) and (2-bromo-6-chlorophenyl)hydrazine hydrochloride (157 mg, 0.65 mmol, 1.2 eq) were dissolved in EtOH (abs., 8 mL). The reaction mixture was heated for 18 h at 75 °C. The mixture was cooled to RT and quenched with H_2_O (50 mL) and a solution of NaHCO_3_ (aq., sat., 30 mL). The formed brown precipitate was filtered and washed with a mixture of MeOH/H_2_O (1:2) and *n-*hexane/diethyl ether (4:1). The product was purified by flash chromatography on silica (*n-*hexane/EtOAc + 2% TEA) to obtain **6f** as a colorless solid. Yield: 85 mg (0.16 mmol, 34%). R_f_-Value TLC: 0.51 (*n*-hexane/EtOAc 1:1). ^1^H NMR (500 MHz, CDCl_3_): δ = 7.51 (dt, *J* = 8.2, 1.0 Hz, 1H), 7.39 (td, *J* = 5.6, 2.8 Hz, 1H), 7.20 (td, *J* = 8.3, 1.0 Hz, 1H), 7.12 (d, *J* = 8.2 Hz, 1H), 6.86 (s, 1H), 6.73 (t, *J* = 5.4 Hz, 1H), 6.68 (s, 1H), 6.45 (d, *J* = 2.4 Hz, 1H), 6.41 (dd, *J* = 8.3, 2.4 Hz, 1H), 5.68 (t, *J* = 5.4 Hz, 1H), 4.29 (d, *J* = 5.6 Hz, 2H), 3.80 (s, 6H) ppm. ^13^C NMR (126 MHz, CDCl_3_): δ = 170.6, 161.1, 160.7, 158.8, 149.0 (t, *J* = 29.9 Hz), 139.0, 135.5, 132.1, 130.6, 126.0, 117.6, 115.6, 113.0, 111.7, 111.1, 109.2, 104.0, 102.8, 98.9, 55.6, 55.5, 45.6 ppm. LC-MS (ESI+): m/z [M+H]^+^ calculated: 539.0. Found: 539.0.

### Synthesis of 5-(3-(difluoromethyl)-1-(2-chlorophenyl)-1*H*-pyrazol-5-yl)-*N*-(2,4-dimethoxybenzyl)thiazol-2-amine (6g)

1-(2-((2,4-Dimethoxybenzyl)amino)thiazol-5-yl)-4,4-difluorobutane-1,3-dione (300 mg, 0.81 mmol, 1 eq) and (2-chlorophenyl)hydrazine hydrochloride (182 mg, 1.02 mmol, 1.2 eq) were dissolved in EtOH (abs., 6 mL). The reaction mixture was heated for 22 h at 75 °C. The mixture was cooled to RT and quenched with H_2_O (50 mL) and a solution of NaHCO_3_ (aq., sat., 30 mL). The formed brown precipitate was filtered and washed with a mixture of MeOH/H_2_O (1:2) and *n-*hexane/diethyl ether (4:1) to obtain **6g** as a colorless solid. Yield: 287 mg (0.21 mmol, 74%). R_f_-Value TLC: 0.70 (*n*-hexane/EtOAc 1:3). ^1^H NMR (500 MHz, CDCl_3_): δ = 7.54 (dd, *J* = 8.0, 1.3 Hz, 1H), 7.51–7.44 (m, 2H), 7.44–7.38 (m, 1H), 7.11 (d, *J* = 8.2 Hz, 1H), 6.75 (s, 1H), 6.72 (t, *J* = 54.9 Hz, 1H), 6.67 (s, 1H), 6.45 (d, *J* = 2.3 Hz, 1H), 6.41 (dd, *J* = 8.2, 2.3 Hz, 1H), 4.30 (s, 2H), 3.81 (s, 3H), 3.80 (s, 3H), 2.79 (s, 1H) ppm. ^13^C NMR (126 MHz, CDCl_3_): δ = 170.2, 161.2, 158.7, 148.0 (t, *J* = 29.9 Hz), 137.9, 136.9, 136.5, 133.2, 131.7, 130.7, 130.6, 130.1, 128.0, 116.9, 112.1, 111.1 (t, *J* = 234.5 Hz), 104.0, 102.6, 98.8, 55.5, 55.5, 45.8 ppm. LC-MS (ESI+): m/z [M+H]^+^ calculated: 477.9. Found: 477.1.

### Synthesis of 5-(3-(difluoromethyl)-1-(2-fluorophenyl)-1*H*-pyrazol-5-yl)-*N*-(2,4-dimethoxybenzyl)thiazol-2-amine (6h)

1-(2-{[(2,4-Dimethoxyphenyl)methyl]amino}-1,3-thiazol-5-yl)-4,4-difluorobutane-1,3-dione (200 mg, 0.54 mmol, 1 eq) and (2-fluorophenyl)hydrazine hydrochloride (105 mg, 0.65 mmol, 1.2 eq) were dissolved in EtOH (abs., 4 mL) and heated for 16 h at 75 °C. The mixture was cooled to RT and quenched with H_2_O (50 mL) and a solution of NaHCO_3_ (aq., sat., 30 mL). The precipitate was filtered and washed with MeOH/H_2_O (1:2) and diethylether/*n*-hexane (1:4) to obtain the product as a yellowish solid. Yield: 86 mg (0.19 mmol, 35%). R_f_-Value TLC: 0.54 (*n*-hexane/EtOAc 1:1). ^1^H NMR (500 MHz, CDCl_3_): δ = 7.51–7.44 (m, 2H), 7.28 (d, *J* = 7.5 Hz, 1H), 7.23–7.19 (m, 1H), 7.12 (d, *J* = 8.2 Hz, 1H), 6.84 (s, 1H), 6.71 (t, *J* = 54.9 Hz, 1H), 6.65 (s, 1H), 6.45–6.40 (m, 2H), 5.78 (s, 1H), 4.29 (d, *J* = 4.1 Hz, 2H), 3.80 (s, 6H) ppm. ^13^C NMR (126 MHz, CDCl_3_): δ = 170.5, 161.0, 158.7, 157.6 (d, *J* = 254.3 Hz), 148.2 (t, *J* = 29.8 Hz), 139.0, 138.6, 131.8 (d, *J* = 7.7 Hz), 130.5, 129.7, 127.3 (d, *J* = 12.4 Hz), 125.0 (d, *J* = 4.0 Hz), 117.5, 117.1 (d, *J* = 19.3 Hz), 112.1, 111.1 (t, *J* = 234.4 Hz), 103.9, 103.0, 98.8, 55.5, 55.5, 45.5 ppm. LC-MS (ESI+): m/z [M+H]^+^ calculated: 461.1. Found: 461.1.

### Synthesis of 5-(3-(difluoromethyl)-1-(*o*-tolyl)-1*H*-pyrazol-5-yl)-*N*-(2,4-dimethoxybenzyl)thiazol-2-amine (6i)

1-(2-((2,4-Dimethoxybenzyl)amino)thiazol-5-yl)-4,4-difluorobutane-1,3-dione (100 mg, 0.27 mmol, 1 eq) and (2-methylphenyl)hydrazine hydrochloride (52 mg, 0.33 mmol, 1.2 eq) were dissolved in EtOH (abs., 2 mL). The reaction mixture was heated for 16 h at 75 °C. The mixture was cooled to RT and quenched with H_2_O (50 mL) and a solution of NaHCO_3_ (aq., sat., 30 mL). The formed brown precipitate was filtered and washed with a mixture of MeOH/H_2_O (1:2) and *n-*hexane/diethyl ether (4:1) to obtain the product as a colorless solid. Yield: 94 mg (0.21 mmol, 76%). R_f_-Value TLC: 0.85 (*n*-hexane/EtOAc 1:3). ^1^H NMR (500 MHz, CDCl_3_): δ = 7.47–7.39 (m, 1H), 7.36–7.27 (m, 3H), 7.10 (d, *J* = 8.2 Hz, 1H), 6.77 (s, 1H), 6.71 (t, *J* = 55.0 Hz, 1H), 6.64 (s, 1H), 6.44 (d, *J* = 2.3 Hz, 1H), 6.41 (dd, *J* = 8.2, 2.3 Hz, 1H), 4.27 (d, *J* = 2.1 Hz, 2H), 3.80 (s, 3H), 3.79 (s, 3H), 1.99 (s, 3H), 1.83 (s, 1H) ppm. ^13^C NMR (126 MHz, CDCl_3_): δ = 170.3, 161.0, 158.7, 147.4 (t, *J* = 29.7 Hz), 138.5, 138.3, 138.0, 136.7, 131.3, 130.5, 130.4, 128.4, 127.1, 117.6, 112.8, 111.3 (t, *J* = 234.0 Hz), 103.9, 101.5, 98.8, 55.5, 55.5, 45.4, 17.3 ppm. LC-MS (ESI+): m/z [M+H]^+^ calculated: 457.2. Found: 457.1

### Synthesis of 5-(3-(difluoromethyl)-1-(2,4,6-trichlorophenyl)-1*H*-pyrazol-5-yl)-*N*-(2,4-dimethoxybenzyl)thiazol-2-amine (6j)

1-(2-((2,4-Dimethoxybenzyl)amino)thiazol-5-yl)-4,4-difluorobutane-1,3-dione (200 mg, 0.54 mmol, 1 eq) and (2,4,6-trichlorophenyl)hydrazine (137 mg, 0.64 mmol, 1.2 eq) were dissolved in EtOH (abs., 4 mL). One drop of HCl (conc.) was added to the solution, which was then heated for 17 h at 75 °C. The mixture was cooled to RT and quenched with H_2_O (50 mL) and a solution of NaHCO_3_ (aq., sat., 30 mL). The formed precipitate was filtered and washed with a mixture of MeOH/H_2_O (1:2), followed by *n-*hexane/diethyl ether (4:1). The solid was dried in the vacuum oven to yield 115 mg (211 µmol, 39%) of compound **6j**. R_f_-Value TLC: 0.45 (*n*-hexane/EtOAc 2:1). ^1^H NMR (500 MHz, CDCl_3_): δ = 9.91 (s, 1H), 7.98 (s, 1H), 7.32 (s, 2H), 7.20 (d, *J* = 8.0 Hz, 1H), 6.61 (bs, 1H), 6.48 (d, *J* = 2.4 Hz, 1H), 6.44 (dd, *J* = 8.0, 2.4 Hz, 1H), 6.11 (t, *J* = 54 Hz, 1H), 4.40 (d, *J* = 5.8 Hz, 2H), 3.84 (s, 3H), 3.81 (s, 3H) ppm. ^13^C NMR (126 MHz, CDCl_3_): δ = 185.7, 176.6, 161.5, 158.8, 151.3, 137.7, 130.9, 129.1, 129.0, 128.0, 127.1, 117.2, 116.4, 115.3, 113.7, 113.4, 104.0, 99.0, 55.6, 46.4, 34.5 ppm. LC-MS (ESI+): m/z [M+Na]^+^ calculated: 547.1. Found: 547.1.

### Synthesis of 5-(1-(2,6-dichloro-4-(trifluoromethyl)phenyl)-3-(difluoromethyl)-1*H*-pyrazol-5-yl)-*N*-(2,4-dimethoxybenzyl)thiazol-2-amine (6k)

1-(2-((2,4-Dimethoxybenzyl)amino)thiazol-5-yl)-4,4-difluorobutane-1,3-dione (200 mg, 540 µmol, 1 eq) and (2,6-dichloro-4-(trifluoromethyl)phenyl)hydrazine (158 mg, 0.11 mmol, 1.2 eq) were dissolved in EtOH (abs., 8 mL). One drop of HCl (conc.) was added to the solution, and the reaction mixture was heated for 17 h at 75 °C. The mixture was cooled to RT and quenched with H_2_O (50 mL) and a solution of NaHCO_3_ (aq., sat., 30 mL). The brown precipitate was filtered and washed with a mixture of MeOH/H_2_O (1:2) and *n-*hexane/diethyl ether (4:1). The crude product was purified by RP-flash chromatography (80% H_2_O/ACN → 20% ACN) to obtain **6k** as a colourless solid. Yield: 61 mg (105 µmol, 20%). R_f_-Value TLC: 0.39 (*n*-hexane/EtOAc 2:1). ^1^H NMR (500 MHz, (CD_3_)_2_CO): δ = 8.12 (s, 2H), 7.18 (d, *J* = 8.2 Hz, 1H), 7.10 (s, 1H), 6.92 (s, 1H), 6.91 (t, *J* = 54 Hz, 1H), 6.53 (d, *J* = 2.4 Hz, 1H), 6.44 (dd, *J* = 8.2, 2.4 Hz, 1H), 4.39 (s, 2H), 3.81 (s, 3H), 3.77 (s, 3H) ppm. ^13^C NMR (126 MHz, (CD_3_)_2_CO): δ = 170.9, 161.8, 159.6, 150.2 (t, *J* = 29.5 Hz), 140.8, 140.4, 139.4, 137.5, 134.7 (q, *J* = 34.3 Hz), 131.0, 127.4 (q, *J* = 3.7 Hz), 123.4 (q, *J* = 273 Hz), 119.2, 114.1, 112.2, 110.9, 110.4, 105.0, 102.9, 99.2, 55.8, 55.7, 44.2 ppm. LC-MS (ESI+): m/z [M+Na]^+^ calculated: 579.1. Found: 579.1.

### Synthesis of 5-(1-(2-chloro-6-fluoro-4-(trifluoromethyl)phenyl)-3-(difluoromethyl)-1*H*-pyrazol-5-yl)-*N*-(2,4-dimethoxybenzyl)thiazol-2-amine (6l)

1-(2-((2,4-Dimethoxybenzyl)amino)thiazol-5-yl)-4,4-difluorobutane-1,3-dione (200 mg, 0.54 mmol, 1 eq) and (2-chloro-6-fluoro-4-(trifluoromethyl)phenyl)hydrazine hydrochloride (142 mg, 0.65 mmol, 1.2 eq) were dissolved in EtOH (abs., 8 mL). The reaction mixture was heated for 18 h at 75 °C. The mixture was cooled to RT and quenched with H_2_O (50 mL) and a solution of NaHCO_3_ (aq., sat., 30 mL). The brown precipitate was filtered and washed with a mixture of MeOH/H_2_O (1:2) and *n-*hexane/diethyl ether (4:1). The crude product was purified by flash chromatography on silica (*n-*hexane/ EtOAc + 2% TEA) to obtain **9l** as a colorless solid. Yield 112 mg (0.20 mmol, 37%). R_f_-Value TLC: 0.71 (*n*-hexane/EtOAc 1:1). ^1^H NMR (500 MHz, CDCl_3_): δ = 7.63 (s, 1H), 7.44 (dd, *J* = 8.1, 1.5 Hz, 1H), 7.12 (d, *J* = 8.2 Hz, 1H), 6.87 (s, 1H), 6.71 (t, *J* = 5.4 Hz, 1H), 6.70 (s, 1H), 6.45 (d, *J* = 2.4 Hz, 1H), 6.41 (dd, *J* = 8.3, 2.4 Hz, 1H), 5.81 (t, *J* = 5.3 Hz, 1H), 4.30 (d, *J* = 5.5 Hz, 2H), 3.80 (s, 6H) ppm. ^13^C NMR (126 MHz, CDCl_3_): δ = 170.8, 161.2, 160.4, 158.8, 158.4, 149.6 (t, *J* = 29.9 Hz), 139.4, 139.3, 136.7, 130.6, 123.2 (q, *J* = 3.7 Hz), 117.5, 113.2 (q, *J* = 3.7 Hz), 113.0 (q, *J* = 3.4 Hz), 112.8, 110.9 (d, *J* = 4.2 Hz), 109.0, 104.0, 103.5, 98.9, 55.6, 55.5, 45.6 ppm. LC-MS (ESI+): m/z [M+H]^+^ calculated: 562.9. Found: 562.9.

### Synthesis of 5-(1-(6-chloro-2,3-difluoro-4-(trifluoromethyl)phenyl)-3-(difluoromethyl)-1*H*-pyrazol-5-yl)-*N*-(2,4-dimethoxybenzyl)thiazol-2-amine (6m)

1-(2-((2,4-Dimethoxybenzyl)amino)thiazol-5-yl)-4,4-difluorobutane-1,3-dione (200 mg, 0.54 mmol, 1 eq) and (6-chloro-2,3-difluoro-4-(trifluoromethyl)phenyl)hydrazine (160 mg, 0.65 mmol, 1.2 eq) were dissolved in EtOH (abs., 8 mL). A drop of HCl (conc.) was added and the mixture was heated for 18 h at 75 °C. The mixture was cooled to RT and quenched with H_2_O (50 mL) and a solution of NaHCO_3_ (aq., sat., 30 mL). The brown precipitate was filtered and washed with a mixture of MeOH/H_2_O (1:2) and *n-*hexane/diethyl ether (4:1). The crude product was purified by flash chromatography on silica (*n-*hexane/EtOAc + 2% TEA) to obtain **6m** as a colorless solid. Yield: 129 mg (0.22 mmol, 41%). R_f_-Value TLC: 0.54 (*n*-hexane/EtOAc 1:1). ^1^H NMR (500 MHz, CDCl_3_): δ = 7.60 (d, *J* = 5.7 Hz, 1H), 7.13 (d, *J* = 8.1 Hz, 1H), 6.90 (s, 1H), 6.71 (t, *J* = 5.4 Hz, 1H), 6.70 (s, 1H), 6.46 (d, *J* = 2.4 Hz, 1H), 6.42 (dd, *J* = 8.3, 2.4 Hz, 1H), 5.79 (t, *J* = 5.3 Hz, 1H), 4.32 (d, *J* = 5.6 Hz, 2H), 3.81 (s, 3H), 3.80 (s, 3H) ppm. ^13^C NMR (126 MHz, CDCl_3_): δ = 170.9, 161.2, 158.8, 150.0 (t, *J* = 29.9 Hz), 147.9, 139.7, 139.4, 130.9, 130.8, 130.6, 122.5 (q, *J* = 3.7 Hz), 119.9, 117.4, 112.6, 110.8, 110.4, 108.9, 104.0, 103.8, 98.9, 55.6, 55,5, 45.7 ppm. LC-MS (ESI-): m/z [M+H]^−^ calculated: 579.1. Found: 579.1.

### Synthesis of *N*-{5-[3-(difluoromethyl)-1-(2,6-difluorophenyl)-1*H*-pyrazol-5-yl]-1,3-thiazol-2-yl}-2-methylpropanamide (8b)

5-[3-(Difluoromethyl)-1-(2,6-difluorophenyl)-1*H*-pyrazol-5-yl]-1,3-thiazol-2-amine (85 mg, 0.26 mmol, 1 eq), 2-methylpropanoyl chloride (69 mg, 0.65 mmol, 2.5 eq), and pyridine (dry, 51 mg, 0.65 mmol, 2.5 eq) were dissolved in DCM (dry, 4 mL). The mixture was stirred for 16 h at RT. The mixture was diluted with EtOAc (30 mL) and washed with an HCl solution (aq., 1M, 10 mL). The organic phase was dried over MgSO_4_, filtered, and the solvent was removed under reduced pressure. The crude product was purified by flash chromatography on silica (*n*-hexane/EtOAc) to obtain **8b** as a colorless solid. Yield: 103 mg (0.31 mmol, 85%). R_f_-Value TLC: 0.56 (*n*-hexane/EtOAc 1:1). ^1^H NMR (500 MHz, DMSO-d_6_): δ = 7.65 (bs, 1H), 7.12 (d, *J* = 8.0 Hz, 1H), 6.99 (bs, 2H), 6.55 (d, *J* = 2.3 Hz, 1H), 6.49 (d, *J* = 8.0 Hz, 1H), 4.46 (d, *J* = 4.24 Hz, 2H), 3.79 (s, 3H), 3.74 (s, 3H) ppm. ^13^C NMR (126 MHz, DMSO-d_6_): δ = 183.2, 159.9, 157.9, 129.5, 118.7, 104.3, 98.3, 55.4, 55.2, 42.6 ppm. LC-MS (ESI+): m/z [M+H]^+^ calculated: 398.9. Found: 398.9. LC-MS (ESI-): m/z [M+H]^−^ calculated: 397.1. Found: 397.1.

### Synthesis of 5-(3-(difluoromethyl)-1-(2,6-dimethylphenyl)-1*H*-pyrazol-5-yl)thiazol-2-amine (8c)

5-(3-(Difluoromethyl)-1-(2,6-dimethylphenyl)-1*H*-pyrazol-5-yl)-*N*-(2,4-dimethoxybenzyl)thiazol-2-amine (91 mg, 0.19 mmol) was dissolved in TFA (1.2 mL) and H_2_O (0.12 mL) and stirred at RT for 20 h. The mixture was quenched with NaHCO_3_ (aq. sat., 20 mL), and the resulting precipitate was filtered and dried in a vacuum oven. The product was purified by flash chromatography on silica (*n-*hexane/EtOAc 9:1 → EtOAc) to afford **8c** as a yellowish solid. Yield 58 mg (0.18 mmol, 93%). R_f_-Value TLC: 0.34 (*n*-hexane/EtOAc 1:1). ^1^H NMR (500 MHz, MeOD): δ = 7.40 (t, *J* = 7.6 Hz, 1H), 7.26 (d, *J* = 7.6 Hz, 2H), 6.88 (s, 1H), 6.86 (s, 1H), 6.80 (t, *J* = 54.8 Hz, 1H), 1.95 (s, 6H) ppm. ^13^C NMR (126 MHz, MeOD): δ = 172.5, 149.2 (t, *J* = 28.8 Hz), 139.6, 138.8, 138.4, 138.4, 131.7, 129.8, 113.3, 112.4 (t, *J* = 233.6 Hz), 101.0 (t, *J* = 1.8 Hz), 17.3 ppm. LC-MS (ESI+): m/z [M+H]^+^ calculated: 321.4. Found: 321.4.

### Synthesis of 5-(3-(difluoromethyl)-1-phenyl-1*H*-pyrazol-5-yl)thiazol-2-amine (8d)

5-(3-(Difluoromethyl)-1-phenyl-1*H*-pyrazol-5-yl)-*N*-(2,4-dimethoxybenzyl)thiazol-2-amine (299 mg, 0.68 mmol) was dissolved in TFA (4.8 mL) and H_2_O (0.5 mL) and stirred for 20 h at RT. The mixture was quenched with NaHCO_3_ (aq. sat., 20 mL), and the resulting precipitate was filtered and dried in a vacuum oven. The product was purified by flash chromatography on silica (*n-*hexane/EtOAc 9:1 → EtOAc) to afford **8d** as a yellowish solid. Yield 153 mg (0.52 mmol, 77%). R_f_-Value TLC: 0.31 (*n*-hexane/EtOAc 1:1). ^1^H NMR (500 MHz, MeOD): δ = 7.57–7.50 (m, 3H), 7.47-7.41 (m, 2H), 6.87 (s, 1H), 6.78 (t, *J* = 54.8 Hz, 1H), 6.76 (s, 1H) ppm. ^13^C NMR (126 MHz, MeOD): δ = 172.8, 148.8 (t, *J* = 29.3 Hz), 140.4, 139.7, 138.6, 130.6, 130.5, 127.8, 113.4, 112.5 (t, *J* = 233.3 Hz), 104.5 (t, *J* = 1.8 Hz) ppm. LC-MS (ESI+): m/z [M+H]^+^ calculated: 293.3. Found: 293.3.

### Synthesis of 5-(1-(2-chloro-6-fluorophenyl)-3-(difluoromethyl)-1*H*-pyrazol-5-yl)thiazol-2-amine (8e)

5-(1-(2-Chloro-6-fluorophenyl)-3-(difluoromethyl)-1*H*-pyrazol-5-yl)-*N*-(2,4-dimethoxybenzyl)thiazol-2-amine (48 mg, 0.10 mmol) was dissolved in TFA (2 mL) and H_2_O (0.5 mL) and stirred at RT for 20 h. The mixture was quenched with NaHCO_3_ (aq. sat., 20 mL), and the resulting precipitate was filtered and dried in a vacuum oven. The product was purified by flash chromatography on silica (*n-*hexane/EtOAc 9:1 → EtOAc) to afford **8e** as a yellowish solid. Yield 30 mg (0.087 mmol, 90%). R_f_-Value TLC: 0.21 (*n*-hexane/EtOAc 1:1). ^1^H NMR (500 MHz, MeOD): δ = 7.66 (td, *J* = 8.4, 5.7 Hz, 1H), 7.52 (dt, *J* = 8.3, 1.2 Hz, 1H), 7.39 (td, *J* = 8.7, 1.2 Hz, 1H), 6.95 (s, 1H), 6.87 (s, 1H), 6.81 (t, *J* = 54 Hz, 1H) ppm. ^13^C NMR (126 MHz, MeOD): δ = 172.7, 162.0, 160.0, 150.4 (t, *J* = 29 Hz), 140.9, 139.5, 136.3, 134.3, 127.3 (d, *J* = 3.6 Hz), 116.6, 112.4, 112.3 (t, *J* = 234 Hz), 103.3 ppm. LC-MS (ESI+): m/z [M+H]^+^ calculated: 344.9. Found: 344.9.

### Synthesis of 5-(1-(2-bromo-6-fluorophenyl)-3-(difluoromethyl)-1*H*-pyrazol-5-yl)thiazol-2-amine (8f)

5-(1-(2-Bromo-6-fluorophenyl)-3-(difluoromethyl)-1*H*-pyrazol-5-yl)-*N*-(2,4-dimethoxybenzyl)thiazol-2-amine (85 mg, 0.15 mmol) was dissolved in TFA (2 mL) and H_2_O (0.5 mL) and stirred for 20 h at RT. The mixture was quenched with NaHCO_3_ (aq. sat., 20 mL), and the resulting precipitate was filtered and dried in a vacuum oven. The product was purified by flash chromatography on silica (*n-*hexane/EtOAc 9:1 → EtOAc) to afford the product as a yellowish solid. Yield: 34 mg (0.087 mmol, 59%). R_f_-Value TLC: 0.22 (*n*-hexane/EtOAc 1:1). ^1^H NMR (500 MHz, MeOD): δ = 7.68 (dt, *J* = 8.3, 1.2 Hz, 1H), 7.59 (td, *J* = 8.4, 5.6 Hz, 1H), 7.42 (td, *J* = 8.5, 1.0 Hz, 1H), 6.95 (s, 1H), 6.86 (s, 1H), 6.81 (t, *J* = 54 Hz, 1H) ppm. ^13^C NMR (126 MHz, MeOD): δ = 172.8, 162.0, 159.9, 150.3 (t, *J* = 29 Hz), 140.7, 139.5, 134.8, 130.4, 125.8, 117.3, 112.5, 112.3 (t, *J* = 234 Hz), 103.3 ppm. LC-MS (ESI+): m/z [M+H]^+^ calculated: 388.9. Found: 388.9.

### Synthesis of 5-(1-(2-chlorophenyl)-3-(difluoromethyl)-1*H*-pyrazol-5-yl)thiazol-2-amine (8g)

5-(3-(Difluoromethyl)-1-(2-chlorophenyl)-1*H*-pyrazol-5-yl)-*N*-(2,4-dimethoxybenzyl)thiazol-2-amine (284 mg, 0.60 mmol) was dissolved in TFA (4.2 mL) and H_2_O (0.4 mL) and stirred for 20 h at RT. The mixture was quenched with NaHCO_3_ (aq. sat., 20 mL), and the resulting precipitate was filtered and dried in a vacuum oven. The product was purified by flash chromatography on silica (*n-*hexane/EtOAc 9:1 → EtOAc) to afford the product as a yellowish solid. Yield 150 mg (0.46 mmol, 77%). R_f_-Value TLC: 0.29 (*n*-hexane/EtOAc 1:1). ^1^H NMR (500 MHz, MeOD): δ = 7.68–7.59 (m, 2H), 7.59–7.50 (m, 2H), 6.85 (s, 1H), 6.80 (s, 1H), 6.78 (t, *J* = 54.7 Hz, 1H) ppm. ^13^C NMR (126 MHz, MeOD): δ =172.6, 149.4 (t, *J* = 29.2 Hz), 140.2, 139.2, 138.0, 134.3, 133.2, 131.6, 131.6, 129.3, 113.1, 112.3 (t, *J* = 233.6 Hz), 103.0 (t, *J* = 1.7 Hz) ppm. LC-MS (ESI+): m/z [M+H]^+^ calculated: 327.0. Found: 327.0.

### Synthesis of 5-(3-(difluoromethyl)-1-(2-fluorophenyl)-1H-pyrazol-5-yl)thiazol-2-amine (8h)

5-[3-(Difluoromethyl)-1-(2-fluorophenyl)-1*H*-pyrazol-5-yl]-*N*-[(2,4-dimethoxyphenyl)methyl]-1,3-thiazol-2-amine (86 mg, 0.19 mmol, 1 eq), was dissolved in TFA (1.4 mL) and H_2_O (0.1 mL) and stirred for 16 h at RT. The mixture was quenched with NaHCO_3_ (aq. sat., 20 mL), and the resulting precipitate was filtered and dried in a vacuum oven. The product was purified by flash chromatography on silica (*n-* hexane/EtOAc 9:1 → EtOAc) to afford the product as an off-white solid. Yield 58 mg (0.13 mmol, 71%). R_f_-Value TLC: 0.36 (*n*-hexane/EtOAc 1:1). ^1^H NMR (500 MHz, MeOD): δ = 7.63–7.59 (m, 1H), 7.53 (td, *J* = 7.7, 1.5 Hz, 1H), 7.38–7.32 (m, 2H), 6.89–6.67 (m, 3H) ppm. ^13^C NMR (126 MHz, MeOD): δ =172.6, 159.1 (d, *J* = 252.5 Hz), 149.6 (t, *J* = 29.2 Hz), 140.2, 139.3, 133.5 (d, *J* = 7.9 Hz), 131.1, 128.1 (d, *J* = 12.6 Hz), 126.3 (d, *J* = 4.0 Hz), 117.9 (d, *J* = 19.4 Hz), 113.0, 112.3 (t, *J* = 233.6 Hz), 103.5 ppm. LC-MS (ESI+): m/z [M+H]^+^ calculated: 311.1. Found: 311.1.

### Synthesis of 5-(3-(difluoromethyl)-1-(o-tolyl)-1*H*-pyrazol-5-yl)thiazol-2-amine (8i)

5-(3-(Difluoromethyl)-1-(*o*-tolyl)-1*H*-pyrazol-5-yl)-*N*-(2,4-dimethoxybenzyl)thiazol-2-amine (78 mg, 0.17 mmol) was dissolved in TFA (1.2 mL), and H_2_O (0.12 mL) and stirred for 20 h at RT. The mixture was quenched with NaHCO_3_ (aq. sat., 20 mL), and the resulting precipitate was filtered and dried in a vacuum oven. The product was purified by flash chromatography on silica (*n-*hexane/EtOAc 9:1 → EtOAc) to afford **8i** as a yellowish solid. Yield 41 mg (0.14 mmol, 78%). R_f_-Value TLC: 0.38 (*n*-hexane/EtOAc 1:1). ^1^H NMR (500 MHz, MeOD): δ = 7.50 (td, *J* = 7.7, 1.0 Hz, 1H, 12), 7.44-7.36 (m, 2H), 7.34 (d, *J* = 7.7 Hz, 1H), 6.84 (s, 1H), 6.80 (s, 1H), 6.78 (t, *J* = 54.8 Hz, 1H), 1.98 (s, 3H) ppm. ^13^C NMR (126 MHz, MeOD): δ = 172.5, 148.8 (t, *J* = 29.0 Hz), 139.7, 139.3, 139.1, 138.0, 132.3, 131.8, 129.5, 128.2, 113.5, 112.4 (t, *J* = 233.5 Hz), 102.4 (t, *J* = 1.8 Hz), 17.1 ppm. LC-MS (ESI+): m/z [M+H]^+^ calculated: 307.3. Found: 307.1.

### Synthesis of 5-(3-(difluoromethyl)-1-(2,4,6-trichlorophenyl)-1*H*-pyrazol-5-yl)thiazol-2-amine (8j)

A solution of 5-(3-(difluoromethyl)-1-(2,4,6-trichlorophenyl)-1*H*-pyrazol-5-yl)-*N*-(2,4-dimethoxybenzyl)thiazol-2-amine (115 mg, 0.21 mmol) was dissolved in TFA (3 mL) and H_2_O (0.5 mL) and stirred for 20 h at RT. The mixture was quenched with NaHCO_3_ (aq. sat., 20 mL), and the resulting precipitate was filtered and dried in a vacuum oven. The product was purified by flash chromatography on silica (*n-*hexane/EtOAc 9:1 → EtOAc) to afford the product as a yellowish solid. Yield: 72 mg (0.18 mmol, 87%). R_f_-Value TLC: 0.54 (*n*-hexane/EtOAc 1:1). ^1^H NMR (500 MHz, MeOD): δ = 7.80 (s, 2H), 7.03 (s, 1H), 6.89 (s, 1H), 6.81 (t, *J* = 54 Hz, 1H) ppm. ^13^C NMR (126 MHz, MeOD): δ = 172.7, 150.6 (t, *J* = 29 Hz), 140.7, 139.7, 139.0, 137.6, 134.9, 130.4, 114.1, 112.2, 112.1, 110.4, 103.2 ppm. LC-MS (ESI+): m/z [M+H]^+^ calculated: 394.9. Found: 394.9.

### Synthesis of 5-(1-(2,6-dichloro-4-(trifluoromethyl)phenyl)-3-(difluoromethyl)-1*H*-pyrazol-5-yl)thiazol-2-amine (8k)

A solution of 5-(1-(2,6-dichloro-4-(trifluoromethyl)phenyl)-3-(trifluoromethyl)-1*H*-pyrazol-5-yl)-*N*-(2,4-dimethoxybenzyl)thiazol-2-amine (61 mg, 0.11 mmol) was dissolved in TFA (3 mL) and H_2_O (0.5 mL) and stirred for 20 h at RT. The mixture was quenched with NaHCO_3_ (aq. sat., 20 mL), and the resulting precipitate was filtered and dried in a vacuum oven. The product was purified by flash chromatography on silica (*n-*hexane/EtOAc 9:1 → EtOAc) to afford **8k** as a yellowish solid. Yield: 39 mg (0.09 mmol, 76%). R_f_-Value TLC: 0.48 (*n*-hexane/EtOAc 1:1). ^1^H NMR (500 MHz, MeOD): δ = 8.07 (s, 2H), 7.01 (s, 1H), 6.92 (s, 1H), 6.83 (t, *J* = 54 Hz, 1H) ppm. ^13^C NMR (126 MHz, MeOD): δ = 172.8, 150.9 (t, *J* = 29 Hz), 140.5, 139.9, 139.3, 138.1, 135.7, 135.4, 127.5 (q, *J* = 3.7 Hz), 124.8, 122.6, 112.2 (t, *J* = 234 Hz), 111.8, 103.5 ppm. LC-MS (ESI+): m/z [M+H]^+^ calculated: 428.9. Found: 428.9.

### Synthesis of 5-(1-(2-chloro-6-fluoro-4-(trifluoromethyl)phenyl)-3-(difluoromethyl)-1*H*-pyrazol-5-yl)thiazol-2-amine (8l)

A solution of 5-(1-(2-chloro-6-fluoro-4-(trifluoromethyl)phenyl)-3-(difluoromethyl)-1*H*-pyrazol-5-yl)-*N*-(2,4-dimethoxybenzyl)thiazol-2-amine (112 mg, 0.19 mmol) was dissolved in TFA (3 mL) and H_2_O (0.5 mL) and stirred for 20 h at RT. The mixture was quenched with NaHCO_3_ (aq. sat., 20 mL), and the resulting precipitate was filtered and dried in a vacuum oven. The product was purified by flash chromatography on silica (*n-* hexane/EtOAc 9:1 → EtOAc) to afford **8l** as a yellowish solid. Yield: 43 mg (0.10 mmol, 55%). R_f_-Value TLC: 0.47 (*n*-hexane/EtOAc 1:1). ^1^H NMR (500 MHz, MeOD): δ = 7.96 (s, 1H), 7.86 (dd, *J* = 8.6, 1.5 Hz, 1H), 7.01 (s, 1H), 6.92 (s, 1H), 6.82 (t, *J* = 54 Hz, 1H) ppm. ^13^C NMR (126 MHz, MeOD): δ = 172.8, 161.9, 159.9, 151.1 (t, *J* = 29 Hz), 141.0, 140.0, 137.8, 135.9 (qd, *J* = 9.2, 34 Hz), 130.2 (d, *J* = 15.5 Hz), 124.6 (q, *J* = 3.8 Hz), 114.6 (dq, *J* = 23.8, 3.6 Hz), 112.2 (t, *J* = 234 Hz), 111.6, 103.8 ppm. LC-MS (ESI+): m/z [M+H]^+^ calculated: 412.9. Found: 412.9.

### Synthesis of 5-(1-(6-chloro-2,3-difluoro-4-(trifluoromethyl)phenyl)-3-(difluoromethyl)-1*H*-pyrazol-5-yl)thiazol-2-amine (8m)

A solution of 5-(1-(6-chloro-2,3-difluoro-4-(trifluoromethyl)phenyl)-3-(difluoromethyl)-1*H*-pyrazol-5-yl)-*N*-(2,4-dimethoxybenzyl)thiazol-2-amine (129 mg, 0.22 mmol) was dissolved in TFA (3 mL) and H_2_O (0.5 mL) and stirred for 20 h at RT. The mixture was quenched with NaHCO_3_ (aq. sat., 20 mL), and the resulting precipitate was filtered and dried in a vacuum oven. The product was purified by flash chromatography on silica (*n-* hexane/EtOAc 9:1 → EtOAc) to afford the product as a yellowish solid. Yield: 57 mg (0.13 mmol, 60%). R_f_-Value TLC: 0.52 (*n*-hexane/EtOAc 1:1). ^1^H NMR (500 MHz, MeOD): δ = 7.98 (dd, *J* = 6.0, 1.8 Hz, 1H), 7.05 (s, 1H), 6.93 (s, 1H), 6.83 (t, *J* = 54 Hz, 1H) ppm. ^13^C NMR (126 MHz, MeOD): δ = 172.8, 161.9, 159.9, 151.1 (t, *J* = 29 Hz), 141.0, 140.0, 137.8, 135.9 (qd, *J* = 9.2, 34 Hz), 130.2 (d, *J* = 15.5 Hz), 124.6 (q, *J* = 3.8 Hz), 114.6 (dq, *J* = 23.8, 3.6 Hz), 112.2 (t, *J* = 234 Hz), 111.6, 103.8 ppm. LC-MS (ESI+): m/z [M+H]^+^ calculated: 430.9. Found: 430.9.

### Synthesis of *N*-(5-(3-(difluoromethyl)-1-(2,6-difluorophenyl)-1*H*-pyrazol-5-yl)thiazol-2-yl)isobutyramide (9b)

5-[3-(Difluoromethyl)-1-(2,6-difluorophenyl)-1*H*-pyrazol-5-yl]-1,3-thiazol-2-amine (85 mg, 0.26 mmol, 1 eq), 2-methylpropanoyl chloride (69 mg, 0.65 mmol, 2.5 eq), and pyridine (dry, 51 mg, 0.65 mmol, 2.5 eq) were dissolved in DCM (dry, 4 mL). The mixture was stirred for 16 h at RT. The reaction was diluted with EtOAc (30 mL) and washed three times with an HCl solution (aq., 1M, 10 mL). The organic phase was dried over MgSO_4_, filtered, and the solvent was removed under reduced pressure. The crude product was purified by flash chromatography on silica (*n*-hexane/EtOAc) to obtain the product as a colorless solid. Yield: 103 mg (0.22 mmol, 85%). Rf-Value TLC: 0.56 (n-hexane/EtOAc 1:1). ^1^H NMR (500 MHz, DMSO-d6): δ = 6.94 – 6.84 (m, 1H), 6.62 (s, 1H), 6.45 (t, *J* = 8.2 Hz, 2H), 6.19 (s, 1H), 6.02 (t, *J* = 54.6 Hz, 1H), 1.89 (sept, *J* = 13.7, 6.9 Hz, 1H), 0.37 (s, 3H), 0.36 (s, 3H) ppm. ^13^C NMR (126 MHz, DMSO-d6): δ = 168.2, 151.8 (d, *J* = 3.0 Hz), 149.8 (d, *J* = 3.0 Hz), 141.2 (t, *J* = 29.5 Hz), 131.2, 129.3, 124.8 (t, *J* = 10.0 Hz), 109.3, 108.1 (t, *J* = 16.5 Hz), 104.6, 104.3 (d, *J* = 3.8 Hz), 104.1 (d, *J* = 3.7 Hz), 102.7, 100.9, 95.2, 26.3, 9.9 ppm. LC-MS (ESI+): m/z [M+H]^+^ calculated: 399.1. Found: 399.0.

### Synthesis of *N*-(5-(3-(difluoromethyl)-1-(2,6-dimethylphenyl)-1*H*-pyrazol-5-yl)thiazol-2-yl)isobutyramide (9c)

5-(3-(Difluoromethyl)-1-(2,6-dimethylphenyl)-1*H*-pyrazol-5-yl)thiazol-2-amine (61 mg, 0.16 mmol, 1 eq), 2-methylpropanoyl chloride (43 mg, 0.40 mmol, 2.5 eq), and pyridine (dry, 32 mg, 0.40 mmol, 2.5 eq) were dissolved in DCM (dry, 4 mL). The mixture was stirred for 16 h at RT. The reaction was diluted with EtOAc (30 mL) and washed three times with an HCl solution (aq., 1M, 10 mL). The organic phase was dried over MgSO_4_, filtered, and the solvent was removed under reduced pressure. The crude product was purified by flash chromatography on silica (*n*-hexane/EtOAc) to obtain the product as a colorless solid. Yield: 51 mg (0.13 mmol, 79%). Rf-Value TLC: 0.63 (n-hexane/EtOAc 1:1). ^1^H NMR (500 MHz, MeOD): δ = 7.43 (t, *J* = 7.7 Hz, 1H), 7.31 – 7.24 (m, 3H), 7.01 (s, 1H), 6.83 (t, *J* = 54.8 Hz, 1H), 2.68 (sept, *J* = 6.9 Hz, 1H), 1.95 (s, 6H), 1.17 (s, 3H), 1.16 (s, 3H). ppm. ^13^C NMR (126 MHz, DMSO-d6): δ = 177.5, 160.8, 149.4 (t, *J* = 29.1 Hz), 139.2, 138.4, 138.2, 138.0, 131.8, 129.9, 119.8, 112.4 (t, *J* = 233.6 Hz)., 103.2, 35.7, 19.4, 17.3 ppm. LC-MS (ESI+): m/z [M+H]^+^ calculated: 391.1. Found: 391.1.

### Synthesis of *N*-(5-(3-(difluoromethyl)-1-phenyl-1*H*-pyrazol-5-yl)thiazol-2-yl)isobutyramide (9d)

5-(3-(Difluoromethyl)-1-phenyl-1*H*-pyrazol-5-yl)thiazol-2-amine (12 mg, 0.04 mmol, 1 eq.), 2-methylpropanoyl chloride (15 mg, 0.14 mmol, 3.5 eq), and pyridine (dry, 15 mg, 0.19 mmol, 4.75 eq) were dissolved in DCM (dry, 2 mL). The mixture was stirred for 16 h at RT. The reaction was diluted with EtOAc (30 mL) and washed three times with an HCl solution (aq., 1M, 10 mL). The organic phase was dried over MgSO_4_, filtered, and the solvent was removed under reduced pressure. The crude product was purified by flash chromatography on silica (*n*-hexane/EtOAc) to obtain the product as a colorless solid. Yield: 7 mg (0.019 mmol, 41%). R_f_-Value TLC: 0.67 (*n*-hexane/EtOAc 1:1). ^1^H NMR (500 MHz, DMSO-d6): δ = 7.57–7.48 (m, 3H), 7.47–7.40 (m, 2H), 7.29 (s, 1H), 6.88 (s, 1H), 6.82 (t, *J* = 54.8 Hz, 1H), 2.70 (sept, *J* = 6.9 Hz, 1H), 1.18 (s, 3H), 1.17 (s, 3H) ppm. ^13^C NMR (126 MHz, DMSO-d6): δ = 177.6, 161.0, 148.9 (t, *J* = 29.3 Hz), 140.3, 139.1, 138.0, 130.7, 130.6, 127.6, 120.0, 112.4 (t, *J* = 233.3 Hz), 105.6 (t, *J* = 1.8 Hz), 35.8, 19.4 ppm. LC-MS (ESI-): m/z [M+H]^−^ calculated: 385.1. Found: 385.1.

### Synthesis of *N*-(5-(1-(2-chloro-6-fluorophenyl)-3-(difluoromethyl)-1*H*-pyrazol-5-yl)thiazol-2-yl)isobutyramide (9e)

5-(1-(2-Chloro-6-fluorophenyl)-3-(difluoromethyl)-1*H*-pyrazol-5-yl)thiazol-2-amine (30 mg, 0.09 mmol, 1 eq.), 2-methylpropanoyl chloride (19 mg, 0.18 mmol, 2.5 eq), and pyridine (dry, 14 mg, 0.18 mmol, 2.5 eq) were dissolved in DCM (dry, 2 mL). The mixture was stirred for 16 h at RT. The reaction was diluted with EtOAc (30 mL) and washed three times with an HCl solution (aq., 1M, 10 mL). The organic phase was dried over MgSO_4_, filtered, and the solvent was removed under reduced pressure. The crude product was purified by flash chromatography on silica (*n*-hexane/EtOAc) to obtain the product as a colorless solid. Yield: 11 mg (0.027 mmol, 29%). R_f_-Value TLC: 0.29 (*n*-hexane/EtOAc 2:1). ^1^H NMR (500 MHz, MeOD): δ = 7.66 (td, *J* = 8.4, 5.8 Hz, 1H), 7.52 (dt, *J* = 8.3, 1.2 Hz, 1H), 7.43 (s, 1H), 7.42 (td, *J* = 8.7, 1.2 Hz, 1H), 7.03 (s, 1H), 6.86 (t, *J* = 54 Hz, 1H), 2,72 (hept, *J* = 6.8 Hz, 1H), 1.20 (d, *J* = 6.8 Hz, 6H) ppm. ^13^C NMR (126 MHz, MeOD): δ = 171.7, 161.1 (t, *J* = 29 Hz), 150.6, 140.4, 138.8, 136.1, 134.5,134.4, 127.4, 126.8, 119.0, 116.7, 112.3 (t, *J* = 233.2 Hz), 104.5, 35.8, 19.5, 19.4 ppm. LC-MS (ESI-): m/z [M+H]^−^ calculated: 413.1. Found: 413.1.

### Synthesis of *N*-(5-(1-(2-bromo-6-fluorophenyl)-3-(difluoromethyl)-1*H*-pyrazol-5-yl)thiazol-2-yl)isobutyramide (9f)

5-(1-(2-Bromo-6-fluorophenyl)-3-(difluoromethyl)-1*H*-pyrazol-5-yl)thiazol-2-amine (34 mg, 0.08 mmol, 1 eq.), 2-methylpropanoyl chloride (18 mg, 0.17 mmol, 2.5 eq), and pyridine (dry, 13 mg, 0.17 mmol, 2.5 eq) were dissolved in DCM (dry, 2 mL). The mixture was stirred for 16 h at RT. The reaction was diluted with EtOAc (30 mL) and washed three times with an HCl solution (aq., 1M, 10 mL). The organic phase was dried over MgSO_4_, filtered, and the solvent was removed under reduced pressure. The crude product was purified by flash chromatography on silica (*n*-hexane/EtOAc) to obtain the product as a colorless solid. Yield: 12 mg (0.026 mmol, 29%). R_f_-Value TLC: 0.29 (*n*-hexane/EtOAc 2:1). ^1^H NMR (500 MHz, MeOD): δ = 7.70 (td, *J* = 8.4, 5.8 Hz, 1H), 7.62 (dt, *J* = 8.3, 1.2 Hz, 1H), 7.45 (td, *J* = 8.7, 1.2 Hz, 1H), 7.43 (s, 1H), 7.03 (s, 1H), 6.86 (t, *J* = 54 Hz, 1H), 2,72 (hept, *J* = 6.8 Hz, 1H), 1.21 (d, *J* = 6.8 Hz, 6H) ppm. ^13^C NMR (126 MHz, MeOD): δ = 177.7, 161.1 (t, *J* = 29 Hz), 150.5, 140.3, 138.8, 136.1, 134.5,134.4, 125.7, 117.4, 117.2, 114.1, 112.3 (t, *J* = 233.2 Hz), 104.5, 35.8, 19.5, 19.4 ppm. LC-MS (ESI-): m/z [M+H]^−^ calculated: 459.0. Found: 459.0.

### Synthesis of *N*-{5-[1-(2-chlorophenyl)-3-(difluoromethyl)-1*H*-pyrazol-5-yl]-1,3-thiazol-2-yl}-2-methylpropanamide (9g)

5-[1-(2-Chlorophenyl)-3-(difluoromethyl)-1*H*-pyrazol-5-yl]-1,3-thiazol-2-amine (78 mg, 0.24 mmol, 1 eq), 2-methylpropanoyl chloride (64 mg, 0.60 mmol, 2.5 eq), and pyridine (dry, 47 mg, 0.60 mmol, 2.5 eq) were dissolved in DCM (dry, 2 mL). The mixture was stirred for 16 h at RT. The reaction was diluted with EtOAc (30 mL) and washed three times with a HCl solution (aq., 1M, 10 mL). The organic phase was dried over MgSO_4_, filtered, and the solvent was removed under reduced pressure. The crude product was purified by flash chromatography on silica (*n*-hexane/EtOAc) to obtain the product as a colorless solid. Yield: 95 mg (0.24 mmol, 62%). R_f_-Value TLC: 0.56 (*n*-hexane/EtOAc 1:1). ^1^H NMR (500 MHz, MeOD): δ = 7.67 – 7.58 (m, 1H), 7.55 (m, 1H), 7.29 (s, 1H), 6.95 (s, 1H), 6.82 (t, *J* = 54.7 Hz, 1H), 2.76 – 2.62 (m, 1H), 1.18 (s, 1H), 1.16 (s, 1H). ppm. ^13^C NMR (126 MHz, MeOD): δ = 177.6, 160.9, 149.5 (t, *J* = 29.2 Hz), 139.7, 138.6, 137.9, 134.1, 133.3, 131.7, 131.5, 129.5, 119.7, 112.3 (t, *J* = 233.6 Hz), 104.2, 49.0, 35.7, 19.4.ppm. LC-MS (ESI+): m/z [M+Na]^+^ calculated: 419.05. Found: 419.0.

### Synthesis of *N*-{5-[3-(difluoromethyl)-1-(2-fluorophenyl)-1H-pyrazol-5-yl]-1,3-thiazol-2-yl}-2-methylpropanamide (9h)

5-[3-(Difluoromethyl)-1-(2-fluorophenyl)-1*H*-pyrazol-5-yl]-1,3-thiazol-2-amine (41 mg, 0.13 mmol, 1 eq), 2-methylpropanoyl chloride (35 mg, 0.33 mmol, 2.5 eq) and dry pyridine (dry, 26 mg, 0.22 mmol, 2.5 eq) were dissolved in DCM (dry, 2 mL) and stirred for 16 h at RT. The mixture was diluted with EtOAc (30 mL) and washed with an HCl solution (aq., 1M, 10 mL). The organic phase was dried over MgSO_4_, filtered, and the solvent was removed under reduced pressure. The crude product was further purified by flash chromatography on silica (*n*-hexane/EtOAc) to obtain *N*-{5-[3-(difluoromethyl)-1-(2-fluorophenyl)-1*H*-pyrazol-5-yl]-1,3-thiazol-2-yl}-2-methyl propanamide as a colorless solid. Yield 50 mg (0.13 mmol, 86%). R_f_-Value TLC: 0.55 (*n*-hexane/EtOAc 1:1). ^1^H NMR (500 MHz, CD_3_OD): δ = 7.65–7.61 (m, 1H), 7.57 (td, *J* = 7.7, 1.6 Hz, 1H), 7.40 (t, *J* = 7.7 Hz, 1H), 7.36–7.32 (m, 2H), 6.93–6.71 (m, 2H), 2.69 (hept, *J* = 6.9 Hz, 1H), 1.17 (d, *J* = 6.9 Hz, 6H) ppm. ^13^C NMR (126 MHz, CD_3_OD): δ = 177.7, 161.0, 159.0 (d, *J* = 252.2 Hz), 149.8 (t, *J* = 29.3 Hz), 139.8, 138.7, 133.7 (d, *J* = 7.9 Hz), 131.1, 128.2 (d, *J* = 12.5 Hz), 126.5 (d, *J* = 4.0 Hz), 119.6, 118.0 (d, *J* = 19.4 Hz), 112.4 (t, *J* = 233.6 Hz), 104.8, 35.8, 19.5 ppm. LC-MS (ESI+): m/z [M+Na]^+^ calculated: 402.9. Found: 402.9.

### Synthesis of *N*-(5-(3-(difluoromethyl)-1-(*o*-tolyl)-1*H*-pyrazol-5-yl)thiazol-2-yl)isobutyramide (9i)

5-(3-(Difluoromethyl)-1-(*o*-tolyl)-1*H*-pyrazol-5-yl)thiazol-2-amine (64 mg, 0.21 mmol, 1 eq.), 2-methylpropanoyl chloride (55 mg, 0.52 mmol, 2.5 eq), and pyridine (dry, 41 mg, 0.52 mmol, 2.5 eq) were dissolved in DCM (dry, 2 mL). The mixture was stirred for 16 h at RT. The reaction was diluted with EtOAc (30 mL) and washed three times with a HCl solution (aq., 1M, 10 mL). The organic phase was dried over MgSO_4_, filtered, and the solvent was removed under reduced pressure. The crude product was purified by flash chromatography on silica (*n*-hexane/EtOAc) to obtain the product as a colorless solid. Yield: 41 mg (0.11 mmol, 53%). R_f_-Value TLC: 0.51 (*n*-hexane/EtOAc 2:1). ^1^H NMR (500 MHz, MeOD): δ = 7.53 (td, *J* = 7.5, 1.4 Hz, 1H), 7.47 – 7.38 (m, 2H), 7.36 (dd, *J* = 7.8, 1.4 Hz, 1H), 7.25 (s, 1H), 6.95 (s, 1H), 6.82 (t, *J* = 54.8 Hz, 1H), 2.68 (dt, *J* = 13.7, 6.9 Hz, 1H), 1.98 (s, 3H), 1.17 (s, 3H), 1.16 (s, 3H) ppm. ^13^C NMR (126 MHz, DMSO-d6): δ = 177.6, 160.8, 149.0 (t, *J* = 29.1 Hz), 139.3, 139.2, 138.3, 137.8, 132.4, 131.9, 129.5, 128.3, 120.0, 112.4 (t, *J* = 233.5 Hz), 103.7, 35.7, 19.4, 17.1 ppm. LC-MS (ESI+): m/z [M+Na]^+^ calculated: 399.1 Found: 399.1

### Synthesis of *N*-(5-(3-(difluoromethyl)-1-(2,4,6-trichlorophenyl)-1*H*-pyrazol-5-yl)thiazol-2-yl)isobutyramide (9j)

5-(3-(Difluoromethyl)-1-(2,4,6-trichlorophenyl)-1*H*-pyrazol-5-yl)thiazol-2-amine (72 mg, 0.17 mmol, 1 eq.), 2-methylpropanoyl chloride (38 mg, 0.36 mmol, 2.5 eq), and pyridine (dry, 28 mg, 0.36 mmol, 2.5 eq) were dissolved in DCM (dry, 2 mL). The mixture was stirred for 16 h at RT. The reaction was diluted with EtOAc (30 mL) and washed three times with a HCl solution (aq., 1M, 10 mL). The organic phase was dried over MgSO_4_, filtered, and the solvent was removed under reduced pressure. The crude product was purified by flash chromatography on silica (*n*-hexane/EtOAc) to obtain the product as a colorless solid. Yield: 34 mg (0.073 mmol, 41%). R_f_-Value TLC: 0.51 (*n*-hexane/EtOAc 2:1). ^1^H NMR (500 MHz, MeOD): δ = 7.80 (s, 2H), 7.46 (s, 2H), 7.03 (s, 1H), 6.85 (t, *J* = 54 Hz, 1H), 2.72 (hept, *J* = 6.8 Hz, 1H), 1.19 (d, *J* = 6.8 Hz, 6H) ppm. ^13^C NMR (126 MHz, MeOD): δ = 181.1, 117.8, 161.1, 150.8 (t, *J* = 29 Hz), 140.2, 139.1, 138.9, 137.4, 134.8, 130.5, 118.7, 112.2 (t, *J* = 233.6 Hz), 104.5, 35.8, 19.5, 19.4 ppm. LC-MS (ESI-): m/z [M-H]^−^ calculated: 463.0. Found: 463.0.

### Synthesis of *N*-(5-(1-(2,6-dichloro-4-(trifluoromethyl)phenyl)-3-(difluoromethyl)-1*H*-pyrazol-5-yl)thiazol-2-yl)isobutyramide (9k)

5-(1-(2,6-Dichloro-4-(trifluoromethyl)phenyl)-3-(difluoromethyl)-1*H*-pyrazol-5-yl)thiazol-2-amine (34 mg, 0.08 mmol, 1 eq.), 2-methylpropanoyl chloride (16 mg, 0.16 mmol, 2.5 eq), and pyridine (dry, 13 mg, 0.16 mmol, 2.5 eq) were dissolved in DCM (dry, 2 mL). The mixture was stirred at RT for 16 h. The reaction was diluted with EtOAc (30 mL) and washed three times with an HCl solution (aq., 1M, 10 mL). The organic phase was dried over MgSO_4_, filtered, and the solvent was removed under reduced pressure. The crude product was purified by flash chromatography on silica (*n*-hexane/EtOAc) to obtain the product as a colorless solid. Yield: 26 mg (90.05 mmol, 26%). R_f_-Value TLC: 0.64 (*n*-hexane/EtOAc 2:1). ^1^H NMR (500 MHz, MeOD): δ = 8.08 (s, 2H), 7.43 (s, 2H), 7.06 (s, 1H), 6.88 (t, *J* = 54 Hz, 1H), 2.73 (hept, *J* = 6.8 Hz, 1H), 1.20 (d, *J* = 6.8 Hz, 6H) ppm. ^13^C NMR (126 MHz, MeOD): δ = 161.1, 151.1 (t, *J* = 29 Hz), 140.0, 139.2, 138.9, 137.9, 135.6 (q, *J* = 34.4 Hz), 127.6 (q, *J* = 3.7 Hz), 126.9, 124.7, 122.6, 120.4, 118.5, 112.2 (t, *J* = 233.2 Hz), 104.9, 35.8, 19.4, 19.1 ppm. C-MS (ESI-): m/z [M-H]^−^ calculated: 497.0. Found: 497.0.

### Synthesis of *N*-(5-(1-(2-chloro-6-fluoro-4-(trifluoromethyl)phenyl)-3-(difluoromethyl)-1*H*-pyrazol-5-yl)thiazol-2-yl)isobutyramide (9l)

5-(1-(2-Chloro-6-fluoro-4-(trifluoromethyl)phenyl)-3-(difluoromethyl)-1*H*-pyrazol-5-yl)thiazol-2-amine (43 mg, 0.10 mmol, 1 eq.), 2-methylpropanoyl chloride (28 mg, 0.26 mmol, 2.5 eq), and pyridine (dry, 21 mg, 0.26 mmol, 2.5 eq) were dissolved in DCM (dry, 2 mL). The mixture was stirred for 16 h at RT. The reaction was diluted with EtOAc (30 mL) and washed three times with a HCl solution (aq., 1M, 10 mL). The organic phase was dried over MgSO_4_, filtered, and the solvent was removed under reduced pressure. The crude product was purified by flash chromatography on silica (*n*-hexane/EtOAc) to obtain the product as a colorless solid. Yield: 25 mg (0.05 mmol, 51%). R_f_-Value TLC: 0.62 (*n*-hexane/EtOAc 2:1). ^1^H NMR (500 MHz, MeOD): δ = 7.96 (s, 1H), 7.86 (dd, *J* = 8.7, 1.5 Hz, 1H), 7.43 (s, 1H), 7.06 (s, 1H), 6.88 (t, *J* = 54 Hz, 1H), 2.73 (hept, *J* = 6.8 Hz, 1H), 1.22 (d, *J* = 6.8 Hz, 6H) ppm. ^13^C NMR (126 MHz, MeOD): δ = 177.8, 161.8, 161.2, 159.7, 151.2 (t, *J* = 29 Hz), 140.4, 139.0, 137.5, 135.8, 130.2, 124.7, 122.6, 118.4, 114.7, 112.2 (t, *J* = 233.2 Hz), 105.2, 35.8, 19.4 ppm. LC-MS (ESI-): m/z [M-H]^−^ calculated: 481.0. Found: 481.0.

### Synthesis of *N*-(5-(1-(6-chloro-2,3-difluoro-4-(trifluoromethyl)phenyl)-3-(difluoromethyl)-1*H*-pyrazol-5-yl)thiazol-2-yl)isobutyramide (9m)

5-(1-(6-Chloro-2,3-difluoro-4-(trifluoromethyl)phenyl)-3-(difluoromethyl)-1*H*-pyrazol-5-yl)thiazol-2-amine (57 mg, 0.13 mmol, 1 eq), 2-methylpropanoyl chloride (28 mg, 0.26 mmol, 2.5 eq), and pyridine (dry, 21 mg, 0.26 mmol, 2.5 eq) were dissolved in DCM (dry, 2 mL). The mixture was stirred for 16 h at RT. The reaction was diluted with EtOAc (30 mL) and washed three times with an HCl solution (aq., 1M, 10 mL). The organic phase was dried over MgSO_4_, filtered, and the solvent was removed under reduced pressure. The crude product was purified by flash chromatography on silica (*n*-hexane/EtOAc) to obtain the product as a colorless solid. Yield: 36 mg (0.072 mmol, 55%). Rf-Value TLC: 0.62 (*n*-hexane/EtOAc 2:1). ^1^H NMR (500 MHz, MeOD): δ = 7.98 (dd, *J* = 8.7, 1.5 Hz, 1H), 7.48 (s, 1H), 7.08 (s, 1H), 6.89 (t, *J* = 54 Hz, 1H), 2.73 (hept, *J* = 6.8 Hz, 1H), 1.21 (d, *J* = 6.8 Hz, 6H) ppm. ^13^C NMR (126 MHz, MeOD): δ = 177.9, 161.3, 151.3 (t, *J* = 29 Hz), 151.0, 149.8, 149.0, 147.7, 140.5, 139.3, 131.7, 124.4 (q, J = 4.8 Hz), 121.4, 118.1, 112.1 (t, *J* = 233.2 Hz), 105.5, 35.9, 19.5, 19.4 ppm. LC-MS (ESI-): m/z [M-H]^−^ calculated: 499.0. Found: 499.0.

### Synthesis of *N*-(4-(3-(difluoromethyl)-5-(2-isobutyramidothiazol-5-yl)-1*H*-pyrazol-1-yl)phenyl)acrylamide (10)

*N*-(5-(1-(4-Aminophenyl)-3-(difluoromethyl)-1*H*-pyrazol-5-yl)thiazol-2-yl)isobutyramide (50 mg, 132 μmol, 1 eq) and DIEA (69 μL, 379 μmol, 3 eq) were dissolved in ACN (dry, 10 mL) and acryloyl chloride (12 µL, 146 µmol, 1.1 eq) was added. The reaction mixture was stirred for 16 h at RT. A solution of NaHCO_3_ (aq., sat., 5 mL) was added, and the mixture was extracted three times with DCM (each 50 mL). The organic phase was dried over MgSO_4_, filtered, and the solvent was removed under reduced pressure. The crude product was purified by RP-flash chromatography (95% H_2_O/ACN → 100% ACN) to yield 17 mg (393 µmol, 30%) of a colorless solid. ^1^H NMR (400 MHz, DMSO-d_6_): δ = 12.25 (s, 1H), 10.46 (s, 1H), 7.84 (d, *J* = 8.8 Hz, 2H), 7.57 (s, 1H), 7.44 (d, *J* = 8.7 Hz, 2H), 7.07 (d, *J* = 19.1 Hz, 2H), 6.49 (dd, *J* = 17.0, 10.1 Hz, 1H), 6.32 (dd, *J* = 16.9, 1.7 Hz, 1H), 5.88 – 5.79 (m, 1H), 2.72 (dt, *J* = 13.9, 7.0 Hz, 1H), 1.09 (d, *J* = 6.8 Hz, 6H) ppm. ^13^C NMR (75 MHz, DMSO-d_6_) δ = 175.4, 163.5, 159.2, 146.9, 146.5, 140.0, 138.3, 136.6, 133.6, 131.6, 127.5, 127.2, 119.6, 117.6, 111.3, 108.2, 104.3, 33.7, 19.0 ppm. LC-MS (ESI+): m/z [M+H]^+^ calculated: 432.1. Found: 432.1. HRMS: m/z [M+H]^+^ calculated for [C20H19F2N5O2S]: 432.1300. Found: 432.1293.

### Synthesis of 5-(3-(difluoromethyl)-1-(4-nitrophenyl)-1*H*-pyrazol-5-yl)-*N*-(2,4-dimethoxybenzyl) thiazol-2-amine (11)

1-(2-((2,4-Dimethoxybenzyl)amino)thiazol-5-yl)ethan-1-one (500 mg, 1.35 mmol, 1.0 eq) was suspended in EtOH (abs., 15 mL), and *para*-nitrophenylhydrazine (307 mg, 1.62 mmol, 1.2 eq) was added. The reaction mixture was stirred for 4 h at 75 °C. The orange solution was diluted with H_2_O (10 mL) and NaHCO_3_ (aq. sat., 5 mL). The mixture was extracted three times with DCM (each 50 mL). The organic phase was dried over MgSO_4,_ filtered, and the solvent was removed under reduced pressure. The product was obtained as a brown solid. Yield: 557 mg (1.14 mmol, 85%). ^1^H NMR (400 MHz, DMSO-d_6_): δ = 8.36 (d, *J* = 9.0 Hz, 2H), 8.15 (t, *J* = 5.5 Hz,1H), 7.76 (d, *J* = 9.0 Hz, 2H), 7.17 (s, 1H), 7.15 – 7.07 (m, 1H), 6.95 (s, 1H), 6.53 (d, *J* = 2.2 Hz, 1H), 6.46 (dd, *J* = 8.3, 2.3 Hz, 1H), 4.29 (d, *J* = 5.5 Hz, 2H), 3.76 (s, 3H), 3.74 (s, 3H) ppm. LC-MS (ESI+): m/z [M+H]^+^ calculated: 488.1. Found: 488.1.

### Synthesis of 5-(3-(difluoromethyl)-1-(4-nitrophenyl)-1*H*-pyrazol-5-yl)thiazol-2-amine (12)

5-(3-(Difluoromethyl)-1-(4-nitrophenyl)-1*H*-pyrazol-5-yl)-*N*-(2,4-dimethoxybenzyl)thiazol-2-amine (533 mg, 1.09 mmol, 1 eq) was dissolved in TFA (6 mL) and H_2_O (0.6 mL). The reaction mixture was stirred for 3 h at 40 °C. H_2_O (30 mL) was added, and the solution was neutralized with NaHCO_3_ (aq., sat., 20 mL). The precipitate was filtered, washed with water, and dried under reduced pressure. The product was obtained as a brown solid. Yield: 236 mg (0.70 mmol, 64%). ^1^H NMR (400 MHz, DMSO-d_6_): δ = 8.38 (d, *J* = 8.9 Hz, 2H), 7.77 (d, *J* = 8.9 Hz, 2H), 7.38 (s, 2H), 7.13 (s, 4H), 7.10 (s, 1H), 6.97 (s, 1H) ppm. LC-MS (ESI+): m/z [M+H]^+^ calculated: 338.0. Found: 338.1.

### Synthesis of *N*-(5-(3-(difluoromethyl)-1-(4-nitrophenyl)-1*H*-pyrazol-5-yl)thiazol-2-yl)-isobutyramide (13)

5-(3-(Difluoromethyl)-1-(4-nitrophenyl)-1*H*-pyrazol-5-yl)thiazol-2-amine (230 mg, 681 μmol, 1 eq) and DIEA (78.6 μL, 750 μmol, 1.1 eq) were dissolved in ACN (dry, 10 mL). Isobutryl chloride (131 μL, 750 μmol, 1.1 eq) was dissolved in ACN (dry, 5 mL) and added dropwise to the solution. The mixture was stirred for 2 h at RT. NaHCO_3_ (aq., sat., 20 mL) was added, and the mixture was extracted three times with DCM (each 20 mL). The organic phase was dried over MgSO_4_, filtered, and the solvent was removed under reduced pressure. The crude product was purified via RP-flash chromatography (95% H_2_O/ACN → 100% ACN) to yield 132 mg (323 µmol, 47%) of **13** as a yellowish solid. ^1^H NMR (400 MHz, DMSO-d_6_): δ = 12.33 (s, 1H), 8.36 (d, *J* = 9.0 Hz, 2H), 7.80 – 7.72 (m, 2H), 7.54 (s, 1H), 7.14 (s, 1H), 7.10 (s, 1H), 2.84 – 2.63 (m, 1H), 1.09 (d, *J* = 6.9 Hz, 6H) ppm. LC-MS (ESI+): m/z [M+H]^+^ calculated: 408.1. Found: 408.1.

### Synthesis of *N*-(5-(1-(4-aminophenyl)-3-(difluoromethyl)-1*H*-pyrazol-5-yl)thiazol-2-yl)isobutyramide (14)

*N*-(5-(3-(difluoromethyl)-1-(4-nitrophenyl)-1*H*-pyrazol-5-yl)thiazol-2-yl)isobutyramide (150 mg, 368 μmol, 1 eq), iron (144 mg, 2.58 mmol, 7 eq) and ammonium chloride (138 mg, 2.58 mmol, 7 eq) were suspended in MeOH (13 mL) and H_2_O (2 mL). The suspension was stirred over 16 h at 75 °C. The reaction mixture was filtered over Celite® and the solvent was removed under reduced pressure. The residue was dissolved in DCM (50 mL) and washed three times with NaHCO_3_ (aq., sat., each 20 mL). The organic phase was dried over MgSO_4_, filtered, and the solvent was removed under reduced pressure. The product was obtained as a colorless solid. Yield: 100 mg (259 µmol, 72%). ^1^H NMR (300 MHz, DMSO-d_6_): δ = 12.19 (s, 1H), 7.57 (s, 1H), 7.08–7.04 (m, 2H), 6.99 (s, 1H), 6.71– 6.56 (m, 2H), 5.58 (s, 2H), 2.72 (q, 1H), 1.09 (d, *J* = 5.9 Hz, 6H) ppm. LC-MS (ESI+): m/z [M+H]^+^ calculated: 378.4. Found: 378.1.

### Synthesis of *N*-(2-(3-((5-(1-(4-acrylamidophenyl)-3-(difluoromethyl)-1*H*-pyrazol-5-yl)thiazol-2-yl)amino)-3-oxopropoxy)ethyl)-5-((3*aS*,4*S*,6*aR*)-2-oxohexahydro-1*H*-thieno[3,4-*d*]imidazol-4-yl)pentanamide (15)

*N*-(2-(3-((5-(1-(4-Aminophenyl)-3-(difluoromethyl)-1*H*-pyrazol-5-yl)thiazol-2-yl)amino)-3-oxopropoxy)ethyl)-5-((3*aS*,4*S*,6*aR*)-2-oxohexahydro-1*H*-thieno[3,4-*d*]imidazol-4-yl)pentanamide (34 mg, 52.4 μmol, 1 eq) and DIEA (27 μL, 157 μmol, 3 eq) were dissolved in ACN (dry, 5 mL). Chloroacetyl chloride (4.7 μL, 57.7 μmol, 1.1 eq) was added, and the reaction was then stirred for 3 h at RT. The solvent was removed under reduced pressure and the crude product was purified via RP-flash chromatography (95% H_2_O/ACN → 100% ACN). The product was obtained as a colorless solid. Yield: 20 mg (28.4 µmol, 54%). ^1^H NMR (400 MHz, DMSO-d_6_): δ = 12.30 (s, 1H), 10.44 (s, 1H), 7.82 (d, *J* = 8.8 Hz, 2H), 7.76 (t, *J* = 5.5 Hz, 1H), 7.56 (s, 1H), 7.42 (d, *J* = 8.8 Hz, 2H), 7.26–6.92 (m, 1H), 7.04 (s, 1H), 6.53–6.26 (m, 4H), 5.81 (dd, *J* = 10.1, 1.9 Hz, 1H), 4.33–4.25 (m, 1H), 4.16–4.06 (m, 1H), 3.65 (t, *J* = 6.2 Hz, 2H), 3.37 (d, *J* = 6.0 Hz, 2H), 3.14 (q, *J* = 5.8 Hz, 2H), 3.10–3.03 (m, 1H), 2.80 (dd, *J* = 12.4, 5.1 Hz, 1H), 2.65 (t, *J* = 6.2 Hz, 2H), 2.56 (d, *J* = 12.4 Hz, 1H), 2.03 (t, *J* = 7.4 Hz, 2H), 1.59 (ddd, *J* = 15.7, 10.9, 6.1 Hz, 1H), 1.55–1.37 (m, 3H), 1.36– 1.20 (m, 2H) ppm. ^13^C NMR (101 MHz, DMSO-d_6_): δ = 172.6, 170.1, 164.0, 163.2, 159.4, 147.0 (t, *^2^J_CF_* = 28.8 Hz), 140.5, 138.8, 137.0, 134.1, 132.1, 128.0, 127.7, 120.1, 118.1, 111.8 (t, *^1^J_CF_* = 232.5 Hz), 104.8, 69.3, 66.13, 61.5, 59.7, 55.9, 38.8, 36.0, 35.6, 28.7, 28.5, 25.7 ppm. LC-MS (ESI+): m/z [M+2H]^+^ calculated: 703.2. Found: 703.2. HRMS: m/z [M+H]^+^ calculated for [C31H36F2N8O5S2]: 703.2291. Found: 703.2274.

### Synthesis of *N*-(4-(3-(difluoromethyl)-5-(2-(*N*-methylisobutyramido)thiazol-5-yl)-1*H*-pyrazol-1-yl)phenyl)acrylamide (16)

*N*-(5-(1-(4-Aminophenyl)-3-(difluoromethyl)-1*H*-pyrazol-5-yl)thiazol-2-yl)-*N*-methylisobutyramide (35 mg, 89.4 µmol, 1 eq) and DIEA (47 µL, 268 μmol, 3 eq) were dissolved in ACN (dry, 5 mL). Acryloyl chloride (11 μL, 134 µmol, 1.5 eq) was added, and the mixture was stirred for 16 h at RT. The reaction was quenched with H_2_O (10 mL), and the solvent was removed under reduced pressure. The residue was dissolved in EtOAc (20 mL) and washed three times with NaHCO_3_ (aq., sat., 20 mL). The organic phase was dried over MgSO_4_, filtered, and the solvent was removed under reduced pressure. The crude product was purified by preparative HPLC (RP, 95% H_2_O/ACN → 100% ACN) to yield 8.7 mg (19.5 µmol, 22%) of a colorless solid. ^1^H NMR (500 MHz, DMSO-d_6_) δ = 10.42 (s, 1H), 7.81 (d, *J* = 8.8 Hz, 2H), 7.58 (s, 1H), 7.52 – 7.38 (m, 2H), 7.21 – 6.96 (m, 1H), 7.02 (s, 1H), 6.46 (dd, *J* = 17.0, 10.1 Hz, 1H), 6.30 (dd, *J* = 17.0, 1.9 Hz, 1H), 5.81 (dd, *J* = 10.1, 1.9 Hz, 1H), 3.66 (s, 3H), 3.18 (dt, *J* = 13.4, 6.7 Hz, 1H), 1.10 (d, *J* = 6.7 Hz, 6H) ppm. ^13^C NMR (126 MHz, DMSO-d_6_): δ = 177.0, 163.94, 160.8, 147.0 (t, ^2^*J*_CF_ = 28.8 Hz), 140.4, 138.0, 136.9, 134.1, 132.1, 128.0, 127.6, 120.1, 119.7, 111.5 (t, ^1^J_CF_ = 232.5 Hz), 105.0, 55.4, 34.9, 31.6, 19.3 ppm. LC-MS (ESI+): m/z [M+H]^+^ calculated: 446.1. Found: 446.1. HRMS: m/z [M+H]^+^ calculated for [C22H22F2N4O2S]: 446.1457. Found: 446.1451.

### Synthesis of 5-(3-(difluoromethyl)-1-(4-nitrophenyl)-1*H*-pyrazol-5-yl)-*N*-(2,4-dimethoxybenzyl)-*N*-methylthiazol-2-amine (17)

5-(3-(Difluoromethyl)-1-(4-nitrophenyl)-1*H*-pyrazol-5-yl)-*N*-(2,4-dimethoxybenzyl)-*N*-thiazol-2-amine (100 mg, 205 μmol, 1 eq) was dissolved in DMF (dry, 10 mL) and NaH (60% on mineral oil, 9.03 mg, 226 µmol, 1,1 eq) was added at 0 °C. The solution was stirred for 10 min at 0 °C. The ice cooling was removed, and MeI (2M in THF, 102 µL, 205 μmol, 1 eq) was added. The mixture was then stirred for 16 h at RT. The reaction was quenched with ammonia (aq., 25%, 10 mL), and the solvent was removed under reduced pressure. The crude product was purified via RP-flash chromatography (95% H_2_O/ACN → 100% ACN). The product was obtained as a yellow solid. Yield: 52 mg (102 µmol, 50%). ^1^H NMR (400 MHz, CDCl_3_): δ = 8.28 (d, *J* = 8.7 Hz, 2H), 7.65 (d, *J* = 8.7 Hz, 2H), 7.15 (s, 1H), 7.06 (d, *J* = 8.2 Hz, 1H), 6.87–6.55 (m, 2H), 6.45 (s, 1H), 6.43 (d, *J* = 8.5 Hz, 1H), 4.52 (s, 2H), 3.79 (d, *J* = 7.7 Hz, 6H), 3.09 (s, 3H) ppm. LC-MS (ESI+): m/z [M+H]^+^ calculated: 503.1. Found: 502.1.

### Synthesis of 5-(3-(difluoromethyl)-1-(4-nitrophenyl)-1*H*-pyrazol-5-yl)-*N*-methylthiazol-2-amine (18)

5-(3-(Difluoromethyl)-1-(4-nitrophenyl)-1*H*-pyrazol-5-yl)-*N*-(2,4-dimethoxybenzyl)-*N*-methylthiazol-2-amine (250 mg, 499 μmol, 1 eq) was dissolved in TFA (4.5 mL) and H_2_O (0.5 mL) and stirred for 4 h at 55 °C. The mixture was quenched with NaHCO_3_ (aq. sat., 20 mL), and the resulting precipitate was filtered and dried in a vacuum oven. The product was obtained as a brown solid. Yield 175 mg (499 µmol, quant.). ^1^H NMR (400 MHz, CDCl_3_): δ = 8.29–8.24 (m, 2H), 7.61–7.58 (m, 2H), 7.36–7.33 (m, 1H), 6.80 (s, *J* = 5.3 Hz, 1H), 6.92–6.58 (m, 1H), 3.76 (s, 3H), 1.26 (d, *J* = 6.8 Hz, 6H) ppm. LC-MS (ESI+): m/z [M+H]^+^ calculated: 352.1. Found: 352.0.

### Synthesis of *N*-(5-(3-(difluoromethyl)-1-(4-nitrophenyl)-1*H*-pyrazol-5-yl)thiazol-2-yl)-*N*-methylisobutyramide (19)

5-(3-(Difluoromethyl)-1-(4-nitrophenyl)-1*H*-pyrazol-5-yl)-*N*-methylthiazol-2-amine (100 mg, 284 μmol, 1 eq) was dissolved in DMF (dry, 10 mL) and NaH (60% in mineral oil, 12.5 mg, 313 μmol, 1.1 eq) was added at 0 °C. The reaction was stirred for 10 min at 0 °C, and isobutryl chloride (33.0 µL, 313 µmol, 1.1 eq) was added. The mixture was then stirred for 1 h at RT. The reaction was quenched by carefully adding H_2_O (50 mL), and the solvents were removed under reduced pressure. The residue was dissolved in EtOAc (50 mL) and washed three times with NaHCO_3_ (aq., sat., 50 mL). The organic phase was dried over MgSO_4_, filtered, and the solvent was removed under reduced pressure. The crude product was purified via RP-flash chromatography (95% H_2_O/ACN → 100% ACN) to obtain 175 mg (284 μmol, quant.) of a yellow solid. ^1^H NMR (400 MHz, CDCl_3_): δ = 8.24 (d, *J* = 9.0 Hz, 2H), 7.59 (d, *J* = 9.0 Hz, 2H), 7.34 (s, 1H), 6.77 (s, 1H), 6.90 – 6.56 (m, 1H), 3.74 (s, 3H), 3.09 (q, *J* = 13.5, 6.7 Hz, 1H), 1.24 (d, *J* = 6.8 Hz, 6H) ppm. LC-MS (ESI+): m/z [M+H]^+^ calculated: 422.1. Found: 422.1.

### Synthesis of *N*-(5-(1-(4-aminophenyl)-3-(difluoromethyl)-1*H*-pyrazol-5-yl)thiazol-2-yl)-*N*-methylisobutyramide (20)

*N*-(5-(3-(Difluoromethyl)-1-(4-nitrophenyl)-1*H*-pyrazol-5-yl)thiazol-2-yl)-*N*-methylisobutyramide (45 mg, 107 μmol, 1 eq), iron (42 mg, 747 µmol, 7 eq), and ammonium chloride (40 mg, 747 µmol, 7 eq) were suspended in MeOH (9 mL) and H_2_O (1 mL) and stirred for 2 h at 75 °C. The reaction mixture was filtered over Celite®, and the solvent was removed under reduced pressure. The residue was dissolved in EtOAc (20 mL) and washed three times with NaHCO_3_ (aq., sat., 20 mL). The organic phase was dried over MgSO_4_, filtered, and the solvent was removed under reduced pressure. The crude product was purified via RP-flash chromatography (95% H_2_O/ACN → 100% ACN) to yield 28 mg (71.5 µmol, 66%) of a colorless solid. ^1^H NMR (400 MHz, DMSO-d_6_): δ = 7.58 (s, 1H), 7.22–7.10 (m, 2H), 7.08–6.88 (m, 2H), 6.83–6.74 (m, *J* = 9.1, 2.3 Hz, 2H), 3.65 (s, 3H), 3.18 (sext, *J* = 13.4, 6.7 Hz, 1H), 1.10 (d, *J* = 6.7 Hz, 6H) ppm. LC-MS (ESI+): m/z [M+H]^+^ calculated: 392.1. Found: 392.1.

### Synthesis of *tert*-butyl (2-(3-((5-(3-(difluoromethyl)-1-(4-nitrophenyl)-1*H*-pyrazol-5-yl)thiazol-2-yl)amino)-3-oxopropoxy)ethyl)carbamate (21)

3-(2-((*Tert*-butoxycarbonyl)amino)ethoxy)propanoic acid (308 mg, 1.32 mmol, 1.1 eq) was dissolved in DMF (dry, 5 mL). DIEA (627 μL, 3.60 mmol, 3.0 eq) and HATU (501 mg, 1.32 mmol, 1.1 eq) were added, and the reaction mixture was stirred for 30 min at RT. Afterward, 5-(3-(difluoromethyl)-1-(4-nitrophenyl)-1*H*-pyrazol-5-yl)thiazol-2-amine (405 mg, 1.20 mmol, 1 eq) was added, and the mixture was stirred for 16 h at RT. The solvent was removed under reduced pressure. The residue was dissolved in EtOAc (50 mL) and washed three times with NaHCO_3_ (aq., sat., 50 mL). The organic phase was dried over MgSO_4_, filtered, and the solvent was removed under reduced pressure. The crude product was purified by flash chromatography on silica (DCM → DCM/MeOH 9:1). The product was obtained as a yellow oil. Yield: 602 mg (1.09 mmol, 90%). ^1^H NMR (400 MHz, DMSO-d_6_): δ = 12.40 (s, 1H), 8.36 (d, *J* = 9.0 Hz, 2H), 7.75 (d, *J* = 9.0 Hz, 2H), 7.55 (s, 1H), 7.15 (m, 1H), 7.10 (s, 1H), 6.72 (t, *J* = 5.5 Hz, 1H), 3.65 (t, *J* = 6.1 Hz, 2H), 3.37 – 3.34 (m, *J* = 6.2 Hz, 2H), 3.02 (q, *J* = 12.1, 6.1 Hz, 2H), 2.67 (t, *J* = 5.7 Hz, 2H), 1.35 (s, 9H) ppm. LC-MS (ESI+): m/z [M+H]^+^ calculated: 553.2. Found: 553.2.

### Synthesis of 3-(2-aminoethoxy)-*N*-(5-(3-(difluoromethyl)-1-(4-nitrophenyl)-1*H*-pyrazol-5-yl)thiazol-2-yl)propanamide (22)

*tert*-Butyl (2-(3-((5-(3-(difluoromethyl)-1-(4-nitrophenyl)-1*H*-pyrazol-5-yl)thiazol-2-yl)amino)-3-oxopropoxy)ethyl)carbamate (600 mg, 1.08 mmol, 1 eq) was dissolved in DCM (5 mL) and TFA (5 mL) was added. The mixture was stirred for 1 h at RT. The solvent was removed under reduced pressure, and the residue was dissolved in DCM (100 mL). The organic phase was dried over MgSO_4_, filtered, and the solvent was removed under reduced pressure. The product was obtained as a yellow solid without any further purification. Yield: 434 mg (0.96 mmol, 88%). ^1^H NMR (400 MHz, DMSO-d_6_): δ = 8.37 – 8.32 (m, 2H), 7.75 (d, *J* = 9.0 Hz, 2H), 7.51 (s, 1H), 7.14 (m, 1H), 7.06 (s, 1H), 3.67 (t, *J* = 6.0 Hz, 2H), 3.41 (t, *J* = 5.6 Hz, 2H), 2.73 (t, *J* = 5.6 Hz, 2H), 2.64 (t, *J* = 6.0 Hz, 2H) ppm. LC-MS (ESI+): m/z [M+H]^+^ calculated: 453.1. Found: 453.1.

### Synthesis of *N*-(2-(3-((5-(3-(difluoromethyl)-1-(4-nitrophenyl)-1*H*-pyrazol-5-yl)thiazol-2-yl)amino)-3-oxopropoxy)ethyl)-5-((3*aS*,4*S*,6*aR*)-2-oxohexahydro-1*H*-thieno[3,4-*d*]imidazol-4-yl)pentanamide (23)

3-(2-Aminoethoxy)-*N*-(5-(3-(difluoromethyl)-1-(4-nitrophenyl)-1*H*-pyrazol-5-yl)thiazol-2-yl)propanamide (420 mg, 928 μmol, 1 eq) was dissolved in DMF (dry, 5 mL), DIEA (178 μL, 1.02 mmol, 1.1 eq), and 2,5-dioxopyrrolidin-1-yl 5-((3*aS*,4*S*,6*aR*)-2-oxohexahydro-1*H*-thieno[3,4-*d*]imidazol-4-yl)pentanoate (349 mg, 1.02 mmol, 1.1 eq) were added. The reaction was stirred for 1 h at RT. The solvent was removed under reduced pressure, and the residue was dissolved in DCM (100 mL). The organic phase was dried over MgSO_4_, filtered, and the solvent was removed under reduced pressure. The crude product was purified by flash chromatography on silica (DCM → DCM/MeOH 9:1). The product was obtained as a yellow solid without any further purification. Yield: 497 mg (732 µmol, 79%). ^1^H NMR (400 MHz, DMSO-d_6_): δ = 12.37 (s, 1H), 8.36 (d, *J* = 9.0 Hz, 2H), 7.82–7.72 (m, *J* = 8.1 Hz, 3H), 7.54 (s, 1H), 7.28–7.01 (m, 1H), 7.10 (s, 1H), 7.01, 6.38 (d, *J* = 21.8 Hz, 2H), 4.35–4.25 (m, 1H), 4.18–4.06 (m, 1H), 3.67 (t, *J* = 6.1 Hz, 2H), 3.37 (t, *J* = 5.9 Hz, 2H), 3.15 (q, *J* = 5.7 Hz, 2H), 3.13–3.01 (m, 1H), 2.80 (dd, *J* = 12.4, 5.0 Hz, 1H), 2.68 (t, *J* = 6.1 Hz, 2H), 2.56 (d, *J* = 12.4 Hz, 1H), 2.03 (t, *J* = 7.4 Hz, 2H), 1.66 – 1.54 (m, 1H), 1.53 – 1.40 (m, 3H), 1.36 – 1.18 (m, 2H) ppm. LC-MS (ESI+): m/z [M+H]^+^ calculated: 679.3. Found: 679.3.

### Synthesis of *N*-(2-(3-((5-(1-(4-aminophenyl)-3-(difluoromethyl)-1*H*-pyrazol-5-yl)thiazol-2-yl)amino)-3-oxopropoxy)ethyl)-5-((3*aS*,4*S*,6*aR*)-2-oxohexahydro-1*H*-thieno[3,4-*d*]imidazol-4-yl)pentanamide (24)

*N*-(2-(3-((5-(3-(difluoromethyl)-1-(4-nitrophenyl)-1*H*-pyrazol-5-yl)thiazol-2-yl)amino)-3-oxopropoxy)ethyl)-5-((3*aS*,4*S*,6*aR*)-2-oxohexahydro-1*H*-thieno[3,4-*d*]imidazol-4-yl)pentanamide (200 mg, 295 μmol, 1 eq), zinc dust (70 mg, 884 μmol, 4 eq), and acetic acid (10 mL) were suspended in MeOH (5 mL). The mixture was stirred for 3 h at RT. The mixture was filtered, and the solvent was removed under reduced pressure. The crude product was purified via RP-flash chromatography (95% H_2_O/ACN → 100% ACN), and the product was obtained as a yellowish solid. Yield: 80 mg (123 µmol, 42%). ^1^H NMR (400 MHz, DMSO-d_6_): δ = 12.24 (s, 1H), 7.76 (t, *J* = 5.4 Hz, 1H), 7.55 (s, 1H), 7.19–6.88 (m, 1H), 7.03 (s, 2H), 6.97 (s, 1H), 6.62 (d, *J* = 8.6 Hz, 2H), 6.38 (d, *J* = 22.0 Hz, 2H), 5.56 (s, 2H), 4.33–4.25 (m, 1H), 4.15–4.04 (m, 1H), 3.66 (t, *J* = 6.2 Hz, 2H), 3.36 (t, *J* = 5.8 Hz, 2H), 3.15 (q, *J* = 5.6 Hz, 2H), 3.10–2.99 (m, 1H), 2.80 (dd, *J* = 12.5, 5.0 Hz, 1H), 2.65 (t, *J* = 6.1 Hz, 2H), 2.56 (d, *J* = 12.4 Hz, 1H), 2.03 (t, *J* = 7.4 Hz, 2H), 1.64–1.53 (m, 1H), 1.50–1.39 (m, 3H), 1.31–1.20 (m, *J* = 14.5, 7.7 Hz, 2H) ppm. LC-MS (ESI+): m/z [M+H]^+^ calculated: 649.3. Found: 649.3.

### Biological and Biochemical Methods

#### LIMK Protein Expression/Purification

LIMK1, LIMK1-C349A, and LIMK2 expression constructs tagged with an N-terminal TEV cleavable 6xHis-Z-tag were expressed in insect cells after baculoviral transfection and purified as described previously.^37^ In brief, exponentially growing cells (2×10 cells/mL, Novagen) cultured in serum-free Insect-Xpress Medium (Lonza) were infected with recombinant baculovirus stock and incubated for 66 h, shaking at 27 °C. Cells were harvested by centrifugation. For lysis, cells were resuspended in lysis buffer (50 mM HEPES pH 7.4, 500 mM NaCl, 20 mM imidazole, 0.5 mM TCEP, 5% glycerol) and sonicated. After clarification, the lysate was loaded onto pre-equilibrated Ni NTA Sepharose beads (Quiagen). After a stringent wash with lysis buffer, the His6-tagged proteins were eluted in lysis buffer supplemented with 300 mM imidazole. After reducing the NaCl concentration to 250 mM, the eluate was loaded onto a SP Sepharose column (Cytiva) and eluted by applying a salt gradient ranging from 250 mM to 2.5 M. Fractions containing LIMK1 were pooled, and the N-terminal tag was cleaved by TEV protease overnight. Contaminating proteins, the cleaved tags, and TEV protease were removed with a sequential reverse SP Sepharose Ni NTA column. The LIMK1 protein was concentrated and subjected to gel filtration using an AKTA Xpress system combined with an S200 16/600 gel filtration column (GE Healthcare) in 20 mM HEPES (pH 7.4), 150 mM NaCl, 0.5 mM TCEP, 5% glycerol, and 2.5 mM MgCl_2_. The elution volume of 92 mL indicated LIMK1 to be monomeric in solution. The final yield for LIMK1 330-637 was 0.2 mg/L of insect cell medium.

##### Expression Constructs

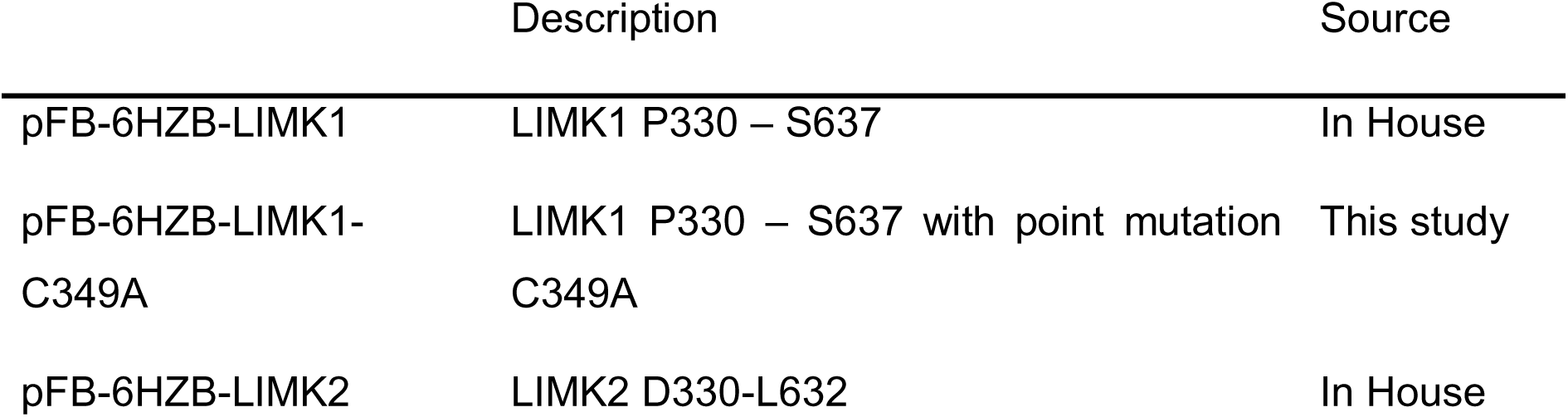

LIMK1/C349A mutation was created by quick change following QuikChange Site-directed mutagenesis kit instructions (Agilent) on pFB-6HZB-LIMK1. Oligonucleotides were ordered from Merck; forward: CTGGGCAAGGGCGCCTTCGGCCAGGC and reverse: GCCTGGCCGAAGGCGCCCTTGCCCAG. For PCR amplification, Herculase II (Agilent, 600675) was used. Successful nucleotide exchange was verified by sequencing at MircoSynth Seqlab.

#### Differential Scanning Fluorimetry (DSF)

Thermal stabilization of LIMK1/2 proteins was measured by DSF assays according to previously established protocols.^38^ Briefly, 2 μM LIMK protein in assay buffer (20 mM HEPES pH 7.4, 150 mM NaCl, 0.5 mM TCEP, 5% glycerol) was mixed with a 1:1000 dilution of SYPRO Orange (Sigma-Aldrich). A sample of 20 µL was added to each well of a 96-well plate (Starlab, white). The respective compound was added with a final concentration of 10 µM. Fluorescence was monitored in an MX3005P real-time PCR instrument (Stratagene) with excitation and emission filters set at 465 and 590 nm while gradually heating (1 °C per min) from 25 to 95°C. Data were analyzed with the MxPro3005 software (Stratagene). Fluorescence was plotted against the temperature and the melting point (*T*_m_) was determined as the minimum of the first derivative relative to the control.

#### DSF-based selectivity screening against a curated kinase library

The assay was performed as previously described. Briefly, recombinant protein kinase domains at a concentration of 2 μM were mixed with 10 μM compound in a buffer containing 20 mM HEPES, pH 7.5, and 500 mM NaCl. SYPRO Orange (5000×, Invitrogen) was added as a fluorescence probe (1 µL per mL). Subsequently, temperature-dependent protein unfolding profiles were measured using the QuantStudio™ 5 real-time PCR machine (Thermo Fisher). Excitation and emission filters were set to 465 nm and 590 nm, respectively. The temperature was raised at a step rate of 3°C per minute. Data points were analyzed with the internal software (Thermal Shift Software^TM^ Version 1.4, Thermo Fisher) using the Boltzmann equation to determine the inflection point of the transition curve.^39,40^

#### Mass spectrometry with LIMK1 wt and mutant

LIMK1 wt, or LIMK1/C349A variant protein (5 µM), was incubated with 10 µM of the compound at room temperature for the indicated time. The modification reaction was stopped by adding an equal volume of MS buffer (0.1% formic acid in dH_2_O). The reaction mixture (5 µL) was injected into an Agilent 6230 Electrospray Ionization Time-of-Flight mass spectrometer coupled to a 1260 Infinity liquid chromatography unit (0.4 mL/min flow rate using a solvent gradient of water to acetonitrile with 0.1% formic acid). Data was acquired using the MassHunter LC/MS Data Acquisition software (Agilent Technology) and analyzed using the BioConfirm vB.08.00 tool (Agilent Technology). Peak intensities of modified and non-modified LIMK1 were quantified, and the ratio was calculated

#### High-Throughput Kinetic Screening Assay for Reversible Compounds

LIMK1 KINETICfinder^®^ assay (Enzymlogic) was based on the binding and displacement of a fluorescent probe to the ATP-binding site of the kinase with TR-FRET detection using terbium-labeled antibodies. The assays were performed in black 384 well microplates containing 0.1 nM of LIMK1 (Carna Biosciences), 30 nM of fluorescent probe and 2 nM of Tb-labeled antibody (Life Technologies) in assay buffer (50 mM HEPES, pH 7.5, 10 mM MgCl_2_, 0.01% Brij-35, 1 mM DTT and 1% DMSO). For all experiments, a 4-point 10-fold serial dilution of 100X concentrated test compounds was prepared in DMSO. The kinetic assays were read continuously at room temperature in a PHERAstar FSX plate reader (BMG LABTECH), and the specific signals were fitted to Motulsky-Mahan’s equation. The affinity (K_d_), association rate constant (*k_on_*), dissociation rate constant (*k_off_*), and residence time (T) of each test compound were calculated using KINPy^®^ software (Enzymlogic).

#### High-Throughput Kinetic Screening Assay for Irreversible Compounds

LIMK1 COVALfinder^®^ (Enzymlogic) is a TR-FRET binding assay that contains 0.1 nM of LIMK1, 30 nM fluorescent probe, and 2 nM terbium antibody in assay buffer. For all experiments, 100x concentrated test compounds were serially diluted (1.75-fold, 20 points) in DMSO. Kinetic measurements were continuously recorded at room temperature, and the specific signals were fitted to a single-exponential equation to determine k_obs_, the apparent first-order rate constant for the interconversion between the initial and final degree of binding. A secondary plot of k_obs_ versus compound concentration enabled the calculation of kinetic constants and inhibition mechanism. In Equation 1, *k_inact_* represents the maximum inactivation rate at infinite compound concentration, while *K*_I_ denotes the concentration required to achieve half-maximal inactivation. Inactivation efficiency is measured by the second-order rate constant *k_inact_*/*K*_I,_ and the half-life for inactivation at this infinite concentration is given by T_1/2_= 0.693/k_inact_. Additionally, dose– response curves were generated at each time point to calculate IC₅₀ values, which were then plotted over time to facilitate the inspection of time dependency.

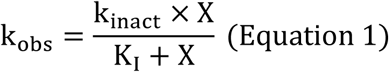

#### NanoBRET cellular target engagement assay

The LIMK1 and LIMK2 NanoBRET assays were performed as described before.^41^ In brief, full-length LIMK1 and LIMK2 (Promega) cloned in frame with a C-terminal NanoLuc-fusion were transfected into HEK293T cells, and proteins were allowed to express for 20 h. For the target engagement assay, serially diluted inhibitor and NanoBRET Kinase Tracer K10 (Promega) at 325 nM for LIMK1 and 375 nM for LIMK2, respectively, were pipetted into white 384-well plates (Greiner 781 207). The corresponding LIMK1 or LIMK2-transfected cells were added and reseeded at a density of 2 × 10^5^ cells/mL after trypsinization and resuspending in Opti-MEM without phenol red (Life Technologies). The system was allowed to equilibrate for 2 hours at 37°C/5% CO_2_ before BRET measurements. To measure BRET, NanoBRET NanoGlo Substrate + Extracellular NanoLuc Inhibitor (Promega) was added as per the manufacturer’s protocol, and filtered luminescence was measured on a PHERAstar FSX plate reader (BMG Labtech) equipped with a 450 nm BP filter (donor) and a 610 nm LP filter (acceptor). Competitive displacement data were then graphed using GraphPad Prism software, applying a 4-parameter curve fit with the following equation: Y=Bottom + (Top-Bottom) / (1+10^((LogIC50-X)*HillSlope))

#### NanoBRET cellular kinetic assay

The kinetic NanoBRET wash-out assays were performed as described before.^30^ The inhibitor at a concentration of 10 times its previously determined EC_50_ was pipetted into white 384-well plates (Greiner 781 207) and the corresponding LIMK1 or LIMK2-transfected cells were added and reseeded at a density of 2 × 10^5^ cells/mL after trypsinization and resuspending in Opti-MEM without phenol red (Life Technologies). The system was allowed to equilibrate for 2 hours at 37°C/5% CO_2_ to reach approximately 90% target occupancy before the wash-out via removal of the medium to extract any unbound inhibitor, washing with Opti-MEM, and re-addition of the removed volume of Opti-MEM medium. The tracer was added before BRET measurements at a final concentration as described for the target engagement assays. To measure BRET, NanoBRET NanoGlo Substrate + Extracellular NanoLuc Inhibitor (Promega) was added as per the manufacturer’s protocol, and filtered luminescence was measured on a PHERAstar plate reader (BMG Labtech) equipped with a 450 nm BP filter (donor) and 610 nm LP filter (acceptor) for 2h. The preincubated and saturated inhibitor-target complex dissociates during the wash-out, and the tracer added associates, resulting in an increasing BRET signal. The background of the kinetic data was corrected and then graphed using GraphPad Prism software using a two-phase association fit with the following equation: SpanFast=(Plateau-Y0)*PercentFast*.01; SpanSlow=(Plateau-Y0)*(100-PercentFast)*.01; Y=Y0+ SpanFast*(1-exp(-KFast*X)) + SpanSlow*(1-exp(-KSlow*X)). The half-life of the slow component fit is reported.

#### NanoBRET-K192 panel

Compound selectivity inside cells was assessed by using the K192 Kinase Selectivity System (Promega, cat. no. NP4050). For plate preparation, the transfection mix was prepared in white 384-well small-volume plates (Greiner, cat. no. 784075) by pre-plating 3 µL of 20 µL/mL FuGene HD (Promega, cat. no.. E2311), diluted in optiMEM medium (Gibco, cat. no. 11058-021). DNA from both DNA vector source plates (1 µL) of the K192 kit was added using an Echo acoustic dispenser (Beckman Coulter). The mix was incubated for 30 min, and HEK293T cells in optiMEM medium (6 µL) were added. The proteins were allowed to be expressed for 20 h. After expression, Tracer K10 was added using the concentrations recommended in the K192 technical manual, and 1 µM inhibitor was added to every second well. After 2 h of equilibration, detection was carried out using a substrate solution comprising optiMEM with a 1:166 dilution of NanoBRET™ Nano-Glo® Substrate and a 1:500 dilution of the Extracellular NanoLuc® Inhibitor. The substrate solution (5 µL) was added to every well, and filtered luminescence was measured on a PHERAstar plate reader (BMG Labtech) equipped with a luminescence filter pair (450 nm BP filter (donor) and 610 nm LP filter (acceptor)). For every kinase, occupancy was calculated and plotted using GraphPad Prism 10.

#### SPR experiment

The SPR analysis was performed on a Biacore T200 (Cytiva Life Sciences). Approximately 7000 RU of biotinylated LIMK1/C349A variant was loaded onto a Series S CM5 chip coated with Streptavidin. The chip was equilibrated with a running buffer containing 20 mM HEPES pH 7.4, 150 mM NaCl, 0.5 mM TCEP, and 0.05% Tween 20. A titration of serially diluted compounds **10** was performed. The compounds were allowed to bind over the surface at a flow rate of 30 μL/min for 60 seconds, followed by disassociation wash for 180 seconds. The sensorgrams were double-reference subtracted and analyzed by a steady-state affinity fit model.

#### GSH assay

The GSH assay for the final compounds was conducted as described earlier^42^ using HPLC analysis with an Agilent 1260 Infinity II system, equipped with a 1260 DAD HS detector (G7117C) set at 254 nm, 280 nm, and 310 nm. Separation was performed on a Poroshell 120 EC-C18 reversed-phase column (Agilent, 3 × 150 mm, 2.7 μm) using a gradient elution method with 0.1% formic acid in water (solvent A) and 0.1% formic acid in acetonitrile (solvent B) as the mobile phase. The assay was carried out in a reaction mixture consisting of 1.8 mL of degassed potassium phosphate buffer (1M, pH 7.4) and 200 µL of acetonitrile at 40 °C. The compound of interest was used at a concentration of 50 µM, while reduced GSH was added in a 100-fold excess (5 mM). To assess system stability, a UV-active internal standard (e.g., BIX 02189) was included at a concentration of 50 µM. Before the addition of GSH, the UV absorbance of a mixture of the covalent compound and the internal standard was recorded. Following GSH addition, 60 µL aliquots were collected at predefined time points. A time-dependent decrease in the UV absorbance of the covalent compound was observed, while the absorbance of the internal standard remained constant. The half-life (t_1/2_) of the reaction with GSH was then calculated using the equations outlined in an earlier work.^43^

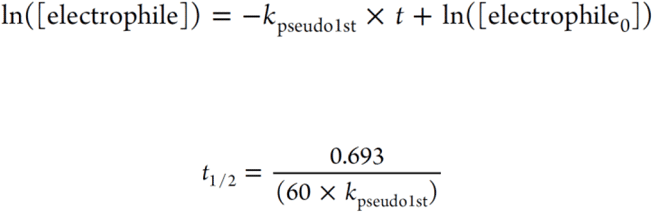

#### Chemoproteomics experiments

HEK293 cells (ATCC CRL-1573) were grown to confluency in 15 cm dishes. The cells were detached with 10 mL phosphate-buffered saline (1x PBS) containing 1 mM EDTA. After washing the cells once with 1x PBS (1 mM EDTA) and removing the supernatant, they were snap-frozen in liquid nitrogen and stored at −80°C. Cells were lysed by resuspension in HNN lysis buffer (50 mM HEPES pH 8.0, 150 mM NaCl, 50 mM NaF, 0.5% IGPAL-630, 400 nM Na_3_VO_4_, 1 mM PMSF, and 0.2% protease inhibitors (Sigma). Following 10 minutes of incubation on ice, the lysate was centrifuged for 30 minutes at 18.000g. After pooling the cleared lysate, it was distributed into 1900 µL aliquots per sample (equivalent to 2×15 cm dishes). The compound **10**, (50 µM) or DMSO was added and pre-incubated for 1 hour at 4°C before the addition of compound SM429 (5 µM) or DMSO to the individual samples, which were then incubated for 15 min at 4°C on a rotating wheel. The experiments were carried out with n=4 replicates per condition. Next, the treated lysate was added to 80 µL of Strep-Tactin beads (50% slurry, IBA Lifesciences) previously modified to be resistant to tryptic cleavagefollowed by incubation of the samples on a rotating wheel for 1 hour at 4°C. After the transfer of the beads to a 96-well 1 µm glass filter plate (Pall corporation), the residual lysate was filtered and the beads were washed two times with HNN-lysis buffer without protease inhibitors and PMSF, twice with 1 mL of HNN lysis buffer without protease inhibitors, PMSF and IGPAL-630, and twice with 1 mL of 100 mM ammonium bicarbonate (ABC). The beads were resuspended in 100 mM ABC and transferred to a 10 kDa cutoff plate (Pall Corporation). The supernatant was removed via centrifugation at 1500g for 15 minutes. The samples were resuspended in 8M urea in 100 mM ABC, reduced with 10 mM TCEP (40 minutes, 37°C, and 200 rpm), and alkylated with 20 mM Iodoacetamide (30 minutes, 37°C and 200 rpm). Afterward, urea was removed by centrifugation at 1500g for 15 minutes, followed by two wash steps with 100 mM ABC, where the centrifugation time was increased to 30 minutes after the last wash. The beads were resuspended in 203 µL of ABC (100 mM), containing 1 µg of trypsin (Promega) and 0.5 µg of LysC (FUJIFILM Wako) for overnight digestion (37°C, 200 rpm). The supernatant containing the peptides was collected via centrifugation at 1500g for 20 minutes, and the remaining peptides were eluted in 100 µL of 100 mM ABC. Both solutions were pooled and acidified with formic acid (FA) to a final concentration of 5%. The peptides were loaded on an equilibrated 96-well C18 plate (Nest group, HNFR S18V) via centrifugation (1500g, 1 min), washed three times with 200 µL of 5% acetonitrile (ACN) with 0.1% FA and finally eluted with 2× 100 µL of 50% ACN with 0.1% FA into a fresh collection plate. Peptides were dried in the speed vac and resuspended in 20 µL of 2% ACN with 0.1% FA and 0.2x iRT standard (Biognosys). Samples were injected into a Waters nanoACQUITY coupled to an Orbitrap Fusion Lumos (injection volume 3 µL). In the samples where the intensity in the chromatogram deviated from the other replicate, the injection volume was adjusted. The peptides were separated on a 30 cm column packed with 3 µm C18 resin (Dr. Maisch) by a 120 min gradient from 3% to 35% buffer B (100% ACN containing 0.1% FA) at a flow rate of 300 nL/min. Each sample was measured in data-independent acquisition (DIA) and data-dependent acquisition (DDA) modes. The DDA mode was performed with the following parameters: the full scan range was 350 – 1’150m/z at 120’000 resolutions. The data-dependent scans were acquired within a cycle time of 3 sec. Only charge states between or equal to 2 – 7 were included. Fragmentation was obtained with an HCD collision energy of 30%. The MS2 spectra were measured with an Orbitrap resolution of 30’000 with isolation windows of 1.6 m/z. The normalized AGC target was set to 200% with a maximum injection time of 54 msec. The DIA mode was performed with the following parameters: the scan range was 350 – 1400 at 120’000 resolutions. The normalized AGC target was set to 50% with a maximum injection time of 100 msec. The RF lens was set to 30%. The targeted MS2 spectra for the desired masses in the variable isolation windows, together with the normalized AGC target percentage listed below, were acquired by fragmentation with an HCD of 28%. The Orbitrap resolution was 30’000 with variable scan ranges. The maximum injection time was set to 54 msec, and the RF lens to 30%. A hybrid spectral library was generated from all DDA and DIA runs, using the Pulsar search engine in Biognosys Spectronaut v.19.1. The standard settings were used except that LysC/P was added to the cleavage rules. The DIA runs were searched in Biognosys Spectronaut v.19.1 against the generated library with the default settings, with the following changes: Cross-Run Normalization was unselected, and Used Biognosys’ iRT Kit was selected. The precursor-level data were exported and further processed in protti version 0.9 R version 4.4.1.^44^ Data normalization was performed on the precursor level, and protein abundances were calculated for proteins with at least three observed precursors in each sample. Following differential abundance calculation, significance was determined using a moderated t-test with Benjamini-Hochberg multiple testing correction. Changes in protein abundances were considered significant if the absolute fold change (log2) was greater than one unit and the adjusted p values were below 0.05.^45^

#### Cofilin phosphorylation (Western Blot assay)

Cells Treatment: LN229 is a glioblastoma cell line and cultured until passage 17 in Dulbecco’s modified Eagle’s medium (DMEM) with GlutaMAX^TM^, 10% fetal bovine serum (FBS), and 1% sodium pyruvate (all from Invitrogen, Carlsbad, CA, USA) with 5% CO_2_ at 37°C. For the treatment, 3 × 10^5^ cells were plated and treated with DMSO as the control, or 100 nM, 500 nM, 1 µM, and 5 µM of each compound for 6 h.

Western Blot analysis: The cells were lysed using RIPA buffer (50 mM Tris HCl pH 7.4, 150 mM NaCl, 1% Triton X-100, 1% Na DOC, 0.1% SDS, 1 mM EDTA), freshly supplemented with protease and phosphatase inhibitor cocktails (Sigma-Aldrich, Darmstadt, Germany). After 1 h of incubation on ice, the lysates were centrifuged for 45 min at 250 × g, and the protein-containing supernatant was collected. The protein concentrations were determined using Protein Assay Dye Reagent Concentrate (Bradford, Bio-Rad, Hercules, CA, USA) and Pre-Diluted Protein Assay Standards: Bovine Serum Albumin (BSA) Set (Sigma, A2153, Darmstadt, Germany). Then, 10 μg of protein was mixed with Roti®-Load 1 dye (Carl Roth, Karlsruhe, Germany) and 4x Laemmli Buffer (Bio-Rad, Hercules, CA, USA), denatured, and run on NuPAGE^TM^ 4−12% bis-Tris gels (Thermo Fisher Scientific, Darmstadt, Germany). Proteins were blotted on methanol-activated PVDF Transfer Membranes (Thermo Fisher Scientific, Darmstadt, Germany) using the wet transfer method (Transfer Buffer +20% methanol). Afterward, the membranes were blocked with 5% milk or 3% BSA in 0.1% TBS-T for 1 h at room temperature and incubated with primary antibodies overnight at 4°C. After incubation with secondary horseradish-peroxidase (HRP)-conjugated antibodies for 2 h (Cell Signaling Technology, MA, USA), membranes were washed with 0.1% TBS-T and developed using an imaging instrument (Azure Biosystems, CA, USA). Band intensities were quantified using FIJI Image J software (version 2.14.0). A list of all antibodies can be found in the table below.

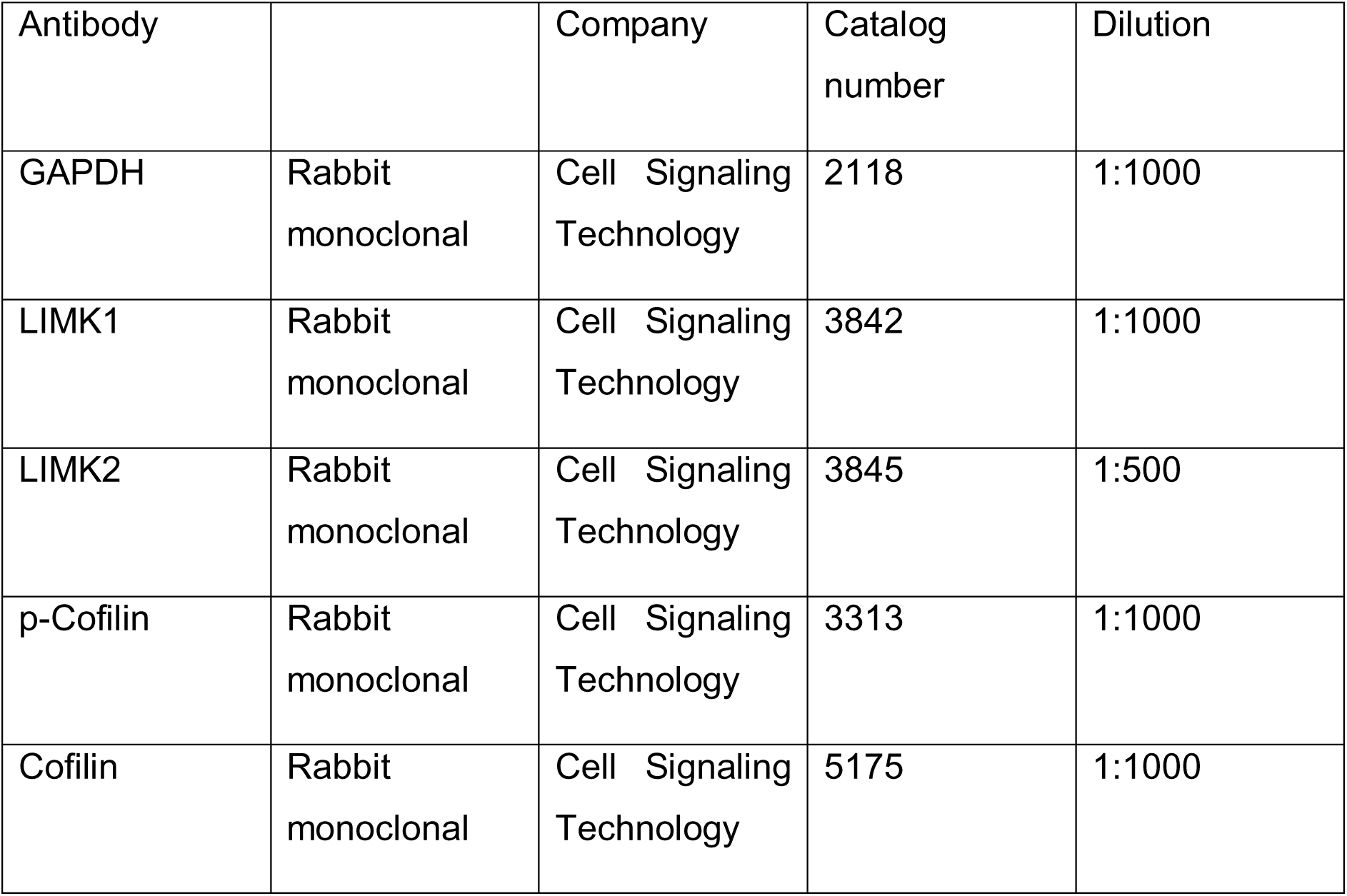

| Antibody |  | Company | Catalog number | Dilution |
| --- | --- | --- | --- | --- |
| GAPDH | Rabbit monoclonal | Cell Signaling Technology | 2118 | 1:1000 |
| LIMK1 | Rabbit monoclonal | Cell Signaling Technology | 3842 | 1:1000 |
| LIMK2 | Rabbit monoclonal | Cell Signaling Technology | 3845 | 1:500 |
| p-Cofilin | Rabbit monoclonal | Cell Signaling Technology | 3313 | 1:1000 |
| Cofilin | Rabbit monoclonal | Cell Signaling Technology | 5175 | 1:1000 |

## Supporting Information

Experimental methods, biochemical assays, compound synthesis, and analytics have been included in the Supporting information Word document. Results from kinetic profiling of compounds and molecular formula strings have been included in separate Excel files.

## Author Information

### Author Contributions

S.M. and T.H. equally contributed to the compound synthesis and experimental data. S.M., S.R., and S.K. wrote the manuscript. TR-FRET assays and kinetic studies were performed by A.C., P.A., N.P., and M.B.-N. Protein production and further biochemical assays were done by K.R.A., A.K., and V.D. Cellular assays were performed by L.-M.B., M.S., B.-T.B., L.E., H.S., and proteomics experiments by F.F. and S.K. The research was supervised by D.S.K., M.G., S.M., S.R., and S.K. All authors edited and approved the manuscript.

### Notes

The authors declare no competing financial interest.

## Acknowledgments

SM, TH, SR, L-MB, MS, B-TB, and SK are grateful for support from the Structural Genomics Consortium (SGC), a registered charity (no: 1097737) that receives funds from Bayer AG, Boehringer Ingelheim, Bristol Myers Squibb, Genentech, Genome Canada through Ontario Genomics Institute, EU/EFPIA/OICR/McGill/KTH/Diamond Innovative Medicines Initiative 2 Joint Undertaking [EUbOPEN grant 875510], Janssen, Pfizer, and Takeda. S.K. is also funded by the German Cancer Research Center (DKTK), the Frankfurt Cancer Institute (FCI). DSK and SK would like to acknowledge funding by the BMBF Cluster for the future program PROXIDRUGS. MPS is funded by the Deutsche Forschungsgemeinschaft CRC1430 (Project-ID 424228829). And SM and SK are also funded by the German Cancer Aid (Krebshilfe), a large pre-clinical program, TACTIC.

## Abbreviations

AAK1: AP2 Associated Kinase 1
ABHD5: Abhydrolase Domain Containing 5
BMP: Bone morphogenetic protein
BMPR2: Bone morphogenetic protein type 2 receptor
CDKL2: Cyclin Dependent Kinase Like 2
DCAF11: DDB1 and CUL4 Associated Factor 11
DIEA: N,N-Diisopropylethylamine
DMF-DMA: N,N-Dimethylformamiddimethylacetal
ESI-TOF: Electrospray ionization time-of-flight
FXR: Fragile X syndrome
LIMK: LIM Domain Kinase
LTP: Long-term potentiation
NLK: Nemo Like Kinase
JAK: Janus Kinase
JNK1-3: Mitogen-Activated Protein Kinase 8-10
SRC: SRC Proto-Oncogene, Non-Receptor Tyrosine Kinase
TCDI: 1,1’-Thiocarbonyldiimidazole
TFA: Trifluoroacetic acid
TXK: TXK Tyrosine Kinase

## References

(1) Chatterjee, D.; Preuss, F.; Dederer, V.; Knapp, S.; Mathea, S. Structural Aspects of LIMK Regulation and Pharmacology. Cells 2022, 11 (1), 142. 10.3390/cells11010142.

(2) Pröschel, C.; Blouin, M. J.; Gutowski, N. J.; Ludwig, R.; Noble, M. Limk1 Is Predominantly Expressed in Neural Tissues and Phosphorylates Serine, Threonine and Tyrosine Residues in Vitro. Oncogene 1995, 11 (7), 1271–1281.

(3) Manetti, F. LIM Kinases Are Attractive Targets with Many Macromolecular Partners and Only a Few Small Molecule Regulators. Med. Res. Rev. 2012, 32 (5), 968–998. 10.1002/med.20230.

(4) Berabez, R.; Routier, S.; Bénédetti, H.; Plé, K.; Vallée, B. LIM Kinases, Promising but Reluctant Therapeutic Targets: Chemistry and Preclinical Validation In Vivo. Cells 2022, 11 (13), 2090. 10.3390/cells11132090.

(5) Ben Zablah, Y.; Zhang, H.; Gugustea, R.; Jia, Z. LIM-Kinases in Synaptic Plasticity, Memory, and Brain Diseases. Cells 2021, 10 (8), 2079. 10.3390/cells10082079.

(6) Prunier, C.; Prudent, R.; Kapur, R.; Sadoul, K.; Lafanechère, L. LIM Kinases: Cofilin and Beyond. Oncotarget 2017, 8 (25), 41749–41763. 10.18632/oncotarget.16978.

(7) Sumi, T.; Matsumoto, K.; Nakamura, T. Specific Activation of LIM Kinase 2 via Phosphorylation of Threonine 505 by ROCK, a Rho-Dependent Protein Kinase. J. Biol. Chem. 2001, 276 (1), 670–676. 10.1074/jbc.M007074200.

(8) Amano, T.; Tanabe, K.; Eto, T.; Narumiya, S.; Mizuno, K. LIM-Kinase 2 Induces Formation of Stress FIbres, Focal Adhesions and Membrane Blebs, Dependent on Its Activation by Rho-Associated Kinase-Catalysed Phosphorylation at Threonine-505. 2001.

(9) Bernard, O. Lim Kinases, Regulators of Actin Dynamics. Int. J. Biochem. Cell Biol. 2007, 39 (6), 1071–1076. 10.1016/j.biocel.2006.11.011.

(10) Wufuer, R.; Ma, H.-X.; Luo, M.-Y.; Xu, K.-Y.; Kang, L. Downregulation of Rac1/PAK1/LIMK1/Cofilin Signaling Pathway in Colon Cancer SW620 Cells Treated with Chlorin E6 Photodynamic Therapy. Photodiagnosis Photodyn. Ther. 2021, 33, 102143. 10.1016/j.pdpdt.2020.102143.

(11) Tavares, S.; Vieira, A. F.; Taubenberger, A. V.; Araújo, M.; Martins, N. P.; Brás-Pereira, C.; Polónia, A.; Herbig, M.; Barreto, C.; Otto, O.; Cardoso, J.; Pereira-Leal, J. B.; Guck, J.; Paredes, J.; Janody, F. Actin Stress Fiber Organization Promotes Cell Stiffening and Proliferation of Pre-Invasive Breast Cancer Cells. Nat. Commun. 2017, 8, 15237. 10.1038/ncomms15237.

(12) Harrison, B. A.; Almstead, Z. Y.; Burgoon, H.; Gardyan, M.; Goodwin, N. C.; Healy, J.; Liu, Y.; Mabon, R.; Marinelli, B.; Samala, L.; Zhang, Y.; Stouch, T. R.; Whitlock, N. A.; Gopinathan, S.; McKnight, B.; Wang, S.; Patel, N.; Wilson, A. G. E.; Hamman, B. D.; Rice, D. S.; Rawlins, D. B. Discovery and Development of LX7101, a Dual LIM-Kinase and ROCK Inhibitor for the Treatment of Glaucoma. ACS Med. Chem. Lett. 2015, 6 (1), 84–88. 10.1021/ml500367g.

(13) Meng, Y.; Zhang, Y.; Tregoubov, V.; Janus, C.; Cruz, L.; Jackson, M.; Lu, W. Y.; MacDonald, J. F.; Wang, J. Y.; Falls, D. L.; Jia, Z. Abnormal Spine Morphology and Enhanced LTP in LIMK-1 Knockout Mice. Neuron 2002, 35 (1), 121–133. 10.1016/s0896-6273(02)00758-4.

(14) Rust, M. B.; Gurniak, C. B.; Renner, M.; Vara, H.; Morando, L.; Görlich, A.; Sassoè-Pognetto, M.; Banchaabouchi, M. A.; Giustetto, M.; Triller, A.; Choquet, D.; Witke, W. Learning, AMPA Receptor Mobility and Synaptic Plasticity Depend on n-Cofilin-Mediated Actin Dynamics. EMBO J. 2010, 29 (11), 1889–1902. 10.1038/emboj.2010.72.

(15) Kashima, R.; Redmond, P. L.; Ghatpande, P.; Roy, S.; Kornberg, T. B.; Hanke, T.; Knapp, S.; Lagna, G.; Hata, A. Hyperactive Locomotion in a *Drosophila* Model Is a Functional Readout for the Synaptic Abnormalities Underlying Fragile X Syndrome. Sci. Signal. 2017, 10 (477), eaai8133. 10.1126/scisignal.aai8133.

(16) Kashima, R.; Roy, S.; Ascano, M.; Martinez-Cerdeno, V.; Ariza-Torres, J.; Kim, S.; Louie, J.; Lu, Y.; Leyton, P.; Bloch, K. D.; Kornberg, T. B.; Hagerman, P. J.; Hagerman, R.; Lagna, G.; Hata, A. Augmented Noncanonical BMP Type II Receptor Signaling Mediates the Synaptic Abnormality of Fragile X Syndrome. Sci. Signal. 2016, 9 (431). 10.1126/scisignal.aaf6060.

(17) Chaikuad, A.; Koch, P.; Laufer, S. A.; Knapp, S. The Cysteinome of Protein Kinases as a Target in Drug Development. Angew. Chem. Int. Ed. 2018, 57 (16), 4372–4385. 10.1002/anie.201707875.

(18) Forster, M.; Chaikuad, A.; Bauer, S. M.; Holstein, J.; Robers, M. B.; Corona, C. R.; Gehringer, M.; Pfaffenrot, E.; Ghoreschi, K.; Knapp, S.; Laufer, S. A. Selective JAK3 Inhibitors with a Covalent Reversible Binding Mode Targeting a New Induced Fit Binding Pocket. Cell Chem. Biol. 2016, 23 (11), 1335–1340. 10.1016/j.chembiol.2016.10.008.

(19) Shi, L.; Zhong, Z.; Li, X.; Zhou, Y.; Pan, Z. Discovery of an Orally Available Janus Kinase 3 Selective Covalent Inhibitor. J. Med. Chem. 2019, 62 (2), 1054–1066. 10.1021/acs.jmedchem.8b01823.

(20) Kempson, J.; Ovalle, D.; Guo, J.; Wrobleski, S. T.; Lin, S.; Spergel, S. H.; Duan, J. J.-W.; Jiang, B.; Lu, Z.; Das, J.; Yang, B. V.; Hynes, J.; Wu, H.; Tokarski, J.; Sack, J. S.; Khan, J.; Schieven, G.; Blatt, Y.; Chaudhry, C.; Salter-Cid, L. M.; Fura, A.; Barrish, J. C.; Carter, P. H.; Pitts, W. J. Discovery of Highly Potent, Selective, Covalent Inhibitors of JAK3. Bioorg. Med. Chem. Lett. 2017, 27 (20), 4622–4625. 10.1016/j.bmcl.2017.09.023.

(21) Kung, A.; Chen, Y.-C.; Schimpl, M.; Ni, F.; Zhu, J.; Turner, M.; Molina, H.; Overman, R.; Zhang, C. Development of Specific, Irreversible Inhibitors for a Receptor Tyrosine Kinase EphB3. J. Am. Chem. Soc. 2016, 138 (33), 10554–10560. 10.1021/jacs.6b05483.

(22) Hanke, T.; Mathea, S.; Woortman, J.; Salah, E.; Berger, B.-T.; Tumber, A.; Kashima, R.; Hata, A.; Kuster, B.; Müller, S.; Knapp, S. Development and Characterization of Type I, Type II, and Type III LIM-Kinase Chemical Probes. J. Med. Chem. 2022, 65 (19), 13264–13287. 10.1021/acs.jmedchem.2c01106.

(23) Ross-Macdonald, P.; de Silva, H.; Guo, Q.; Xiao, H.; Hung, C.-Y.; Penhallow, B.; Markwalder, J.; He, L.; Attar, R. M.; Lin, T.; Seitz, S.; Tilford, C.; Wardwell-Swanson, J.; Jackson, D. Identification of a Nonkinase Target Mediating Cytotoxicity of Novel Kinase Inhibitors. Mol. Cancer Ther. 2008, 7 (11), 3490–3498. 10.1158/1535-7163.MCT-08-0826.

(24) Medina, C.; De La Fuente, V.; Tom Dieck, S.; Nassim-Assir, B.; Dalmay, T.; Bartnik, I.; Lunardi, P.; De Oliveira Alvares, L.; Schuman, E. M.; Letzkus, J. J.; Romano, A. LIMK, Cofilin 1 and Actin Dynamics Involvement in Fear Memory Processing. Neurobiol. Learn. Mem. 2020, 173, 107275. 10.1016/j.nlm.2020.107275.

(25) Harrison, B. A.; Whitlock, N. A.; Voronkov, M. V.; Almstead, Z. Y.; Gu, K.; Mabon, R.; Gardyan, M.; Hamman, B. D.; Allen, J.; Gopinathan, S.; McKnight, B.; Crist, M.; Zhang, Y.; Liu, Y.; Courtney, L. F.; Key, B.; Zhou, J.; Patel, N.; Yates, P. W.; Liu, Q.; Wilson, A. G. E.; Kimball, S. D.; Crosson, C. E.; Rice, D. S.; Rawlins, D. B. Novel Class of LIM-Kinase 2 Inhibitors for the Treatment of Ocular Hypertension and Associated Glaucoma. J. Med. Chem. 2009, 52 (21), 6515–6518. 10.1021/jm901226j.

(26) Mandel, S.; Hanke, T.; Mathea, S.; Chatterjee, D.; Saraswati, H.; Berger, B.-T.; Schwalm, M.-P.; Yamamoto, S.; Tawada, M.; Takagi, T.; Röhm, S.; Corrionero, A.; Alfonso, P.; Baena, M.; Elson, L.; Menge, A.; Krämer, A.; Pereira, R.; Müller, S.; Krause, D. S.; Knapp, S. Repurposing of the RIPK1 Selective Benzo[1,4]Oxazepin-4-One Scaffold for the Development of a Type-III LIMK1/2 Inhibitor. February 3, 2025. 10.1101/2025.02.03.636296.

(27) Baldwin, A. G.; Foley, D. W.; Collins, R.; Lee, H.; Jones, D. H.; Wahab, B.; Waters, L.; Pedder, J.; Paine, M.; Feng, G. J.; Privitera, L.; Ashall-Kelly, A.; Thomas, C.; Gillespie, J. A.; Schino, L.;Belelli, D.; Rocha, C.; Maussion, G.; Krahn, A. I.; Durcan, T. M.; Elkins, J. M.; Lambert, J. J.; Atack, J. R.; Ward, S. E. Discovery of MDI-114215: A Potent and Selective LIMK Inhibitor To Treat Fragile X Syndrome. J. Med. Chem. 2025, 68 (1), 719–752. 10.1021/acs.jmedchem.4c02694.

(28) Rao, S.; Gurbani, D.; Du, G.; Everley, R. A.; Browne, C. M.; Chaikuad, A.; Tan, L.; Schröder, M.; Gondi, S.; Ficarro, S. B.; Sim, T.; Kim, N. D.; Berberich, M. J.; Knapp, S.; Marto, J. A.; Westover, K. D.; Sorger, P. K.; Gray, N. S. Leveraging Compound Promiscuity to Identify Targetable Cysteines within the Kinome. Cell Chem. Biol. 2019, 26 (6), 818–829.e9. 10.1016/j.chembiol.2019.02.021.

(29) Heroven, C.; Georgi, V.; Ganotra, G. K.; Brennan, P.; Wolfreys, F.; Wade, R. C.; Fernández-Montalván, A. E.; Chaikuad, A.; Knapp, S. Halogen-Aromatic π Interactions Modulate Inhibitor Residence Times. Angew. Chem. Int. Ed Engl. 2018, 57 (24), 7220–7224. 10.1002/anie.201801666.

(30) Berger, B.-T.; Amaral, M.; Kokh, D. B.; Nunes-Alves, A.; Musil, D.; Heinrich, T.; Schröder, M.; Neil, R.; Wang, J.; Navratilova, I.; Bomke, J.; Elkins, J. M.; Müller, S.; Frech, M.; Wade, R. C.; Knapp, S. Structure-Kinetic Relationship Reveals the Mechanism of Selectivity of FAK Inhibitors over PYK2. Cell Chem. Biol. 2021, 28 (5), 686–698.e7. 10.1016/j.chembiol.2021.01.003.

(31) Lu, W.; Liu, Y.; Gao, Y.; Geng, Q.; Gurbani, D.; Li, L.; Ficarro, S. B.; Meyer, C. J.; Sinha, D.; You, I.; Tse, J.; He, Z.; Ji, W.; Che, J.; Kim, A. Y.; Yu, T.; Wen, K.; Anderson, K. C.; Marto, J. A.; Westover, K. D.; Zhang, T.; Gray, N. S. Development of a Covalent Inhibitor of C-Jun N-Terminal Protein Kinase (JNK) 2/3 with Selectivity over JNK1. J. Med. Chem. 2023, 66 (5), 3356–3371. 10.1021/acs.jmedchem.2c01834.

(32) Bálint, D.; Póti, Á. L.; Alexa, A.; Sok, P.; Albert, K.; Torda, L.; Földesi-Nagy, D.; Csókás, D.; Turczel, G.; Imre, T.; Szarka, E.; Fekete, F.; Bento, I.; Bojtár, M.; Palkó, R.; Szabó, P.; Monostory, K.; Pápai, I.; Soós, T.; Reményi, A. Reversible Covalent C-Jun N-Terminal Kinase Inhibitors Targeting a Specific Cysteine by Precision-Guided Michael-Acceptor Warheads. Nat. Commun. 2024, 15 (1), 8606. 10.1038/s41467-024-52573-2.

(33) Geiger, T.; Wehner, A.; Schaab, C.; Cox, J.; Mann, M. Comparative Proteomic Analysis of Eleven Common Cell Lines Reveals Ubiquitous but Varying Expression of Most Proteins. Mol. Cell. Proteomics 2012, 11 (3), M111.014050. 10.1074/mcp.M111.014050.

(34) Xue, G.; Xie, J.; Hinterndorfer, M.; Cigler, M.; Dötsch, L.; Imrichova, H.; Lampe, P.; Cheng, X.; Adariani, S. R.; Winter, G. E.; Waldmann, H. Discovery of a Drug-like, Natural Product-Inspired DCAF11 Ligand Chemotype. Nat. Commun. 2023, 14 (1), 7908. 10.1038/s41467-023-43657-6.

(35) Tin, G.; Cigler, M.; Hinterndorfer, M.; Dong, K. D.; Imrichova, H.; Gygi, S. P.; Winter, G. E. Discovery of a DCAF11-Dependent Cyanoacrylamide-Containing Covalent Degrader of BET-Proteins. Bioorg. Med. Chem. Lett. 2024, 107, 129779. 10.1016/j.bmcl.2024.129779.

(36) Zhang, H.; Xu, D.; Huang, H.; Jiang, H.; Hu, L.; Liu, L.; Sun, G.; Gao, J.; Li, Y.; Xia, C.; Chen, S.; Zhou, H.; Kong, X.; Wang, M.; Luo, C. Discovery of a Covalent Inhibitor Selectively Targeting the Autophosphorylation Site of C-Src Kinase. ACS Chem. Biol. 2024, 19 (4), 999–1010. 10.1021/acschembio.4c00048.

(37) Salah, E.; Chatterjee, D.; Beltrami, A.; Tumber, A.; Preuss, F.; Canning, P.; Chaikuad, A.; Knaus, P.; Knapp, S.; Bullock, A. N.; Mathea, S. Lessons from LIMK1 Enzymology and Their Impact on Inhibitor Design. Biochem. J. 2019, 476 (21), 3197–3209. 10.1042/BCJ20190517.

(38) Niesen, F. H.; Berglund, H.; Vedadi, M. The Use of Differential Scanning Fluorimetry to Detect Ligand Interactions That Promote Protein Stability. Nat. Protoc. 2007, 2 (9), 2212–2221. 10.1038/nprot.2007.321.

(39) Krämer, A.; Kurz, C. G.; Berger, B.-T.; Celik, I. E.; Tjaden, A.; Greco, F. A.; Knapp, S.; Hanke, T. Optimization of Pyrazolo[1,5-a]Pyrimidines Lead to the Identification of a Highly Selective Casein Kinase 2 Inhibitor. Eur. J. Med. Chem. 2020, 208, 112770. 10.1016/j.ejmech.2020.112770.

(40) Fedorov, O.; Niesen, F. H.; Knapp, S. Kinase Inhibitor Selectivity Profiling Using Differential Scanning Fluorimetry. In Kinase Inhibitors; Kuster, B., Ed.; Methods in Molecular Biology; Humana Press: Totowa, NJ, 2012; Vol. 795, pp 109–118. 10.1007/978-1-61779-337-0_7.

(41) Vasta, J. D.; Corona, C. R.; Wilkinson, J.; Zimprich, C. A.; Hartnett, J. R.; Ingold, M. R.; Zimmerman, K.; Machleidt, T.; Kirkland, T. A.; Huwiler, K. G.; Ohana, R. F.; Slater, M.; Otto, P.; Cong, M.; Wells, C. I.; Berger, B.-T.; Hanke, T.; Glas, C.; Ding, K.; Drewry, D. H.; Huber, K. V. M.; Willson, T. M.; Knapp, S.; Müller, S.; Meisenheimer, P. L.; Fan, F.; Wood, K. V.; Robers, M. B. Quantitative, Wide-Spectrum Kinase Profiling in Live Cells for Assessing the Effect of Cellular ATP on Target Engagement. Cell Chem. Biol. 2018, 25 (2), 206–214.e11. 10.1016/j.chembiol.2017.10.010.

(42) Flanagan, M. E.; Abramite, J. A.; Anderson, D. P.; Aulabaugh, A.; Dahal, U. P.; Gilbert, A. M.; Li, C.; Montgomery, J.; Oppenheimer, S. R.; Ryder, T.; Schuff, B. P.; Uccello, D. P.; Walker, G. S.; Wu, Y.; Brown, M. F.; Chen, J. M.; Hayward, M. M.; Noe, M. C.; Obach, R. S.; Philippe, L.; Shanmugasundaram, V.; Shapiro, M. J.; Starr, J.; Stroh, J.; Che, Y. Chemical and Computational Methods for the Characterization of Covalent Reactive Groups for the Prospective Design of Irreversible Inhibitors. J. Med. Chem. 2014, 57 (23), 10072–10079. 10.1021/jm501412a.

(43) Böhme, A.; Thaens, D.; Paschke, A.; Schüürmann, G. Kinetic Glutathione Chemoassay to Quantify Thiol Reactivity of Organic Electrophiles--Application to Alpha,Beta-Unsaturated Ketones, Acrylates, and Propiolates. Chem. Res. Toxicol. 2009, 22 (4), 742–750. 10.1021/tx800492x.

(44) Quast, J.-P.; Schuster, D.; Picotti, P. Protti: An R Package for Comprehensive Data Analysis of Peptide- and Protein-Centric Bottom-up Proteomics Data. Bioinforma. Adv. 2022, 2 (1), vbab041. 10.1093/bioadv/vbab041.

(45) Rafiee, M.; Sigismondo, G.; Kalxdorf, M.; Förster, L.; Brügger, B.; Béthune, J.; Krijgsveld, J. Protease-resistant Streptavidin for Interaction Proteomics. Mol. Syst. Biol. 2020, 16 (5), e9370. 10.15252/msb.20199370.

